# GPatch enables chromosome-scale, gap-free pseudoassemblies from fragmented draft genomes

**DOI:** 10.1101/2025.05.22.655567

**Authors:** Adam Diehl, Alan Boyle

## Abstract

Recent advancements in sequencing technologies have yielded numerous long-read draft genomes, promising to enhance understanding of genomic variation. However, draft genomes are typically highly fragmented, posing challenges for functional genomics. We introduce GPatch, a tool that constructs chromosome-scale pseudoassemblies from fragmented drafts using alignments to a reference assembly. GPatch produces complete, accurate, gap-free assemblies preserving over 95% of nucleotides from human and non-human draft genomes. We show that GPatch pseudoassemblies can be used to construct Hi-C matrices, whereas fragmented draft assemblies cannot. Until complete genome assembly becomes routine, GPatch presents a necessary tool for maximizing the utility of draft genomes.

**GRAPHICAL ABSTRACT:** 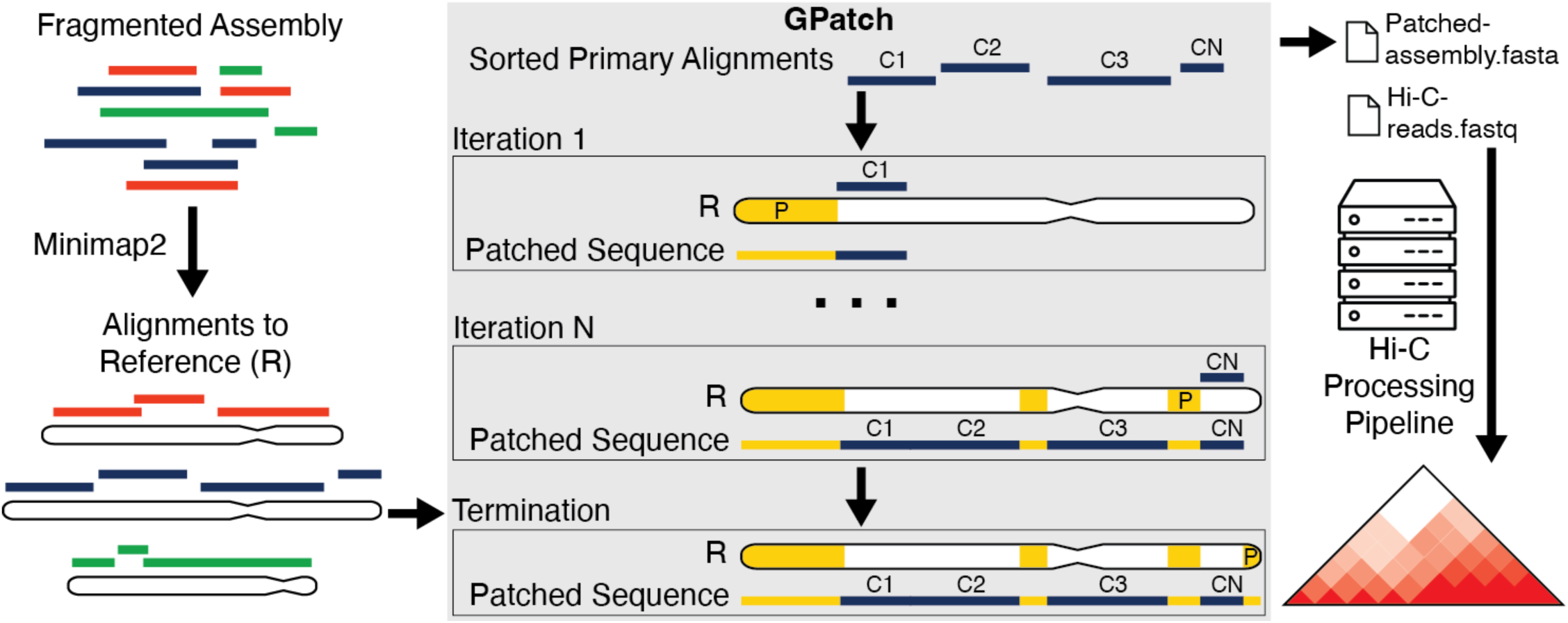

## INTRODUCTION

Since the publication of the second human genome, scientists have strived to understand the functional consequences of genomic variation. In ensuing years, numerous advances in sequencing technology and accompanying development of population, functional, comparative, and quantitative genomics methods have allowed researchers to generate an abundance of data in service of that goal [1–5]. These methods have enabled annotation of genome sequences with various types of functional evidence and efficient identification of variant loci, allowing insights into genome function and the consequences of variation. As a result, our understanding of how genomes function and how sequence variation within and between species produces differences in observable phenotypes has flourished. Still, important gaps in our understanding remain. This can be partially attributed to one inherent limitation of current genomics workflows: reliance on mapping to a reference genome assembly as the first step.

Reference genome assemblies represent the complete sequence of a species, serving as a scaffold for functional annotations, such as genes, and facilitating position-based comparisons both between individuals of the same or different species. Although reference assemblies may be based on a single individual or a sampling of multiple individuals [6], they invariably consist of a single sequence meant to represent the entire species. Unfortunately, as single sequences, reference genomes cannot adequately represent variation within a species [6], are often laden with rare alleles, and are missing numerous structural variants (SVs), some of which represent major alleles in the population [7–11]. These rare and missing variants, SVs especially, directly interfere with read mapping [7,12,13], leading to biases and misleading results in genomics methods including genotyping[12] and allele-specific RNA-seq[13]. Indeed, when missing variants from individual genomes are included in the reference genome, mapping rates and quality improve, with one study demonstrating recovery of 2.6% of unmapped reads, and increased quality scores for 25.7% of reads mapping to SVs after insertion of missing SVs into the GRCh38 reference genome [7].

To avoid these confounding effects, a complete, Telomere-to-Telomere (T2T) personal genome matched to the donor cell line would ideally be used to map reads in place of a reference assembly in genomics analyses. In this case, the genome assembly used for mapping will include matching alleles for all variants within the sequencing data, thus leading to improvements in mapping rates and quality. These improvements would be expected, in turn, to lead to improvements in sensitivity and specificity in downstream assays. Notably, advances in graph genomes and related software tools now make it possible to use personal genome assemblies in population and comparative genomics studies [14,15], eliminating the need for a single reference genome against which to map and compare results. However, very few functional genomics methods have been adapted to use graph genome data, relying on alignments to a single genome. Most of these methods can, however, utilize personal genomes in place of a reference assembly. Unfortunately, availability of suitable personal genome assemblies remains limited, and the cost and resource requirements to generate reference-grade personal genome assemblies remains high, particularly for mammalian-sized genomes. Indeed, in human, such efforts have been mostly confined to large, consortium-based projects [8,14,16–18] and, thus far, these efforts have produced only two truly chromosome-scale genomes: T2T-CHM13 [18] and T2T-HG002 [18,19]. These efforts were enabled by recent advances in long-read sequencing methods, such as Pacific Biosciences (PacBio) HiFi and Oxford-Nanopore (ONT) sequencing.

Long read alignments have the advantage of being able to span many repetitive regions of the genome, which are problematic to assemble when their length exceeds the read length [20–23], thus enabling gaps to be closed between contigs flanking a repeat locus. With current long read sequencing methods, it is cost-effective to routinely achieve sequence fragments exceeding the length of most genomic repeat regions at sufficient sequencing depths for de-novo genome assembly. However, to date, assembling these fragments into truly T2T genomes has required a combination of deep sequencing and additional data types to assemble the entire genome into complete chromosomes. T2T-HG002, for example, achieved this milestone using a combination of HiFi data at 120X depth and ONT data at 710X [14,19], while, for T2T CHM13, 30X HiFi and 120X ONT sequencing data, in combination with additional strand-seq, Hi-C, and BioNano optical map data were required [18]. Such resources are beyond the reach of most individual labs. Another group was recently able to fully-assemble 63.3% of chromosomes across 28 diploid genomes [16] using more attainable sequencing depths of 45.7X for HiFi and 60.5X for ONT. These assemblies are not T2T, with an average assembly containing only 14 complete chromosomes and a variable number of unassigned contigs per phased, haploid genome. Meanwhile, several large consortia, including groups from the 1000 Genomes Project (1KG) [17], Human Genome Structural Variation Consortium (HGSVC) [8], and Human Pangenome Reference Consortium (HPRC) [24], have collectively produced hundreds of publicly-available draft personal genomes, derived primarily from immortalized lymphoblastoid cell lines (LCLs), using various combinations of sequencing methods and depths. Unfortunately, these genomes exist in varying states of completion, as sets of dozens to thousands of unassigned contigs, making them unsuitable for direct replacement of reference genomes in most functional genomics applications. While continued development of modern assemblers has yielded improvements in the ability to yield chromosome-scale assemblies with fewer data requirements [25–30], it is unlikely that these methods will be applied to all existing long-read assemblies in the immediate future, thus fragmentation is likely to remain a problem for the foreseeable future. Furthermore, we are unaware of any current efforts to systematically produce long-read genome assemblies for commonly-used non-LCL cell lines, posing a problem for numerous researchers broadly focused on functional, developmental, and disease biology.

The fragmented nature of currently-available long-read personal genomes presents a problem for many functional genomics assays, particularly those relying on proximity within the genome. The most-notable of these involve chromatin conformation capture, such as Hi-C [31] and ChIA-PET [32]. Indeed, Hi-C processing pipelines are not designed for fragmented genomes, relying on contiguity across entire chromosomes to detect cis-interactions between distant loci. In this case, genome fragmentation hinders detection of features artificially split across contig boundaries, inflates the number of intrachromosomal (trans) contacts, and can prevent construction of contact matrices from raw data. Fragmentation also poses challenges in annotating features of interest within individual assemblies and evaluating their spatial relationships, for example, linking distal enhancer elements to their target genes [33]. Furthermore, since many genomics operations, such as Hi-C matrix construction and normalization, scale in resource requirements proportionally to the number of reference contigs, certain analyses are intractable using a fragmented draft genome. Thus, methods for assembling fragmented draft genomes into chromosome-scale pseudoassemblies are necessary. There are two common approaches for doing so: reference-free, and reference-guided methods.

Reference-free scaffolding methods often use mappings to supplementary datasets, such as optical, physical, or linkage maps [34,35], or long-range interaction data, such as Hi-C [36,37] or linked reads [38], to arrange contigs into single molecules. These methods rely on overlaps between contigs and the map(s) used, or the observed contact frequency between loci to order and orient contigs. Because these mappings are not uniformly distributed throughout the genome, specifically being depleted in low-complexity and repetitive regions, these methods often fail to produce chromosome-scale scaffolds. Since resulting scaffolds often contain gaps and discontinuities, the resulting genomes cannot be considered complete. For example, when applied to a contig-level assembly of the NA12878 genome containing 2,886 contigs [39], SALSA2, a popular Hi-C scaffolding tool, assembled only seven chromosomes to near-chromosome-scale, with the P and Q arms of most chromosomes split into separate scaffolds [37]. Furthermore, the mappings used to establish order and orientation often lack information on the length of any implied gaps between scaffolded contigs, nor can they be used to impute the sequence content between them. Finally, these methods all require additional assays and data processing separate from the initial sequencing step, which can substantially inflate the cost of an analysis.

By contrast, reference-guided approaches utilize sequence alignments to a complete reference assembly rather than external data to assemble contigs into chromosome-scale scaffolds. Reference-guided scaffolding assembles contigs into pseudomolecules by arranging them along chromosomes based on their aligned positions, with intervening gaps filled with N characters. This has the distinct advantage of harnessing the contiguity of the reference assembly to scaffold contigs even across long repetitive regions, such as centromeres, which are problematic for reference-free methods. A handful of methods exist for reference-based scaffolding [40–47]. However, since these methods all leave N-gaps in scaffolded sequences and often do not impute gap length, their ability to serve as drop-in substitutes for a reference genome is limited. Reference-guided patching goes a step further by filling N-gaps with sequence imputed from the corresponding loci in the reference genome, thus yielding contiguous, gap-free, chromosome-scale pseudoassemblies which can, theoretically, be used interchangeably with a reference assembly. We are aware of only one publicly-available tool for reference-based patching: RagTag Patch [48], an extension of RaGOO [49]. However, RagTag patch is presented as a beta release that has not been rigorously tested or presented in the published literature, and has not been substantively updated since October 2021.

Here we present GPatch, a utility that presents a straightforward, intuitive, and flexible approach to reference-guided patching. We show that GPatch is able to reproducibly assemble full-length, chromosome-scale pseudoassemblies from both human and non-human long-read assemblies with varying degrees of fragmentation, producing pseudochromosomes that closely approximate the structure of their target chromosomes. In particular, we demonstrate that GPatch faithfully produces gap-free, chromosome-scale pseudoassemblies given only a draft genome and a reference assembly, something that RagTag Patch could not replicate. We show that GPatch pseudoassemblies compare favorably with reference assemblies in terms of contiguity, Hi-C read mapping rates, and alignment quality, while incorporating over 95% of nucleotides from the draft assembly. Notably, we were able to construct and normalize Hi-C matrices and call loops using a GPatch pseudoassembly, whereas initial matrix construction failed using the unpatched source assembly. From these matrices, we were able to recover loop predictions spanning contig boundaries in the draft genome, and those that are anchored by sequences not appearing in T2T-CHM13, demonstrating the advantages of using the GPatch pseudoassembly over either the source draft assembly or the T2T-CHM13 reference assembly. We show that GPatch performs equally well on human and non-human draft assemblies, demonstrating high fidelity between a patched pseudoassembly built from tomato M82 contigs [49], with both the target M82 assembly and the Heinz 1706 reference assembly [50]. Thus, GPatch is applicable to any species or cell type for which a draft personal genome and reference assembly are available. We conclude that GPatch is a valuable tool to facilitate the use of long-read personal genomes in everyday computational analyses, particularly where genomic fragmentation would hinder data analysis and interpretation.

## MATERIALS AND METHODS

### Data Simulation

We initially selected two genomes from which to model the contig-length distribution: NA12878 from the Human Genome Structural Variation Consortium (HGSVC NA1287) [8], and HG002 from the Human Pangenome Reference Consortium [14] (Supplementary Table S1: Data Sources). Genomes were downloaded in FASTA format and the lengths of all contigs therein were extracted using awk and stored in text files. These were then supplied to a custom Python script, along with the T2T-CHM13 genome assembly, to break each T2T-CHM13 chromosome into a set of pseudocontigs. Briefly, we loop over reference chromosomes, initially setting a position tracker *(p)* to zero to mark the start of a chromosome. Next, we draw a length *(l)* from the contig-length distribution of the model assembly. We then extract a fragment of length *l* from the current chromosome sequence and store it as a FASTA record in the output file. Finally, we set *p* equal to the end position of the extracted sequence fragment. We continue drawing values of *l* and extracting pseudocontigs of corresponding lengths until we reach the end of the chromosome. This process is then repeated until all reference chromosomes are exhausted. The end result is a pseudoassembly consisting of contigs closely matching the length distribution from the model assembly. In addition to the FASTA output, the script also produces a BED file documenting the coordinates of all pseudocontigs within the reference genome.

### Indel Simulation with SURVIVOR

Starting with the NA12878 and HG002 pseudoassemblies described in the *Data Simulation* section, we produced pseudoassemblies containing 5,000-10,000 random indels and single-nucleotide variants at a 1% rate using the SURVIVOR software package [53]. As contigs with lengths less than a fixed 10kb threshold are silently dropped from SURVIVOR output, we had to post-process the SURVIVOR FASTA output to add-back a small number of short contigs in each assembly using a custom script. Since indels within the SURVIVOR output genome change the coordinate system of the pseudoassembly, we could not use the unmodified reference genome, nor pseudocontig BED coordinates based on the T2T-CHM13 reference, as the target genome for comparison. Accordingly, we produced complete target genomes from the SURVIVOR-mutated assemblies by concatenating contigs into chromosomes based on their known order in T2T-CHM13. BED coordinates for each contig in the frame of the target genome were stored to facilitate position-based comparisons between the patched and target genomes.

### Patching Simulated Data with GPatch

BASH shell scripts were used to automate alignment and patching of the simulated NA12878 and HG002 genomes, along with realignment to the target genome and dot-plot construction with R, as described in *Dot Plot Construction* below. Pseudoassemblies were first aligned to the T2T-CHM13 reference genome [18] with minimap2 [51] using parameters: *‘minimap2 <REFERENCE_GENOME> <PSEUDOASSEMBLY> -x asm20 -t 24 -à*. The *‘-x asm20*’ argument was used to increase the aligner’s tolerance for large gaps, while the -a argument toggles SAM output format and base-alignment within the minimap2 algorithm. Notably, using *‘-x asm5’*, the default for RagTag patch, resulted in fewer placed contigs and a relative increase in patch nucleotides in optimization trials (data not shown). SAM output was converted to BAM format on-the-fly by piping minimap2 output into SAMtools view [62]. The resulting BAM and the T2T-CHM13 reference genome were then supplied to GPatch, with parameters ’*GPatch.py -q <$prefix.pseudocontigs.bam> -r <REFERENCE_GENOME> -x <PREFIX> -d -m 10 -t*’ where -d causes GPatch to drop reference contigs without alignments from the output, -m 10 sets the mapping quality threshold for contig alignments to 10, and -t indicates that overlapping contig alignments should not be 5’ trimmed. Mapq 10 was chosen as this is the default for RagTag patch. Please note that the -t option has been removed in versions of GPatch beyond 0.3.6, wherein contig trimming is disabled as no sequence-level overlaps between distinct contigs should be present in properly-formed genome assemblies. Statistics on patched assembly length, contig content, genome completeness, and accuracy were then assembled into text files using a combination of awk and custom Python scripts.

### GPatch Performance vs. Mapping Quality Threshold

In order to determine the effect of mapping quality (mapq) threshold on GPatch performance, we repeated patching steps outlined in *Patching Simulated Data with GPatch*, at mapq thresholds 0, 10, 20, 30, and 40, for the NA12878 SURVIVOR simulated dataset. At each mapq threshold, we compiled statistics on patched assembly length, contig content, genome completeness, and accuracy into text files using a combination of awk and custom Python scripts, and prepared bar plots for Supplementary Figure S8 with a custom R script. To compile counts presented in Table S4, samtools was used to extract primary alignments for all unplaced contigs from the bam file for the NA12878 SURVIVOR alignment to CHM13. SAM flags and quality scores were examined to determine the number of contigs dropped due to: 1) no mapping, 2) failure to pass the mapq threshold, and 3) nested mapping.

### Patching Simulated Data with RagTag Patch

BASH shell scripts were used to automate processing of simulated NA12878 and HG002 genomes with RagTag Patch, along with realignment for dot-plot production in R. After extensive optimization with the goal of maximizing contig recall in the results, we ran RagTag Patch with the following parameters: *‘ragtag.py patch --aligner minimap2 --mm2-params “-x asm20 -c -t 24” -o $OUT_DIR -f 100 -s 10000 <PSEUDOASSEMBLY> <REFERENCE_GENOME>’.* As part of its preprocessing steps, RagTag patch renames all sequences from the query and target genomes, ostensibly to avoid naming collisions in the patching process. RagTag Patch also includes all unscaffolded contigs in the output, along with any patched results. These features made it difficult to evaluate RagTag Patch results and compare them to GPatch. To ease interpretation and comparisons with GPatch results, we utilized a custom script to filter unscaffolded contigs from RagTag Patch fasta output and rename scaffolds according to their corresponding reference chromosomes. We used a second script to process the agp-formatted data on contig-placement within scaffolds, converting it into a BED format compatible with GPatch output. Statistics on patched assembly length, contig content, genome completeness, and accuracy were then assembled into text files using the same set of awk commands and scripts used for the GPatch results.

### Patching Simulated Data with RagTag Scaffold

BASH shell scripts were used to automate processing of simulated NA12878 and HG002 genomes with RagTag Scaffold. We used the following parameters: *‘ragtag.py scaffold --aligner minimap2 --mm2-params “-x asm20 -c -t 24” -o $OUT_DIR -r <REFERENCE_GENOME> <PSEUDOASSEMBLY>’.* We used awk to convert the agp-formatted data into a BED format. Statistics on patched assembly length, contig content, genome completeness, and accuracy were then assembled into text files using the same set of awk commands and scripts used for the GPatch results.

### Dot Plot Construction

For all patched genomes, we produced dot-plots in comparison to the corresponding target genome using the ‘pafr’ R package (https://dwinter.github.io/pafr/). We first aligned the patched genome to its respective target genome with minimap2 [51], with parameters *‘minimap2 -x asm20 -t 24 <REFERENCE_FASTA> <PATCHED_FASTA>’.* PAF-formatted output was read into R with pafr’s ‘read_paf’ function, after which we used pafr’s dotplot function to produce dotplots, in PDF format, for all autosomes and the X and Y chromosomes.

### Ideogram Construction

We utilized the PhenoGram web application (https://visualization.ritchielab.org/phenograms/plot) to produce ideograms illustrating the locations of patches within GPatch results. We first used awk to reconfigure the patch coordinates from GPatch’s patches.bed into the text format used by PhenoGram. This was supplied as the input file for PhenoGram. We supplied the chrom.sizes file for T2T_CHM13, augmented with telomere locations obtained from the UCSC Table Browser [63], as the genome. We additionally selected “Standard Algorithm” for “Phenotype Spacing”, and checked the “Chromosome only” box to toggle printing only the ideogram without accompanying annotations. Resulting images were stored in PNG format and further processed locally to desaturate and improve contrast.

### Patching Biological Data with GPatch

BASH shell scripts were used to automate the GPatch patching process for the HGSVC NA12878 and HPRC HG002 genomes. Initial alignment of each genome to the T2T-CHM13 reference genome was performed with minimap2 using parameters *‘minimap2 <REFERENCE_GENOME> <CONTIG_ASSEMBLY> -x asm20 -t 24 -à*, with output piped into SAMtools view to convert SAM output to BAM. Resulting BAM files were processed with GPatch using parameters ‘*GPatch.py -q <$prefix.contigs.bam> -r <REFERENCE_GENOME> -x <PREFIX> -d -m 10 -t’* where -d causes GPatch to drop reference contigs without alignments from the output, -m 10 sets the mapping quality threshold for contig alignments to 10, and -t indicates that overlapping contig alignments should not be 5’ trimmed. Mapq 10 was used for consistency with the analysis of simulated data. Please note that the -t option has been removed in versions of GPatch beyond 0.3.6, wherein contig trimming is disabled as no sequence-level overlaps between distinct contigs should be present in properly-formed genome assemblies. Statistics on patched assembly length, contig content, and genome completeness were then assembled into text files using a combination of awk and custom python scripts. Finally, dot plots were prepared according to steps outlined in the *Dot Plot Construction* methods section to compare the initial patched genome to the T2T-CHM13 reference, and to matched target genomes, T2T-NA12878 [16] and T2T-HG002 [52].

### Patching the Tomato M82 Assembly

A BASH shell script was used to automate the GPatch patching process for the tomato M82 assembly [49]. Contigs were first recovered from chromosome-scale scaffolds in the M82 assembly by splitting sequences at stretches of 20 or more ‘N’ characters using a custom python script. Initial alignment of recovered contigs to the SL3 reference assembly [50] was performed with minimap2 using parameters *‘minimap2 <REFERENCE_GENOME> <CONTIG_ASSEMBLY> -x asm20 -t 24 -à*, with output piped into SAMtools view to convert SAM output to BAM. Resulting BAM files were processed with GPatch using parameters ‘*GPatch.py -q <$prefix.contigs.bam> -r <REFERENCE_GENOME> -x <PREFIX> -d -m 10 -t’* where -d causes GPatch to drop reference contigs without alignments from the output, -m 10 sets the mapping quality threshold for contig alignments to 10, and -t indicates that overlapping contig alignments should not be 5’ trimmed. Mapq 10 was used for consistency with the analysis of simulated data. Please note that the -t option has been removed in versions of GPatch beyond 0.3.6, wherein contig trimming is disabled as no sequence-level overlaps between distinct contigs should be present in properly-formed genome assemblies. Statistics on patched assembly length, contig content, and genome completeness were then assembled into text files using a combination of awk and custom python scripts. Finally, dot plots were prepared according to steps outlined in the *Dot Plot Construction* methods section to compare the initial patched genome to the SL3 reference, and to the unfragmented M82 assembly.

### Automated Contig-Breaking

PAF alignments generated during dot-plot preparation in *Patching Biological Data with GPatch* were further processed with a custom python script, included in the GPatch GitHub repository, to identify large-scale rearrangements within the patched genome that represent likely misjoins in the draft assembly. These misjoins can be between different regions of the same chromosome or between different chromosomes, and may represent any class of structural variant. These were identified as alignments or clusters of partial-alignments, within a maximum specified distance of each other that are inverted, translocated, or duplicated in the patched genome relative to the reference genome. To locate these, we looped over contigs in the input assembly, first retrieving all overlapping partial alignments based on their coordinates in the patched genome. Partial alignments were then clustered based on their chromosome, strand, and mapped position (i.e., position in the reference/target genome) such that partial-alignments within a maximum distance from each other, on the same chromosome and strand (including overlapping and nested alignments), are merged into a single interval. Merged intervals were then filtered to identify clusters representing inversions, duplications, and translocations: i.e., those that map to the reverse strand, or whose mapped position in T2T-CHM13 has shifted by a user-configurable minimum distance threshold of 1MB. Remaining intervals were then filtered to retain only those exceeding a user-configurable size threshold of 1MB. Breakpoint loci within the patched genome were then supplied to a second Python script to identify corresponding contigs from the source assembly and break them at the inferred positions, with output written in fasta format. These assemblies were then reprocessed with GPatch following the same steps described in *Patching Biological Data with GPatch.* Dot-plots comparing GPatch results following contig breaking to T2T-CHM13 and corresponding target genomes were prepared as described in *Dot Plot Construction*.

### Hi-C Data Mapping and Processing

Hi-C read data for NA12878, in FASTQ format, were obtained from the Sequence Read Archive (SRA) repository for Rao and Huntley, 2014 [31] (Supplementary Table S1). Given the extraordinary amount of sequence data for NA12878 generated in this study, we selected only the two largest FASTA files for processing (Supplementary Table S1), totalling approximately 1.75 billion reads. We utilized a custom Hi-C analysis pipeline based on the 4DN Hi-C Processing pipeline [55]. Briefly, FastQC was used to verify the quality of Hi-C read data prior to mapping to a given reference genome with BWA-MEM [64]. Mapped reads with quality scores 40 and above were extracted from the BAM output, converted into pairs format, sorted, and replicates merged, using a combination of pairtools [65] commands. Pairtools was further employed to mark and exclude duplicate pairs, and select only “UU”, “UR”, and “RU” contact pairs for further processing. Restriction enzyme fragment data were then superimposed on pairs data using the *fragment_4dnpairs.pl* script from the 4DN consortium. Finally, we used Juicer [56] to construct Hi-C matrices from the final pairs file and apply normalization using the KR method [31]. All analysis steps were automated with Makefiles to ensure reproducibility, and were performed on a local compute cluster using memory-optimized nodes allocated with 500GB RAM, 12 3.0 GHz Intel Xeon Gold 6154 compute cores, and 4TB HDD. Hi-C loop calling was performed using the GPU-enabled version of hiccups, on a local server equipped with an NVIDIA Titan-V GPU and 270GB available RAM, using the Juicer Tools juicer_postprocessing.sh script [56]. Loop calling was performed with hiccups [56] at 10kb and 5kb resolutions for Hi-C data mapped to both the T2T-CHM13 and GPatch NA12878 assemblies. Due to unexplained crashes of the hiccups software, loop calling could only be completed for resolutions 5kb and 10kb.

### Hi-C Visualization

Normalized matrices in .hic format were browsed locally in Juicebox for initial comparison across stored resolutions before identifying 80kb as the optimal resolution for visualization. Hi-C heat map plotting for individual chromosomes was then automated with a custom Python script utilizing the CoolBox API [66]. Plots were generated at 80kb resolution with balanced normalization, for both T2T-CHM13 and GPatch-NA12878 based Hi-C matrices, with plots stored in SVG format. SVG images for each dataset were then converted to PDF format using the online tool: https://tools.pdf24.org/en/svg-to-pdf [67], combined into a single PDF file in Adobe Acrobat, and reduced to sets of six thumbnails per standard letter sized page (landscape format) using the MacOS Preview tool. Thumbnails were extracted from the resulting PDF files, edited, and arranged into final figures in Adobe Illustrator, with the following edits applied uniformly to each image: 1) Bilateral scaling to 120%; 2) Adjust embedded heatmap and scale images to improve contrast using the “levels” tool from the Phantasm plugin (https://astutegraphics.com/plugins/phantasm), with input levels for the RGB channel set to 50:0.5:255, and output levels set to 0:240. Final figures were saved in PDF format.

### Hi-C Loop Comparison

In order to compare the positions of individual loops and loop anchors between T2T-CHM13 and GPatch NA12878, we first prepared liftover chains using the nf-lo Nextflow pipeline [68], following steps described at https://genome.ucsc.edu/cgi-bin/hgTrackUi?hgsid=1390827233_st3mhvM8SnGwC9IK4FZ9ysMqCN54&db=hg38&c=chrX&g=chm13LiftOver, with T2T-CHM13 as the target and GPatch NA12878 as the source, and using minimap2 as the aligner, using the same mapping parameters as used previously in simulated and biological data analyses. Starting with loop predictions from the patched NA12878 assembly, loop predictions at 5kb and 10kb resolution were decomposed into individual loop anchor loci by splitting each line in the BEDPE source file into separate lines for the upstream and downstream loop anchors and assigning the BED name field for both with a matched ID number, with upstream and downstream anchors tagged with _1 and _2 suffixes, respectively. Resulting BED files were processed with the UCSC LiftOver tool [63] to convert from GPatch NA12878 frame to T2T-CHM13 frame. We first counted loops for which one or both anchors failed to liftover, indicating presence in sequence unique to the GPatch NA12878 assembly. Next, BedTools [57] was used to identify lifted GPatch NA12878 loop anchors that overlap T2T-CHM13 loop anchors, using the command *‘bedtools intersect -a <NA12878 anchors> -b <T2T-CHM13 anchors> -ù*. A custom python script was then used to reassemble intersecting anchors into complete or partial loop annotations in BEDPE format, with “missing” loop anchors, from loops where only one anchor intersects a T2T-CHM13 anchor, denoted by ‘.’ characters in the affected BEDPE fields. From this, we were able to count completely and partially shared loops between GPatch NA12878 and T2T-CHM13. Loops unique to GPatch NA12878 were identified based on records in the “unmapped” fraction of the LiftOver results.

To determine which loops have anchors that are separated by at least one contig or patch boundary in GPatch NA18278, we decomposed loops into loop intervals, defined as the maximal interval between the upstream and downstream loop anchor coordinates, and stored these in BED format. Likewise, the single-base coordinates of 5’ and 3’ contig boundaries, in the frame of the GPatch NA12878 assembly, were extracted from the corresponding contigs.bed file generated within the patching step. We then used bedtools intersect, with options ‘-a <loop_intevals.bed> -b <contig_boundaries.bed> -wa -ù to reduce intersections between loop intervals and contigs to a list of unique loop intervals that overlap at least one contig boundary. This was repeated for loop calling resolutions 10kb and 5kb. Average loop lengths for 5kb and 10kb resolutions were calculated in awk by totaling the lengths of all loop intervals from each resolution and dividing by the number of total loop predictions.

## RESULTS

### The GPatch Algorithm

GPatch (Fig. 1) is designed to simultaneously scaffold contigs based on alignments to a reference genome and fill intervening gaps with reference sequence to yield a chromosome-scale pseudoassembly. GPatch takes an alignment of contigs from a draft assembly to a reference genome assembly, such as T2T-CHM13, in BAM format (Fig. 1A). This file need not be sorted nor indexed, and can be produced using any aligner capable of producing BAM output including a CIGAR string, but we strongly recommend minimap2 [51]. Initially, GPatch isolates all primary alignments passing the chosen mapping quality threshold (default 30) from the input BAM. Multimapped contigs are incorporated based on their designated primary alignment. However, it is, in principle, possible to filter the input BAM to remove multimapped contigs prior to running GPatch if desired. Indeed, as long as the final alignment is in BAM format and includes CIGAR strings for all primary alignments, any processing pipeline may be used to refine input alignments for GPatch. Next, 5’ and 3’ breakpoints in the reference sequence are inferred for each alignment by padding start and end coordinates with the lengths of any 5’ or 3’ soft-clipped regions found in the alignment’s CIGAR string. Alignments are then sorted by position and, by default, filtered to remove nested alignments, which cannot be unambiguously ordered. Optionally, alignments can also be filtered to exclude alignments falling entirely within blacklist regions, supplied as a BED file. The final, filtered, position-sorted alignments (Fig. 1B) are then processed with the core GPatch algorithm (Fig. 1C-F).

**Figure 1:**
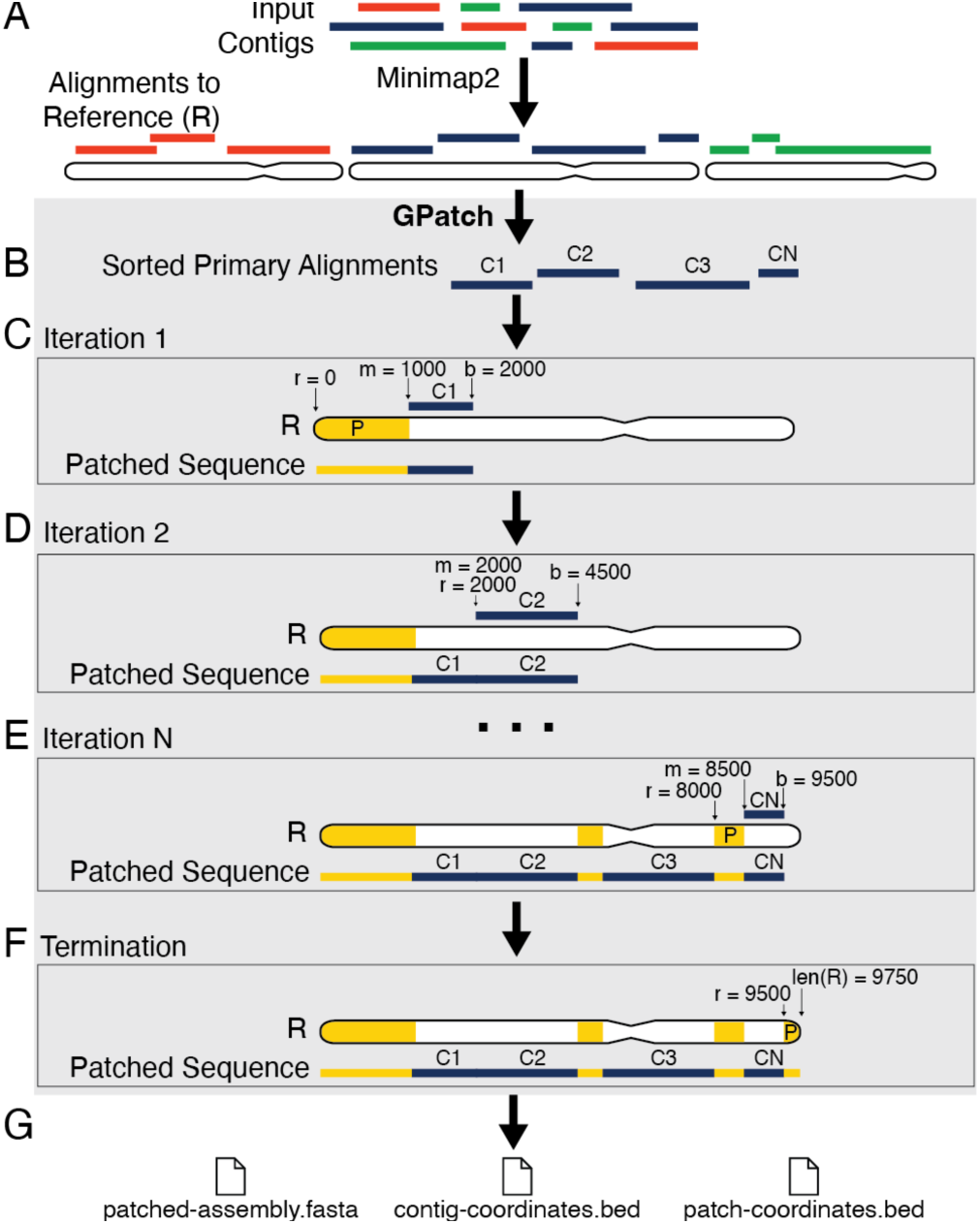
The GPatch Algorithm. A) Contigs are first aligned against the reference genome. GPatch takes as inputs alignments from (A), and a reference genome *R*. The grey box indicates the core GPatch recurrence, which is repeated for each reference chromosome. Primary alignments are first filtered to remove low-quality and nested alignments and sorted by position. B) Primary alignments to the current reference chromosome are retrieved and the patched sequence *S* is initialized to an empty string. C) Patching begins by drawing contig C1 from the sorted list, and setting *r*=0, while *m* and *b* are set to C1’s 5’ and 3’ reference breakpoints. If *m > r*, we append a patch, *P = R[r:m]* to *S*, followed by the entire contig sequence, including any soft-clipped regions. Finally, *r* is set equal to *b*. D) In iteration 2, we draw contig C2 from the sorted list and set *m* and *b* to its 5’ and 3’ alignment breakpoints. Since *m == r*, no patch is necessary: the contig sequence is appended directly to *S*. Finally, *r* is set equal to *b*. E) This process is repeated until all contigs have been incorporated into *S*. F) To terminate the patched sequence, *r* is compared to the length of the chromosome, *len(R)* and, if *r < len(R)*, a terminal patch is appended to *S*. Steps B through F are repeated until all reference chromosomes have been patched. G) GPatch produces three output files: patched-assembly.fasta = the patched genome, contig-coordinates.bed = contig boundaries in the coordinate frame of the patched genome, and patch-coordinates.bed = patch boundaries in the coordinate frame of the reference genome. Alt text: Flow chart describing the GPatch algorithm.

The core GPatch algorithm loops over chromosomes in the reference assembly. We first retrieve all alignments falling on a given chromosome from the final sorted BAM file (Fig. 1B). The patched chromosome sequence is initialized as an empty string and a tracker for the current position within the reference sequence, *r*, is set to zero. We then loop over contig alignments, alternately incorporating contigs and, as needed, patches, until all contigs have been incorporated into the patched sequence and the end of the reference chromosome is reached. For each sorted contig, we first compare the 5’ contig breakpoint, *m*, to *r*. If *m > r*, we add a patch spanning the reference interval from *r* to *m* to the chromosome sequence, followed by the complete, unmodified contig sequence (Fig. 1C). If *m* == *r*, contig mappings are bookended, thus no patch from the reference is necessary and the unmodified contig sequence is appended to the chromosome string (Fig. 1D). If *m < r*, there is an overlap between neighboring contig mappings. In these cases, contigs will be bookended. In versions through 0.3.6, the default behavior was to trim the 5’ end of the overlapping contig end, truncating the subsequence spanning the interval between *m* and *r*. However, since properly-constructed assemblies should not contain sequence-level overlaps between distinct contigs, trimming proved to be universally undesirable in preliminary tests (data not shown), thus the -t/–no_trim option was used in all presented analyses and trimming has been disabled in GPatch versions above 0.3.6. Contigs for which *m < r and b < r*, where *b* is the 3’ contig breakpoint, represent nested mappings, which cannot be unambiguously ordered, likely representing erroneous/repetitive alignments. These are dropped by default, but may be optionally bookended after the enclosing contig in GPatch versions above 0.3.6. This process is repeated until all contigs have been placed (Fig. 1E). To terminate the patched sequence, we compare *r* to the length of the reference chromosome and, if necessary, incorporate a final patch to reach the 3’ terminus of the reference chromosome (Fig. 1F). The patched chromosome record is then written to output in FASTA format, while coordinates of contigs and patches are written to their own files, in BED format (Fig. 1G). By checking whether the first and last contigs encompass the 5’ and 3’ termini of the reference chromosome and applying terminal patches as needed, we ensure that the algorithm always produces complete sequences with respect to target chromosome span. However, GPatch versions above 0.3.6 may be configured to leave off 5’ and 3’ patches, in which case the patched sequence will be terminated at both ends by contigs. Importantly, BED coordinates for all contigs and patches both document the exact composition of the patched genome and render the patching process fully-traceable. Excluding alignment time, this process can be completed in about 6-20 minutes for a typically-sized mammalian genome.

### Analysis of Simulated Data

To establish an upper-bound estimate for the performance and accuracy of GPatch, we generated four simulated draft genomes of varying difficulties. We selected two publicly available human genomes on which to model our simulated data: NA12878 from the HGSVC consortium [52], representing a highly-fragmented genome consisting of thousands of contigs, and HG002 from the HPRC consortium [24], representing a highly-contiguous genome consisting of hundreds of contigs (Supplementary Table S1). For each genome, the T2T-CHM13 reference genome was fragmented into a set of simulated contigs matching the length distribution of the model assembly randomly tiling each reference chromosome. This yielded the NA12878 and HG002 “no-indel” simulated draft assemblies, which contain no structural or nucleotide-level variation relative to the reference assembly (Supplementary Table 2). These datasets represent an idealized case where the reference assembly contains a perfect alignment match for every contig in the respective simulated draft assembly. To present a more realistic scenario, we used SURVIVOR [53] to randomly introduce 5,000-10,000 indels and single-nucleotide-variants at a 1% rate, starting with the no-indel simulated draft assemblies, which we call the SURVIVOR datasets (Supplementary Table 2). As a basis for comparison, all contigs from the SURVIVOR dataset were concatenated according to their known order in the T2T-CHM13 genome to produce simulated target assemblies for use in direct sequence comparisons. For simplicity, we apply the term “target assembly”, which is used as the baseline for comparison with patched pseudoassemblies, to either the complete T2T-CHM13 sequence, for the no-indel set, or to the concatenated SURVIVOR simulated draft genomes. All four datasets were processed with both GPatch and RagTag Patch to produce patched pseudoassemblies. For both methods, we noted that the completeness of the patched pseudoassembly relied heavily on the choice of aligner and alignment parameters used to create the initial alignment. Therefore, we used minimap2 as the aligner for both GPatch and RagTag patch and fixed alignment parameters across both methods. For each method, we separately optimized patching parameters to achieve an optimal tradeoff between maximizing contig capture and placement accuracy, using a mapping quality (mapq) threshold of 10 as this was the default used by the RagTag method.

We selected a set of metrics by which to compare patched pseudoassembly accuracy and completeness, using comparisons of contig order, orientation, and placement in the patched pseudoassembly to their known values in the target assembly, and direct sequence comparisons between the patched and target chromosome sequences. Assembly completeness was evaluated using the fraction of contigs and nucleotides from the simulated draft assembly that are placed in the patched pseudoassembly and the percentage of adjacent contig pairs from the target assembly recovered in the patched pseudoassembly. Ordering and orientation accuracy were evaluated in three ways. First, ordering edit distance was defined as the Levenshtein distance between the true and observed vectors of contig IDs along each chromosome from the patched and target assemblies. This measures departures from the true contig ordering, including the effect of dropped contigs. Second, pair recall is defined as the fraction of correct adjacent pairs from the target assembly recovered in the patched pseudoassembly, offering another view of contig-completeness including the effects of dropped contigs. Third, we define the ordering error rate as the fraction of contigs placed out-of-order in the patched pseudoassembly relative to the target assembly, calculated as half the Levenshtein distance between the observed and sorted contig ID vectors from each patched chromosome, offering a view of ordering accuracy that is not affected by dropped contigs. Finally, we used bag distance to approximate the edit distance between each patched chromosome and its counterpart in the target assembly. Dot plots were used to visualize any departures in collinearity between the patched pseudoassemblies and target assemblies.

GPatch performed exceptionally well on the no-indel dataset, placing 98.02% of NA12878 and 98.62% of HG002 simulated contigs, and capturing 99.87% and 99.99% of nucleotides from their respective simulated draft assemblies (Fig. 2A), with 100% of contigs placed with the correct chromosome, order, and orientation. We observed ordering edit distances less than 2% for both sets (Fig. 2B) and pair recall of 97.40% for NA12878 and 97.58% for HG002, respectively, indicating a low rate of dropped contigs in the patched genome. Remarkably, we were able to recover the exact sequence for all target chromosomes in the patched genome, as evidenced by identical N50, identical MD5sums, and an average normalized bag distance of 0 between patched and target genomes (Fig. 2C). Furthermore, GPatch achieved 100% pair-accuracy (Fig. 1E) and a 0% ordering error rate (Fig. 1F), placing all contigs in their correct relative order. This was clearly visible in dot-plots against T2T-CHM13, where we observed a near 1:1 correspondence between patched and target genomes (Supplementary Fig. S1-S2). Where patch sequences occurred, they were relatively short and tended to fall near/within telomeres and pericentromeric regions (Supplementary Fig. S3A-B).

**Figure 2:**
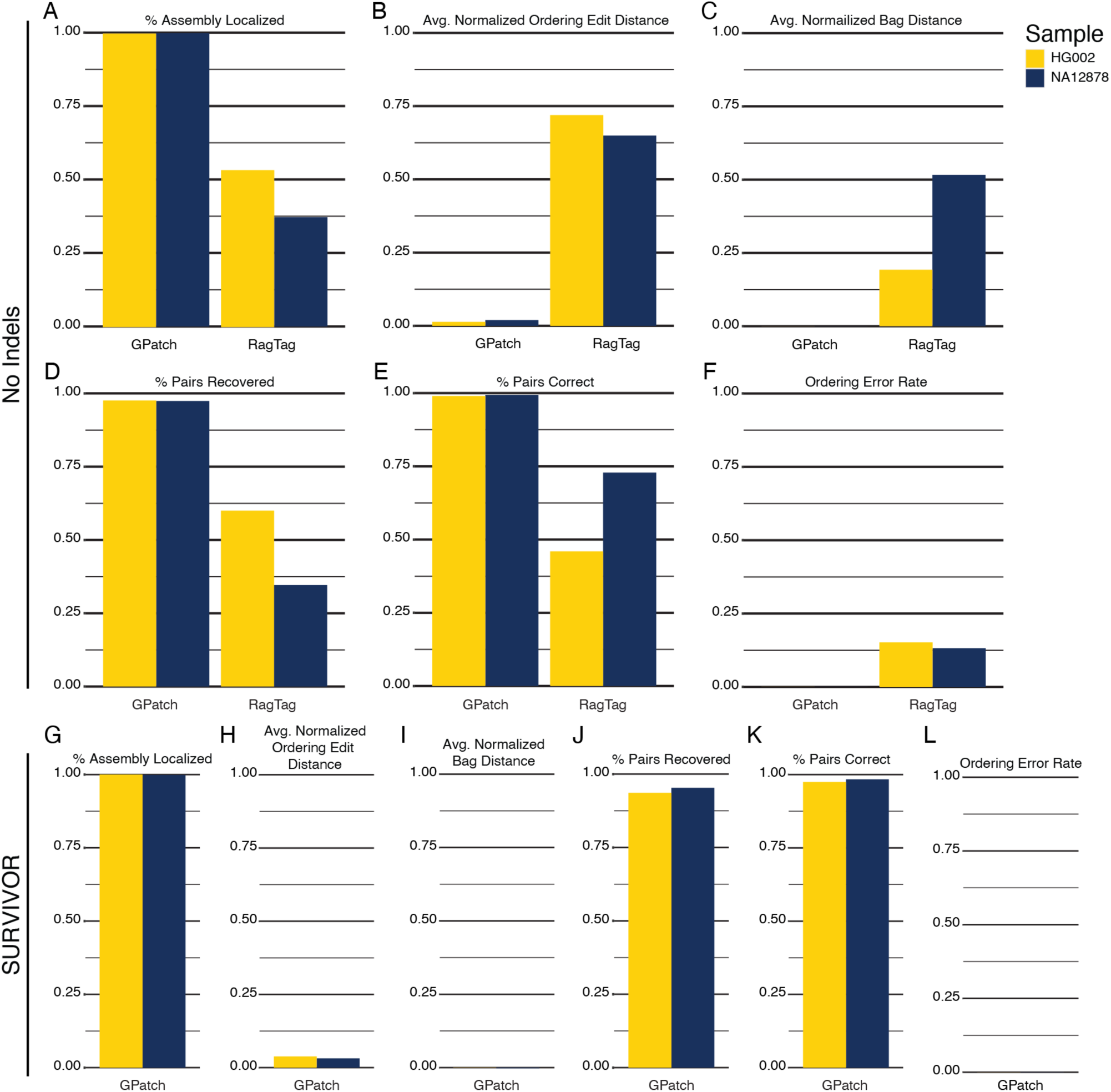
Simulation Analysis Results. All panels: Maize bars = HG002, Blue bars = NA12878. A and G) The percentage of nucleotides from the target assembly localized in the patched assembly. B and H) Average Levenshtein distance between the observed and actual contig ordering vectors divided by the length of the target sequence. C and I) Average bag distance between patched and target chromosome sequences divided by target sequence length. D and J) The fraction of adjacent contig pairs from the target genome recovered in the patched genome. E and K) The fraction of adjacent pairs in the patched genome that are also found in the target genome. F and L) The fraction of contigs placed out-of-order relative to neighboring contigs in the patched sequence. Alt text: Simulation analysis results expressed as bar charts describing relative performance of GPatch and RagTag patch on simulated datasets.

By contrast, RagTag patch performed poorly despite extensive optimization of patching parameters and experimentation with alternate aligners. With default patching parameters, RagTag Patch dropped most contigs from the output pseudoassembly, producing a small set of incomplete scaffolds and falling well short of yielding the desired complete pseudoassembly. Optimizing input parameters improved performance marginally but the best performance was about 47% contig placement (Supplementary Table S3). We also noted there is not a 1-to-1 relationship between reference chromosomes and scaffolds in the RagTag patch results. First, entire chromosomes were omitted from the patched pseudoassembly, with scaffolds recovered for only nine chromosomes in NA12878 and twelve in HG002, many of which were truncated and/or contained large, likely-spurious, rearrangements relative to T2T-CHM13 (Supplementary Fig. S4-S5). Second, we noted that, in NA12878, chromosomes 5, 12, and X were each split into three separate scaffolds. Third, we observed that three pairs of chromosomes in the HG002 results (14-21; 3-19; 4-15) were incorrectly joined into hybrid scaffolds. Finally, we noted that scaffolds always begin and end with a contig sequence, such that scaffolds are always 5’ and/or 3’ truncated in the event that contig sequence does not capture the entirety of the telomere. In all, RagTag pseudoassemblies captured only 43.07% of contigs for NA12878 and 47.39% for HG002, representing only 37.16% and 53.41% of their target assemblies, respectively (Fig. 2A). Likewise, pair recall of 34.60% for NA12878 and 60.0% for HG002 (Fig. 2D), and ordering edit distances of 64.93% and 71.91%, respectively, (Fig. 2B) indicate a high rate of dropped contigs. Furthermore, the combination of pair accuracies of only 72.86% and 45.95% (Fig. 2E), ordering error rates of 13.20% and 15.97% (Fig. 2F) for NA12878 and HG002, respectively, indicate relatively high rates of contigs being placed in the wrong relative order when compared to their neighbors in the target assembly. Normalized bag distances of 51.65% and 19.25% (Fig. 2C) suggest large-scale sequence differences between the patched pseudoassemblies and target assemblies. Dot-plots between the patched pseudoassemblies and T2T-CHM13 show that RagTag Patch was able to successfully scaffold only three chromosomes from NA12878 and nine from HG002 (Supplementary Fig. S4-S5).

To evaluate the effects of nucleotide-level and structural variation relative to the reference assembly on patching, SURVIVOR assemblies for NA12878 and HG002 were processed with GPatch and RagTag patch using the alignment and patching parameters optimized for the no-indel sets. GPatch maintained its high performance even in the presence of variation, localizing 96.88% of NA12878 and 96.15% of HG002 contigs, representing 99.83% and 99.96% of their respective simulated draft assemblies (Fig. 2G) and recovering 95.29% and 93.60% of adjacent contig pairs (Fig. 2J), with ordering edit distance remaining below 5% in both patched pseudoassemblies (Fig. 2H). Placement accuracy remained high, with 99.99% of NA12878 and 100% of HG002 contigs placed on the correct chromosome and strand, and pair-accuracy exceeding 97% (Fig. 2K) and negligible ordering error rates (Fig. 2L) for both patched pseudoassemblies. Likewise, average normalized bag distances of 0.24% for NA12878 and 0.04% for HG002 (Fig. 2I) indicate minimal sequence content divergence from their respective target assemblies. Dot-plots show nearly one-to-one correspondence between patched pseudoassemblies and target assemblies (Supplementary Fig. S6-S7). In contrast to the no-indel set, while still relatively short compared to contig lengths, patches were distributed more broadly and randomly across patched chromosomes (Supplementary Fig. 3C-D). Not surprisingly, we observed many more patches in the NA12878 genome compared to HG002, which we attribute to more uncertainty in the alignment of shorter contigs to the reference assembly, particularly in repetitive regions. We were unable to obtain results for comparison from RagTag patch, which consistently failed to scaffold any contigs from either SURVIVOR pseudoassembly, despite extensive optimization of input parameters, even when allowing RagTag patch to use its preferred aligner, nucmer [54]. Since RagTag patch failed to produce any output, we decided to test whether RagTag scaffold could successfully scaffold SURVIVOR pseudoassemblies. In contrast to RagTag patch, RagTag scaffold successfully scaffolded all chromosomes in both NA12878 and HG002 SURVIVOR simulated draft assemblies, although overall performance in all metrics was slightly below that observed for GPatch on the same datasets (Supplementary Fig. S8, Supplementary Table S3), with the largest difference being an approximate 2-fold increase in the average ordering edit distance (Supplementary Fig. S8G) relative to GPatch.

We additionally tested the effects of the choice of mapping quality (mapq) score thresholds on GPatch performance using the NA12878 SURVIVOR simulated draft assembly, at mapq thresholds 0, 10, 20, 30, and 40 (Supplementary Fig. S9). We observed only modest effects on overall performance, with the strongest effect being on the fraction of contigs placed (Supplementary Fig. S9A), ranging between 99.4% and 92.3% for mapq levels from 0-40. However, the fraction of the target assembly localized in the patched pseudoassembly remained nearly unchanged (Supplementary Fig. S9B), ranging from 99.98%-99.52%. Accuracy metrics were, likewise, only modestly affected (Fig S9E-H), with the largest effect observed being an approximately 8-fold increase in the ordering error rate observed for mapq 0 vs 10 (Supplementary Fig. S9F), reflecting a relatively high rate of incorrect mappings at quality scores below 10. For mapq 10-40, this rate remained relatively stable, manifested by inclusion of the same two misplaced contigs across patched pseudoassembles at all mapq levels. This result underscores the reliance of GPatch on the quality and completeness of the input alignment. Overall, the predominant reason for non-inclusion of contigs at mapq thresholds above zero was failure to meet the mapq threshold, while 2.6%-8.1% of contigs were dropped because their mapped intervals were nested within another placed contig (Supplementary Table S4).

### Patching NA12878 and HG002 Draft Genomes

To evaluate GPatch’s ability to construct chromosome-scale pseudoassemblies from biological data, we patched the same draft assemblies for NA12878 and HG002 used to model contig-length distributions in our simulation analysis. This represents a highly-fragmented assembly, NA12878, comprised of 10,406 contigs with a median length of 16,248bp, and a highly contiguous assembly, HG002, comprised of 762 contigs with a median length of 40,201bp. RagTag patch was not included in this analysis due to its failure to produce any scaffolds when applied to the same datasets. Contigs were initially mapped to T2T-CHM13 with minimap2 [51], and resulting BAM files were processed with GPatch using optimized parameter values from our simulated data analysis and the same mapq threshold of 10 for consistency. Patched genomes were evaluated for completeness using contig placement (Fig. 3A), the fraction of nucleotides from the draft assembly localized in the patched pseudoassembly (Fig. 3B), and for composition using the fraction of nucleotides in the patched pseudoassembly derived from contigs (Fig. 3C), with the goals of maximizing the rates for each metric. We also used recently-published chromosome-scale assemblies for both NA12878 (T2T-NA12878) [16] and HG002 (T2T-HG002) [52] as target assemblies for direct sequence comparison via dot-plots.

**Figure 3:**
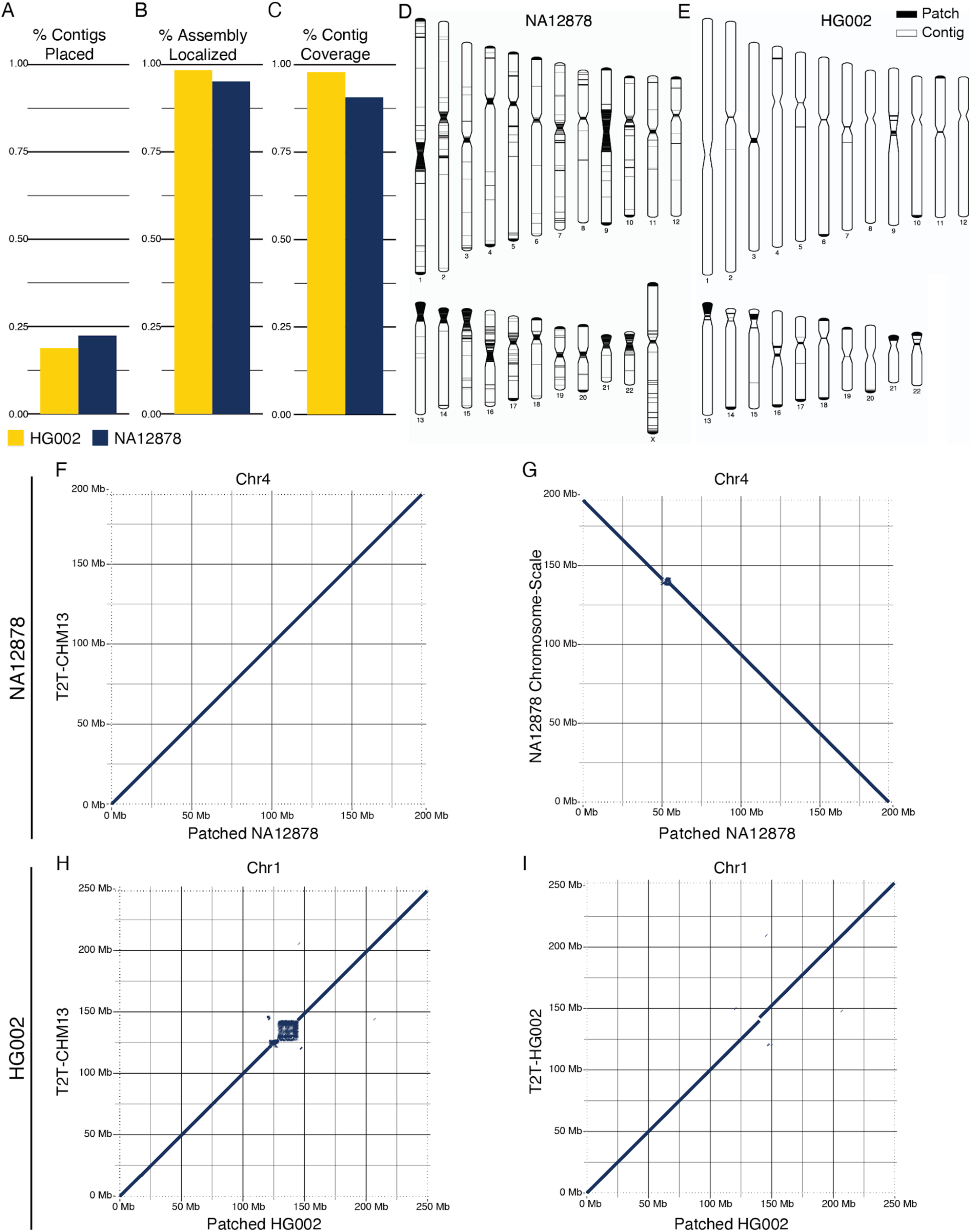
Patching Results for Biological Datasets. Panels A-C: Maize bars = HG002, Blue bars = NA12878. A) The fraction of contigs from the source assembly placed in the patched assembly. B) The percentage of nucleotides from the source assembly localized in the patched assembly. C) The fraction of nucleotides in the patched genome derived from contig sequences. D-E) Ideograms illustrating the locations of contigs and patches along each chromosome in the patched genome. D) HGSVC NA12878 genome patched with T2T-CHM13. E) HPRC HG002 genome patched with T2T-CHM13. F-I) Dot plots of patched chromosomes against T2T-CHM13 (F and H), or a chromosome-scale assembly for the same cell line (G and I). F) Patched HGSVC NA12878 chr4 aligned to T2T-CHM13 chr4. G) Patched HGSVC NA12878 chr4 aligned to full-length chr4 sequence from Porubsky et al. H) Patched HPRC HG002 chr1 aligned to T2T-CHM13 chr1. I) Patched HPRC HG002 aligned to T2T-HG002 chr1. Alt text: Bar charts showing GPatch performance on HG002 and NA12878 draft assemblies; ideograms showing patch locations in resulting pseudoassemblies; dot plots showing patched pseudoassembly contiguity relative to the reference.

We were surprised to find that only 22.02% of HGSVC NA12878 contigs, and 18.77% of HPRC HG002 contigs were placed in their respective patched genomes (Fig. 3A, Supplementary Table S5). However, there was a distinct bias toward placement of longer contigs, with placed contigs being significantly longer than unplaced contigs (NA12878: p-value=7.57e-167, HG002: p-value=2.25e-61, Wilcoxon Rank-Sum tests), evident as a large difference between placed and unplaced contig lengths at all quartiles (Supplementary Table S6). As a result, patched pseudoassemblies still captured the majority of nucleotides in each draft assembly: 94.25% for NA12878, and 95.13% for HG002 (Fig. 3B). Likewise, patched pseudoassemblies consisted primarily of contig-derived nucleotides: 88.9% for NA12878 and 97.85% for HG002 (Fig. 3C). As expected, patched pseudoassemblies substantially increased in N50 relative to their respective draft genomes, approaching values observed for the T2T-CHM13 reference assembly and their respective target assemblies (Supplementary Table S5). As we observed in simulated datasets, the predominant reason for non-inclusion of a contig was failure to meet the mapq threshold (NA12878=90.9%, HG002=64.9%), followed by nested mappings (NA12878=9.0%, HG002=20.5%), and finally no mapping (NA12878=0.16%, HG002=14.5%) (Supplementary Table S7).

Consistent with our expectations, we observed that the longer contigs in the HG002 assembly yielded better performance in terms of nucleotide localization and contig coverage in the patched pseudoassembly, with the HG002 pseudoassembly containing fewer and smaller patches than the NA12878 pseudoassembly (Fig. 3D-E). Indeed, HG002 chromosomes 1, 8, and 12 contained only contig sequence with no intervening patches (Fig. 3E). In both patched pseudoassemblies, patches tended to concentrate in/near telomeres, pericentromeric regions, and acrocentric short arms, in some cases contributing the majority of sequence in these regions. This likely reflects the repetitive nature of these regions, which tend to harbor shorter contigs and have reduced mapping ability relative to non-repetitive regions.

In order to evaluate large-scale genome structure, we compared patched pseudoassemblies to the T2T-CHM13 reference assembly and respective chromosome-scale target assemblies using dot-plots (Fig. 3F-I). Both patched genomes were found to be highly collinear with T2T-CHM13 (Fig. 3F, 3H, Supplementary Fig. S10-S11), with most departures associating with pericentromeric regions (Fig. 3H). Aside from these regions, we observed large-scale structural differences between the GPatch NA12878 pseudoassembly and T2T-CHM13 on six chromosomes (Fig. 4A, Supplementary Fig. S10), however, no comparable divergence was noted between the GPatch HG002 pseudoassembly and T2T-CHM13 (Supplementary Fig. S11). All the divergent loci noted in NA12878 included at least one breakpoint within a placed contig, suggesting they represent either true structural variants or misjoins in the source assembly rather than artifacts of the patching process. Dot plots of patched pseudoassemblies against their respective target assemblies showed similar patterns, largely recapitulating those obtained from comparisons with the T2T-CHM13 reference (Fig. 3G, 3I, Supplementary Fig. S12-S13). This suggests that patched pseudoassemblies indeed recapitulate their target assemblies, at least at large-scale. Indeed, in some cases, we observed closer correspondence between the patched pseudoassemblies and target assemblies than with the T2T-CHM13 reference, as in HG002 chr1 (Fig. 3H-I).

**Figure 4:**
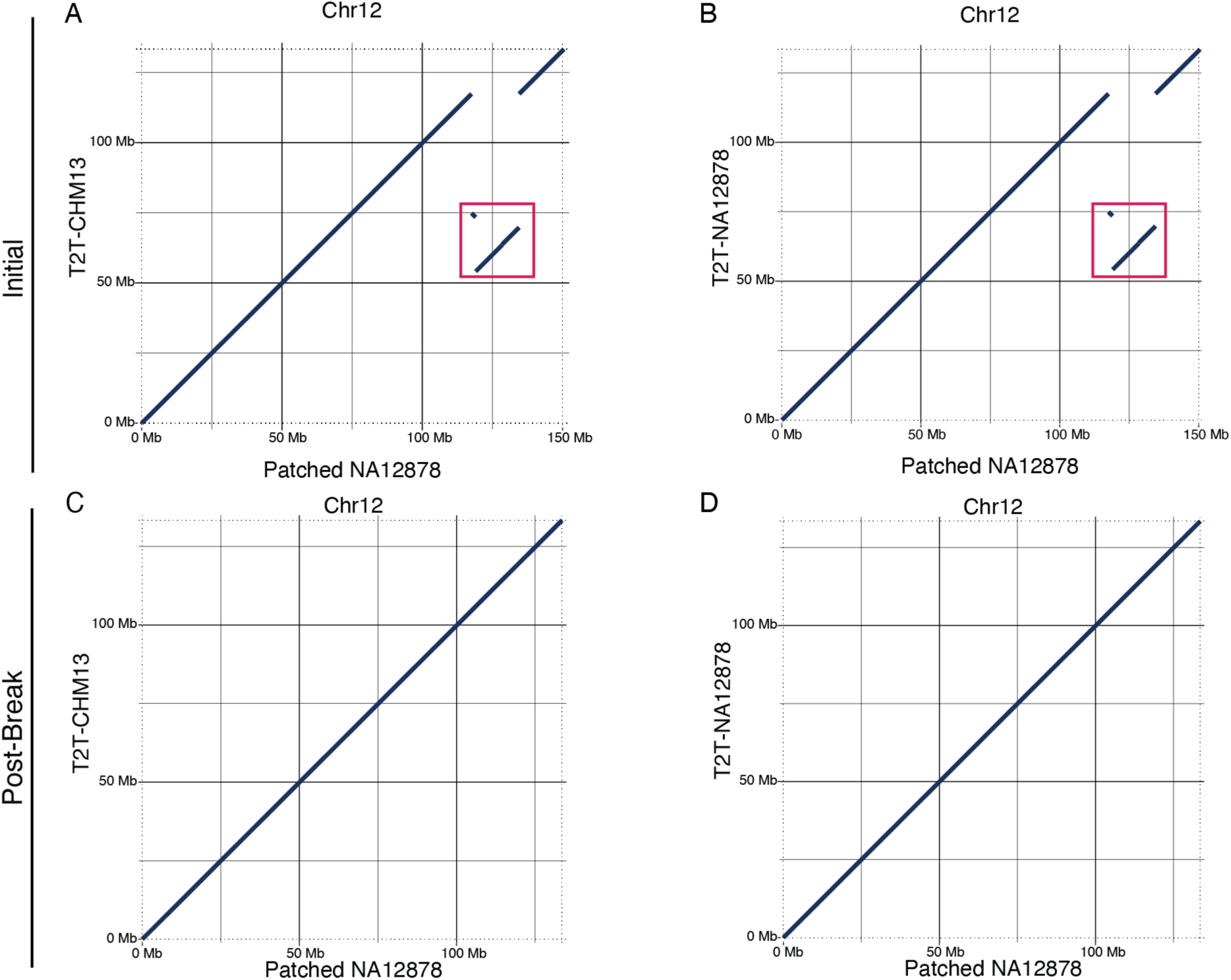
Identifying and Correcting Likely Misjoins. All Panels: Dot plots of patched NA12878 chromosomes against T2T-CHM13 (A and C), or a chromosome-scale assembly NA12878 (B and D). A) Patched HGSVC NA12878 chr12 aligned to T2T-CHM13 chr12. The red box indicates the location of two apparent misjoins. B) Patched HGSVC NA12878 chr12 aligned to full-length chr12 sequence from Porubsky et al. The red box indicates the location of two apparent misjoins. C) Patched NA12878 chr12 after a single-round of contig-breaking at misjoins visible in A and C, aligned to T2T-CHM13 chr12. D) Patched HGSVC NA12878 chr12 after a single round of contig-breaking at misjoins visible in A and C, aligned to full-length chr12 sequence from Porubsky et al. Alt text: Dot plots of patched NA12878 assembly against T2T-CHM13 and Porubsky NA12878 showing discontinuities at putative misjoin loci.

### Patching the Tomato M82 Genome

To demonstrate GPatch’s ability to patch non-human assemblies, we obtained the S. lycopersicum M82 assembly, build 1.3 (M82) [48], and the Heinz 1706 reference assembly, build 3.0 (SL3) [50] from the Sol Genomics Network (Supplementary Table S1). As the M82 assembly was presented as scaffolded contigs, we first extracted contig sequences by splitting input FASTA sequences at stretches of N characters >= 20bp in length. The resulting dataset contained 1,597 contigs with a median length of 103,435bp. Contigs were initially mapped to SL3 with minimap2 [51], and resulting BAM files were processed with GPatch using the same set of parameters previously used with the human datasets. Patched genomes were evaluated for completeness using contig placement, the fraction of nucleotides from the draft assembly localized in the patched pseudoassembly, and for composition using the fraction of nucleotides in the patched pseudoassembly derived from contigs (Supplementary Table S5).

The patched M82 pseudoassembly totalled 850,138,583 bp in length, incorporating 98.49% of nucleotides, spanning 73.95% of contigs from the input assembly, and consisting of 91.83% contig sequence. As with the human data, placed contigs significantly exceeded unplaced contigs in length across all quartiles (Supplementary Table S6; p-value=8.21e-125, Wilcoxon rank-sum test). Likewise, the patched pseudoassembly increased markedly in N50 compared to the input contigs, approaching the N50 value of the target M82 assembly (Additonal File 2: Table S5). In contrast to the human data, the predominant reason for contig non-placement was nesting within other contig mappings, accounting for 72.36% of dropped contigs, with failure to reach the mapq threshold and non-mapping accounting for 17.55% and 10.10% of dropped contigs, respectively.

To evaluate large-scale genome structure, we produced dot-plots for each chromosome for the patched M82 pseudoassembly compared to both the SL3 reference assembly (Supplementary Fig. S14) and the unfragmented M82 assembly (Supplementary Figure S15). We observed a high degree of collinearity with the SL3 reference assembly for all chromosomes in the patched assembly. However, most chromosomes contained at least one gap of approximately 1-5MB, relative to the SL3 reference, which likely correspond to repetitive regions with poor mappability. Likewise, dot-plots to the unfragmented M82 assembly were highly collinear, with fewer and smaller gaps overall compared to alignments to SL3. The notable exceptions were chromosomes 2, 3, and 6, which all contained insertions of approximately 2-4Mb relative to M82, all three of which appear to be SL3-derived sequences in the patched pseudoassembly. These sequences appear to be absent from the M82 assembly based on reciprocal patterns visible in the same regions of dot plots between alignments of the unfragmented M82 assembly to SL3 (Supplementary Fig. S16). On chromosome 3, we also noted the presence of a large inversion of ∼8MB. This locus contained involves 10 contigs from the input M82 assembly, 6 of which were derived from the same locus in the original M82 assembly, and four of which were derived from M82 chromosome 0, which is derived from unscaffolded contigs in the original M82 assembly. This inversion is clearly visible in dot plots comparing unfragmented M82 to SL3 (Supplementary Fig. S16); this and the involvement of multiple contigs seems to rule out the presence of a misjoin. If we accept the unfragmented M82 assembly as the ground truth, we must conclude that these four variants within the patched pseudoassembly reflect reference bias introduced by the patching process based on the alignment to SL3.

### Automated Misjoin Correction

When examining dot plots of the GPatch NA12878 pseudoassembly against T2T-CHM13, we noted large-scale rearrangements on chromosomes 3, 7, 9, 12, 17, and 22 (Fig. 4A, Supplementary Fig. S10). Since these same rearrangements were present in dot-plots between the GPatch NA12878 pseudoassembly and full-length chromosomes from a recently-published chromosome-scale NA12878 assembly [16] (Fig. 4B, Supplementary Fig. S12), we were able to exclude the possibility that these reflect genuine structural variants in NA12878 relative to T2T-CHM13. Therefore, these rearrangements must reflect misjoins in the HGSVC NA12878 assembly. Indeed, distinct off-diagonal signals in Hi-C data mapped to the GPatch NA12878 pseudoassembly confirmed the presence of these misjoins (Supplementary Fig. S17). Since these features are likely to negatively affect performance when using patched pseudoassemblies in place of a reference assembly in downstream analyses, we developed a method to break contigs at the coordinates of likely misjoins prior to realignment and patching with GPatch.

When applied to the HGSVC NA12878 pseudoassembly, we identified 22 loci on nine chromosomes representing likely misjoins. After one round of contig-breaking, we observed no grossly-visible rearrangements on any chromosome, except chromosome 9, when comparing the GPatch pseudoassembly to T2T-CHM13 (Fig. 4C, Supplementary Fig. S18), and to full-length NA12878 chromosomes (Fig. 4D, Supplementary Fig. S19). A second-round of contig-breaking identified two additional loci on chromosome 9 that appeared to originate from a contig that was absent from the initial patched pseudoassembly. The final patched pseudoassembly, after two rounds of contig-breaking, had no grossly-visible rearrangements when compared to either T2T-CHM13 or full-length NA12878 chromosomes (Supplementary Fig. S20-S21). Overall, contig-breaking appeared to improve the quality of the patched NA12878 pseudoassembly, as indicated by improved metrics for the percentage of NA12878 nucleotides recovered in the patched pseudoassembly and the fraction of the patched pseudoassembly derived from contigs after each contig-breaking iteration (Supplementary Table S5). This suggests that contig-breaking at likely misjoins is beneficial for overall patching performance.

Although we did not observe any large-scale rearrangements in the GPatch HG002 pseudoassembly (Supplementary Fig. S11&S13) comparable to those we saw in NA12878 data (Supplementary Fig. S10&S12), our script did detect 48 potential misjoins based on the alignment to T2T-CHM13. However, breaking contigs at the corresponding loci actually decreased performance relative to the initial patched pseudoassembly as indicated by drops in both the fraction of contig nucleotides captured and contig nucleotide coverage (Supplementary Table S5) while dot-plots against T2T-CHM13 and T2T-HG002 remained largely unchanged (Supplementary Fig. S22-S23). Furthermore, contig-breaking caused the patched pseudoassembly to grow in size, mostly reflecting a 38.5% increase in patch nucleotide content. We conclude that contig-breaking is not beneficial when alignments between the patched pseudoassembly and the reference do not reveal any large-scale divergence. Therefore, caution should be exercised when applying contig-breaking, paying particular attention to its effects on contig nucleotide recovery and patched pseudoassembly size and composition.

### Patched Pseudoassemblies Recapitulate Hi-C Patterns in T2T-CHM13

To show that GPatch pseudoassemblies are interchangeable with reference assemblies in genomic analyses, we obtained publicly available Hi-C sequencing data for NA12878 [31] and processed it through the loop-calling stage using the 4D-Nucleome Hi-C analysis pipeline [55] using the GPatch NA12878 pseudoassembly as the reference genome. For comparison, we analyzed the same dataset using the T2T-CHM13 reference assembly, and the HGSVC NA12878 draft assembly (Supplementary Table S8), using the same Hi-C data and analytical pipeline. At the mapping level, we noted similar rates of overall read mapping (99.44% and 99.45%, respectively) and mate mapping (98.94% and 98.95%, respectively) between GPatch NA12878 and T2T-CHM13. However, GPatch NA12878 actually outperformed HGSVC NA12878 (% Mapped = 99.09%, % Mate Mapped = 98.35%). Notably, reads where the mate maps to a different chromosome occurred at a ∼53% higher rate in HGSVC NA12878 (39.35%) than in either GPatch NA12878 or T2T-CHM13 (25.74% and 25.83%, respectively). This disparity remains evident when mapped reads are decomposed to individual pairs of genomic contact loci, with 35.37% of read pairs split across contigs in HGSVC NA12878 compared to 20.95% in GPatch NA12878 and 21.01% in T2T-CHM13. While many of these likely represent legitimate interchromosomal contacts, some reflect “cryptic” intrachromosomal contacts which cannot be recovered in the draft genome because their anchor loci are split between different contigs. GPatch allows many of these cryptic contacts to be recovered.

Pairs files for T2T-CHM13 and GPatch NA12878 were assembled into Hi-C matrices and normalized with Juicer Tools [56]. However, for HGSVC NA12878, matrix construction failed with an out-of-memory error, having exhausted all available RAM (1.5TB) on a memory-optimized compute node. Since differences in the coordinate systems between T2T-CHM13 and GPatch NA12878 genomes precluded direct, quantitative matrix comparisons, such as stratum-adjusted correlation coefficient, Hi-C matrices were plotted as heat maps for visual comparison (Supplementary Fig. S24-S25). We observed very few grossly-visible differences between heat maps prepared using GPatch NA12878 and T2T-CHM13, with GPatch NA12878 heat maps largely recapitulating patterns seen with T2T-CHM13 (Fig. 5, Supplementary Fig. S24-S25). These results suggest that GPatch assemblies can substitute for reference assemblies, while being composed mostly of sequence derived from the draft assembly, whereas the unpatched draft assembly cannot serve interchangeably as a reference assembly.

**Figure 5:**
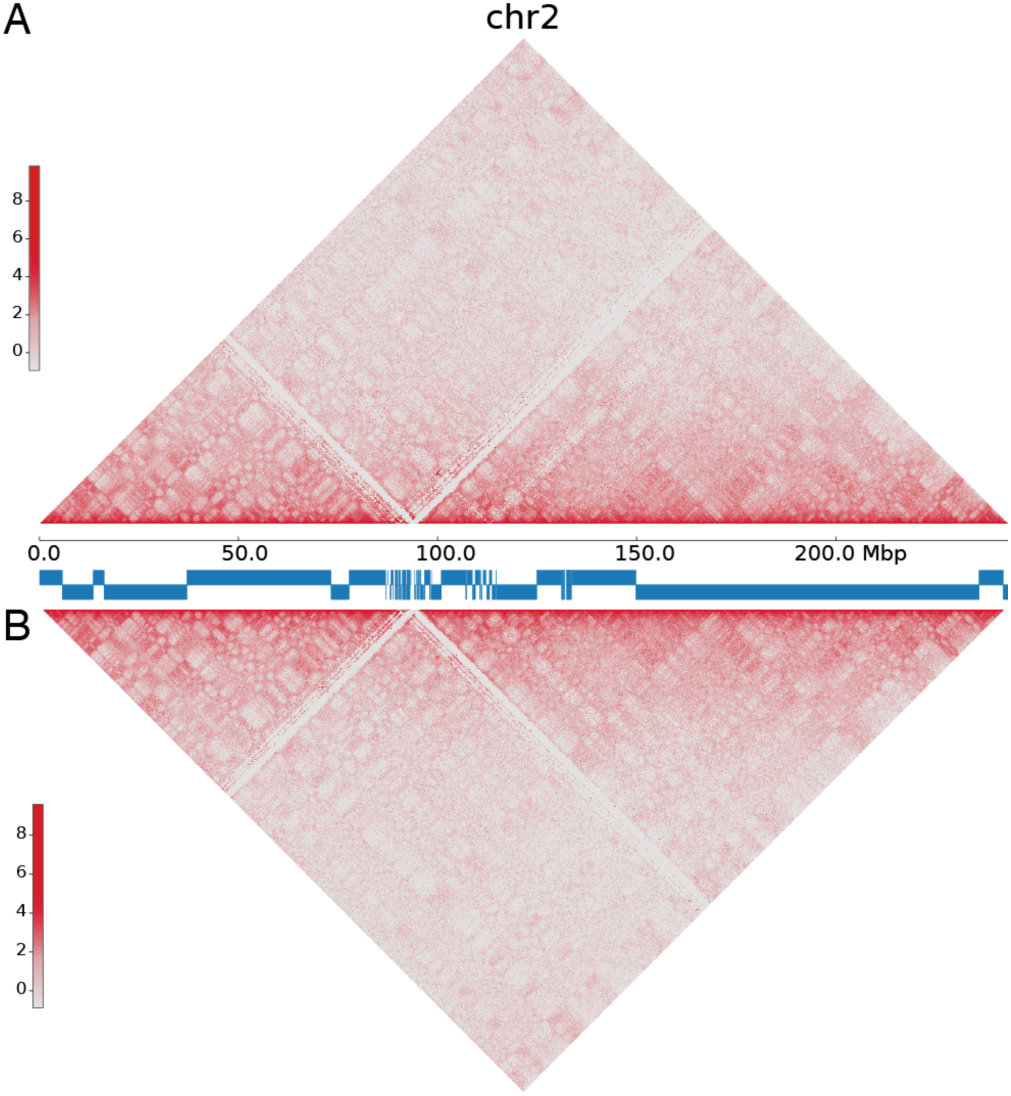
NA12878 Hi-C Heat Maps. All panels: Hi-C heat maps showing contact density at 80kb resolution with balanced normalization, with the X-axis indicating the position on human chromosome 2. A) NA12878 Hi-C data mapped to the HGSVC NA12878 assembly patched with T2T-CHM13. Blue bars below the X-axis indicate the positions of contigs from the original NA12878 assembly within patched chromosome 2, whereas blank regions indicate patches taken from the reference sequence at corresponding loci. B) The same NA12878 Hi-C data mapped to T2T-CHM13. Alt text: Heat maps showing equivalent patterns when NA12878 Hi-C data are mapped to T2T-CHM13 and Patched NA12878.

### GPatch Enables Recovery of Cryptic Loops from the HGSVC NA12878 Genome

Hi-C matrices for both T2T-CHM13-mapped and GPatch NA12878-mapped data were processed with hiccups [56] to produce chromatin loop predictions at 5kb and 10kb resolution. At both resolutions, we found comparable numbers of loops at comparable average sizes at using both data mapped to GPatch NA12878 and T2T-CHM13 (Supplementary Table S8), showing that the GPatch pseudoassembly is, again, able to recapitulate results produced using a reference assembly. However, we did identify notable differences between the GPatch NA12878 and T2T-CHM13 data. First, we observed many loops at both resolutions for which the 5’ and 3’ anchors map to different contigs in HGSVC NA12878 assembly. We call these cryptic loops, since they would be impossible to recover in data mapped directly to HGSVC NA12878. We recovered 126 cryptic loops (∼1.29% of all loops) at 5kb resolution, and 599 cryptic loops (∼2.99% of all loops) at 10kb resolution, demonstrating that GPatch allows recovery of genomic features that would otherwise be missed using an unpatched draft assembly.

In order to directly compare loop predictions from GPatch NA12878 with those from T2T-CHM13, we first lifted over GPatch NA12878 loop anchor loci at both resolutions to the T2T-CHM13 coordinate frame. These were then intersected with T2T-CHM13 loop anchors using BEDtools [57]. Lifted and intersected data were then used to reconstruct individual loops in order to determine the number of fully-shared, partially-shared, and NA12878-only loops (Supplementary Table S8). In total, we observed 88.86% of GPatch NA12878 loops at 5kb resolution and 95.21% of loops at 10kb resolution that shared at least one anchor with T2T-CHM13. This leaves a substantial fraction of loops for which neither loop anchor is shared with T2T-CHM13 data, with ∼95% of these existing in sequence that is physically present in both the reference assembly and the GPatch NA12878 pseudoassembly. Notably, we detected 66 loops at 5kb resolution and 137 loops at 10kb resolution for which at least one anchor is in sequence unique to GPatch NA12878. This included four 5kb loops and five 10kb loops for which neither anchor could be mapped to T2T-CHM13. Thus, GPatch pseudoassemblies allow recovery of genomic features that would otherwise be missed when using a reference assembly.

## DISCUSSION

The combination of long-read sequencing and advanced algorithms for de-novo genome assembly have yielded an abundance of publicly-available draft personal genomes. By capturing donor-specific variations relative to reference assemblies, these genomes carry the promise of increased sensitivity and specificity when used in place of a reference assembly for mapping and downstream genomic analyses. However, most of these draft genomes exist as sets of hundreds to thousands of individual contigs, which are typically not assigned chromosomal identity. This presents several challenges that, ultimately, render these draft genomes unsuitable to replace a reference assembly for most genomic assays, particularly those relying on spatial relationships between genomic loci.

While recent advances in de-novo assembly have increased the contiguity of draft genomes, a number of unresolved challenges remain. Chief among these is their inability to completely assemble highly-repetitive genomic regions, such as centromeres, telomeres, HLA genes, the rDNA region [58], and acrocentric short arms [59], particularly in regards to phased genomes. Novel approaches, such as repeat graphs [29] minimizer hashing [27], and multiplex de Bruijn-graphs [25,59] have yielded notable improvements in ability to assemble such regions. However, none are currently able to produce T2T genomes without multiple data types [25,26] and/or extensive manual curation [30,58]. Until de-novo assembly software and/or long-read sequencing advance to the point where complete, T2T assemblies can be reasonably built in-house within a typical research lab, these challenges will limit the utility of long-read draft personal genomes. At present, this remains impractical and methods are needed to bridge the gap between draft genomes and reference assemblies. Here we present GPatch, which utilizes alignments to a reference assembly to order and orient contigs from draft genomes into gap-free, chromosome-scale pseudoassemblies.

We have demonstrated that GPatch faithfully produces chromosome-scale pseudoassemblies given only a draft assembly and a reference assembly, without the need for additional data types to assist in ordering and orienting contigs. The resulting patched pseudoassemblies consistently capture over 95% of nucleotides from their source draft assemblies, are composed of 89-98% contig sequence, and are highly collinear with both the reference and target genome assemblies, in both simulated and biological data analyses. GPatch outperformed the only competing software of which we are aware, RagTag Patch, in all metrics we tested. RagTag Patch performed poorly even on simulated data containing no indels, often partially-scaffolding chromosomes, splitting chromosomes across multiple scaffolds, and frequently dropping chromosomes from the output. Furthermore, RagTag Patch output is difficult to interpret since it does not name scaffolds according to their corresponding reference chromosomes, necessitating further steps to establish chromosomal identities. Last, since RagTag Patch will only terminate a scaffold with a contig, resulting scaffolds will always be incomplete if contigs do not completely encompass the telomeres. While RagTag scaffold, by comparison, performed well in our tests, gaps between mapped contigs are padded with N characters, not sequence form the reference assembly, thus leaving their output pseudoassemblies incomplete. GPatch answers all these shortcomings. By looping over reference chromosomes and tracking whether contig alignments extend to the ends of telomeres, the GPatch algorithm guarantees inclusion of full-length, gapless scaffolds for all reference chromosomes in the output, with all mapped contigs assigned unambiguously to a chromosome. Therefore, GPatch pseudoassemblies are gap-free, complete with respect to target chromosome extent, and can be used in place of reference genome assemblies for Hi-C and other genomics applications for which genome fragmentation is undesirable.

The largest potential drawback of GPatch is its reliance on contig alignments to a reference genome. We have shown that alignment quality and completeness significantly impact the content of GPatch assemblies. Indeed, alignment quality is the primary reason contigs are dropped from GPatch genomes, with many dropped contigs mapping to genomic regions that are traditionally difficult to align within, such as pericentromeric and telomeric repeat arrays. This implies that donor-specific variation within these regions is likely to be missed in GPatch alignments, thus perpetuating reference bias. It is important to note that this limitation is not unique to GPatch, but will, by definition, be shared by any approach to genome patching, or scaffolding, that relies on alignment to a reference assembly. Likewise, for any approach relying on alignment to a reference assembly, the choice of aligners and specific alignment parameters is of great importance. While we use minimap in this analysis, it is important to note that GPatch will work with the output of any aligner capable of outputting alignments in SAM/BAM format. Of particular interest are long-read aligners, such as DeChat[60] and Winnowmap[61], designed to accommodate repeat-rich regions, which are typically enriched for low-quality and erroneous alignments, and which, in turn, host the majority of patch sequences in our results.

Given the strong reliance on accurate alignments, the specific choice of reference assemblies will also greatly influence the accuracy of GPatch and all reference-guided approaches, whether scaffolding or patching are the ultimate results. Indeed, it is important to choose a high-quality reference assembly that is as closely related as possible to the target genome GPatch is intended to approximate. Intuitively, as divergence between the reference assembly and target genome increases, so will the number and extent of structural differences between them, resulting in an elevated rate of ordering and orientation errors within the patched pseudoassembly. Furthermore, reduced mapping quality is likely to increase the number of contigs dropped from the final assembly, leading to decreased contig coverage in the patched pseudoassembly and a concomitant increase in patch content. Since patches are likely to contain some divergent nucleotides relative to the target sequences they approximate, this will likely artificially increase the divergence rate between the patched pseudoassembly and the target genome. Carried to one extreme, GPatch will happily generate a patched pseudoassembly from contig data aligned to a reference assembly that contains a different number of chromosomes than the target genome, e.g., one from another species. The result would then contain the wrong number of chromosomes, and likely harbor many structural inaccuracies and patches containing DNA foreign to the target genome. For this reason, we advise against applying GPatch to draft and reference assemblies that are not derived from the same species. Should it be necessary to employ a divergent reference assembly, running GPatch with the –scaffold_only option, available in releases 0.4.0 and greater, will cause gaps between contigs to be filled with N characters rather than reference-derived patch sequences, thus eliminating the possibility of foreign sequence inclusion.

At present, no currently-available reference-free approach can patch a typical draft genome de-novo in the absence of extremely deep long-read sequencing and/or additional data that are typically unattainable for individual researchers. Furthermore, reference-free approaches are typically much slower than reference-guided approaches and yield genomes containing unresolved N-gaps that reduce mappability. Therefore, de-novo approaches are often unsatisfactory, and do not represent an alternative to reference-guided patching. Accepting a degree of reference bias seems, at present, to be a necessary compromise in utilizing draft personal genomes to their full potential in genomics analyses. GPatch answers the noted shortcomings of de-novo scaffolding, yielding gap-free, chromosome-scale pseudoassemblies that we show can be used in place of reference assemblies for most applications, albeit with the inevitable inclusion of some reference bias, as seen on tomato M82 chromosomes 2, 3, and 6. As for completeness, it is true that the upper-bound on completeness of GPatch pseudoassemblies is the completeness of the input draft personal genome. In practice, many short contigs from the draft assembly are lost from the final GPatch pseudoassembly. These dropped contigs are likely enriched for long repeats and perhaps structural variants, lost due to shortcomings in our ability to confidently align these sequences given current technology. As mentioned above, repeat-aware aligners may alleviate this problem to a degree. However, unless all contigs can be confidently placed, patch sequences will inevitably be substituted in place of dropped contigs, in some cases erroneously introducing sequence that does not match the equivalent locus in the personal genome, leading to a simultaneous decrease in completeness and increase in reference bias inherent to the patched pseudoassembly.

Indeed, it is important to acknowledge that, owing to uncertainty in the content and correctness of patch sequences, patched pseudoassemblies may contain sequences foreign to their target draft assemblies. This, in turn, may erroneously inflate estimates of sequence-level and structural conservation relative to the reference genome used, while artificially decreasing observed conservation relative to other related personal genomes. This should be taken into account when choosing whether patched pseudoassemblies are appropriate for any given analysis, weighing the potential advantages for sequenced read mapping against the potential for erroneous conservation conclusions. As an example, GPatch pseudogenomes may not be suitable for quantifying divergence between assemblies, or for comprehensive de-novo variant identification. However, here we demonstrate that GPatch pseudoassemblies can improve mapping performance in analysis of genomics data and alleviate the negative effects of genome fragmentation in downstream analysis. Furthermore, for applications where only the relative ordering of contigs is important, GPatch versions above 0.3.6 offer a scaffold-only mode, whereby gaps between contigs are padded with N characters rather than reference-derived sequences, thus avoiding overestimation of sequence-level conservation with the chosen reference assembly.

We demonstrate that GPatch pseudoassemblies, when aligned back to the reference genome, can highlight misjoins within contigs from the draft assembly. These typically appear as large-scale rearrangements visible in dot-plots between the GPatch pseudoassembly and the reference genome. The GPatch github repository includes a set of scripts designed to identify the boundaries of such events within an alignment of a GPatch pseudoassembly to the chosen reference assembly and break affected contigs at the corresponding breakpoint(s), allowing the resulting contig fragments to align independently within the reference genome. We show that, after contig-breaking, assemblies harboring misjoins perform better within GPatch, with resulting pseudoassemblies incorporating more contig sequence and less patch sequence, and being highly collinear with both the chosen reference and target assemblies. It is important to note that contig-breaking has the potential to obscure genuine structural variants, thus increasing reference bias. For this reason, we recommend caution when choosing whether to perform contig-breaking and secondary patching, with careful consideration of dot-plots as a necessary step in deciding whether to employ it. We note that the tendency to overcorrect can be minimized by careful choice of parameters in the breakpoint-identification step, in particular the minimum rearrangement size for breakpoint flagging. Importantly, if known structural variants are present in a draft genome, contig-breaking at these loci can be prevented by simply removing the corresponding records from the breakpoints text file used in the contig-breaking step.

Finally, we show definitively that GPatch pseudoassemblies can substitute for a reference assembly in Hi-C data analysis without sacrificing alignment rates or quality. Indeed, the GPatch NA12878 pseudoassembly achieved mapping and pairs-construction performance comparable to or better than either the T2T-CHM13 reference or the unpatched HGSVC NA12878 assemblies. By contrast, we demonstrate that unpatched draft assembly for HGSVC NA12878 could not be used in place of reference genome, with Hi-C analysis failing at the matrix construction stage. Importantly, even if matrix construction had succeeded for HGSVC NA12878, the challenges related to interpreting over 10,000 individual contact matrices cannot be overstated. Furthermore, since cis- and trans-interactions are conflated among these matrices, and since many cis-interactions are artificially split across contigs, loop calling and other spatially-based analyses will be negatively affected, leading to cryptic loops that cannot be detected even if the computational analysis could be completed. We show that GPatch allows us to recover a large number of these cryptic loops, which would otherwise be obscured since their 5’ and 3’ anchors are split between different contigs within the unpatched genome. However, it must be acknowledged that some of these loops may represent false positives resulting from misplaced contigs and/or “foreign” DNA sequences within patches. We believe that the benefits of obtaining results when it would be otherwise impossible outweigh this risk, but caution each user to carefully consider the implications in the scope of their own analyses.

Surprisingly, while up to 95% of loops we identified were at least partially-conserved (i.e., at least one anchor is shared between a GPatch NA12878 loop and a T2T-CHM13 loop), a surprisingly large number of loops included at least one anchor not shared with T2T-CHM13. Most of these are within sequence that is present in both genome assemblies, suggesting that these differences likely stem from differences in fine-mapping of reads in the NA12878 pseudoassembly relative to T2T-CHM13. That such small-scale differences can influence such a large number of loop predictions is striking, speaking to the importance of the “missing” variation captured in the GPatch NA12878 pseudoassembly but missing from T2T-CHM13 in its effects on nucleotide-level read mapping. While we cannot speculate on the biological consequences of such variation, GPatch makes it accessible to a broad range of downstream genomic methods, allowing us to gain critical information toward understanding the consequences of intraspecific variation.

Draft personal genomes based on long-read sequencing are becoming increasingly available. While these genomes offer the attractive prospect of mapping sequenced reads to a personal genome assembly matching the sequence donor for an assay, their fragmented nature presents significant obstacles to their use in functional genomics assays. In particular, assays based on spatial relationships between loci are unduly affected by genome fragmentation. Fragmented draft genomes present problems in both data-analysis, where many methods quickly become computationally intractable as the number of contigs increases, and interpretation, with the extreme example of interpreting over 10,000 individual Hi-C contact matrices had matrix construction for HGSVC NA12878 succeeded. GPatch overcomes these challenges, reducing a large number of individual contigs into a complete set of fully-assembled pseudochromosomes, given only a draft assembly and a reference genome. We show that GPatch pseudoassemblies are functionally interchangeable with polished reference genomes while incorporating over 95% of nucleotides from the source draft genome and achieving contiguity measures comparable to the reference assembly. These features enable their use in assays that would otherwise not be practical using the unpatched draft genome. Notably, we were able to utilize a GPatch pseudoassembly as the reference assembly for Hi-C data analysis and loop prediction, whereas the same analysis using the unpatched draft genome failed, despite generous resource allocations. We demonstrate that GPatch is robust to varying levels of genome fragmentation, and, more importantly, that it performs well on assemblies built using only resources already within the reach of many individual researchers. We conclude that, until it becomes realistic for individual labs to routinely build polished T2T genomes from scratch, methods for reliably assembling contigs into chromosome-scale pseudoassemblies are necessary to make the most of newly-available draft genomes. GPatch achieves this milestone, thus bridging the gap between draft genomes and polished reference assemblies.

## ACKNOWLEDGEMENTS

The authors would like to acknowledge members of the Boyle lab for helpful discussions and feedback provided throughout the analysis and manuscript preparation processes.

## AUTHOR CONTRIBUTIONS

AD designed and wrote the GPatch software, performed all data collection, preparation, and analysis steps, and prepared the figures and manuscript for submission. AB secured funding and provided oversight and guidance throughout the analysis. All authors read and approved the final manuscript.

## SUPPLEMENTARY DATA

Supplementary-Figures.pdf. Supplementary Figures. Figures S1-S24.

Supplementary-Tables.xlsx. Supplementary Tables. Tables S1-S8.

Supplementary Data are available at NAR online.

## CONFLICT OF INTEREST

The authors declare that they have no competing interests.

## FUNDING

This work was supported by the National Institutes of Health (R01GM144484).

## DATA AVAILABILITY

GPatch is open-source and available for download at https://github.com/adadiehl/GPatch under the MIT license. All analyses were performed using GPatch release 0.3.6, available at https://github.com/adadiehl/GPatch/archive/refs/tags/0.3.6.tar.gz. Please note that 0.3.6 is deprecated and the authors recommend using version 0.4.0 or later. GPatch is implemented in Python, and designed to run on the Linux operating system. GPatch requires Python >= 3.7, samtools (https://github.com/samtools/samtools), biopython (https://biopython.org/), and pysam (https://github.com/pysam-developers/pysam). Custom scripts used to identify misjoin breakpoints and perform contig-breaking are included in the main github repository, and documented at https://github.com/adadiehl/GPatch/tree/master/scripts. Tools in the scripts directory also require minimap2 (https://github.com/lh3/minimap2). All custom code and commands used in this analysis are documented and presented in the manuscript github repository at https://github.com/Boyle-Lab/GPatch-Manuscript. Sources for all publicly-available data used in this study are documented in Supplementary Table S1.

## Notes

https://github.com/Boyle-Lab/GPatch-Manuscript

### Competing Interest Statement

The authors have declared no competing interest.

### Summary of Updates

Simulated data analysis has been augmented to include testing over a range of parameter values. Comparison to RagTag scaffold has been added to simulated data analysis. Biological data analysis now includes patching of the tomato M82 genome in addition to human samples. Automated misjoin detection is now validated using Hi-C data.

