## Supplementary Figures for "GPatch enables chromosome-scale, gap-free pseudoassemblies from fragmented draft genomes"

**Figure S1:** Dot plots comparing GPatch-patched simulated-NA12878 no-indel draft assembly with T2T-CHM13.
**Figure S2:** Dot plots comparing GPatch-patched simulated-HG002 no-indel draft assembly with T2T-CHM13.
**Figure S3:** Patch locations in GPatch-patched simulated genomes.
**Figure S4:** Dot plots comparing RagTag Patch-patched simulated NA12878 no-indel draft assembly with T2T-CHM13.
**Figure S5:** Dot plots comparing RagTag Patch-patched simulated HG002 no-indel draft assembly with T2T-CHM13.
**Figure S6:** Dot plots comparing GPatch-patched simulated NA12878 SURVIVOR draft assembly with T2T-CHM13.
**Figure S7:** Dot plots comparing GPatch-patched simulated HG002 SURVIVOR draft assembly with T2T-CHM13.
**Figure S8:** RagTag Scaffold Performance on simulated NA12878 SURVIVOR draft assembly.
**Figure S9:** GPatch Performance vs. Mapping Quality Threshold.
**Figure S10:** Dot plots comparing initial GPatch NA12878 assembly with T2T-CHM13.
**Figure S11:** Dot plots comparing initial GPatch HG002 assembly with T2T-CHM13.
**Figure S12:** Dot plots comparing initial GPatch NA12878 assembly with T2T-NA12878.
**Figure S13:** Dot plots comparing initial GPatch HG002 assembly with T2T-HG002.
**Figure S14:** Dot plots comparing GPatch M82 pseudoassembly with SL3 reference assembly.
**Figure S15:** Dot plots comparing GPatch M82 pseudoassembly with M82 reference assembly.
**Figure S16:** Dot plots comparing GPatch M82 reference assembly with SL3 reference assembly.
**Figure S17:** Hi-C Validation of NA12878 Misjoins.
**Figure S18:** Dot plots comparing GPatch NA12878 assembly with T2T-CHM13 after one round contig-breaking.
**Figure S19:** Dot plots comparing GPatch NA12878 assembly with T2T-NA12878 after one round contig-breaking.
**Figure S20:** Dot plots comparing GPatch NA12878 assembly with T2T-CHM13 after two rounds contig-breaking.

**Figure S21:** Dot plots comparing GPatch NA12878 assembly with T2T-NA12878 after two rounds contig-breaking.
**Figure S22:** Dot plots comparing GPatch HG002 assembly with T2T-CHM13 after one round contig-breaking.
**Figure S23:** Dot plots comparing GPatch HG002 assembly with T2T-HG002 after one round contig-breaking.
**Figure S24:** NA12878 Hi-C Data Processed with the GPatch NA12878 Pseudoassembly as Reference.
**Figure S25:** NA12878 Hi-C Data Processed with the T2T-CHM13 Assembly as Reference.

#### Fig. S1: Dot plots comparing GPatch-patched simulated-NA12878 no-indel draft assembly with T2T-CHM13

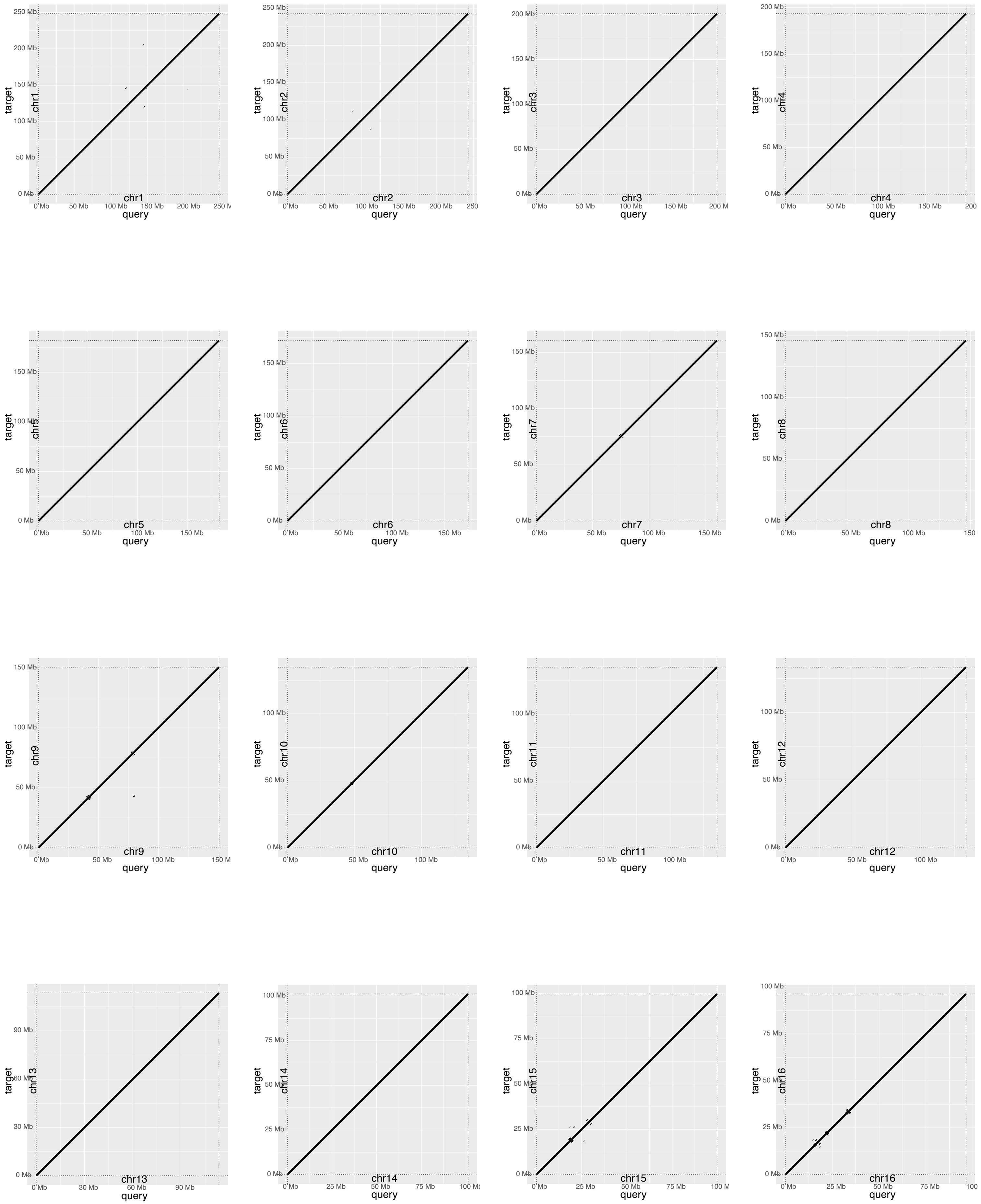


All Panels: target=T2T-CHM13, query=GPatch simulated-NA12878.

#### Fig. S1: Dot plots comparing GPatch-patched simulated-NA12878 no-indel draft assembly with T2T-CHM13

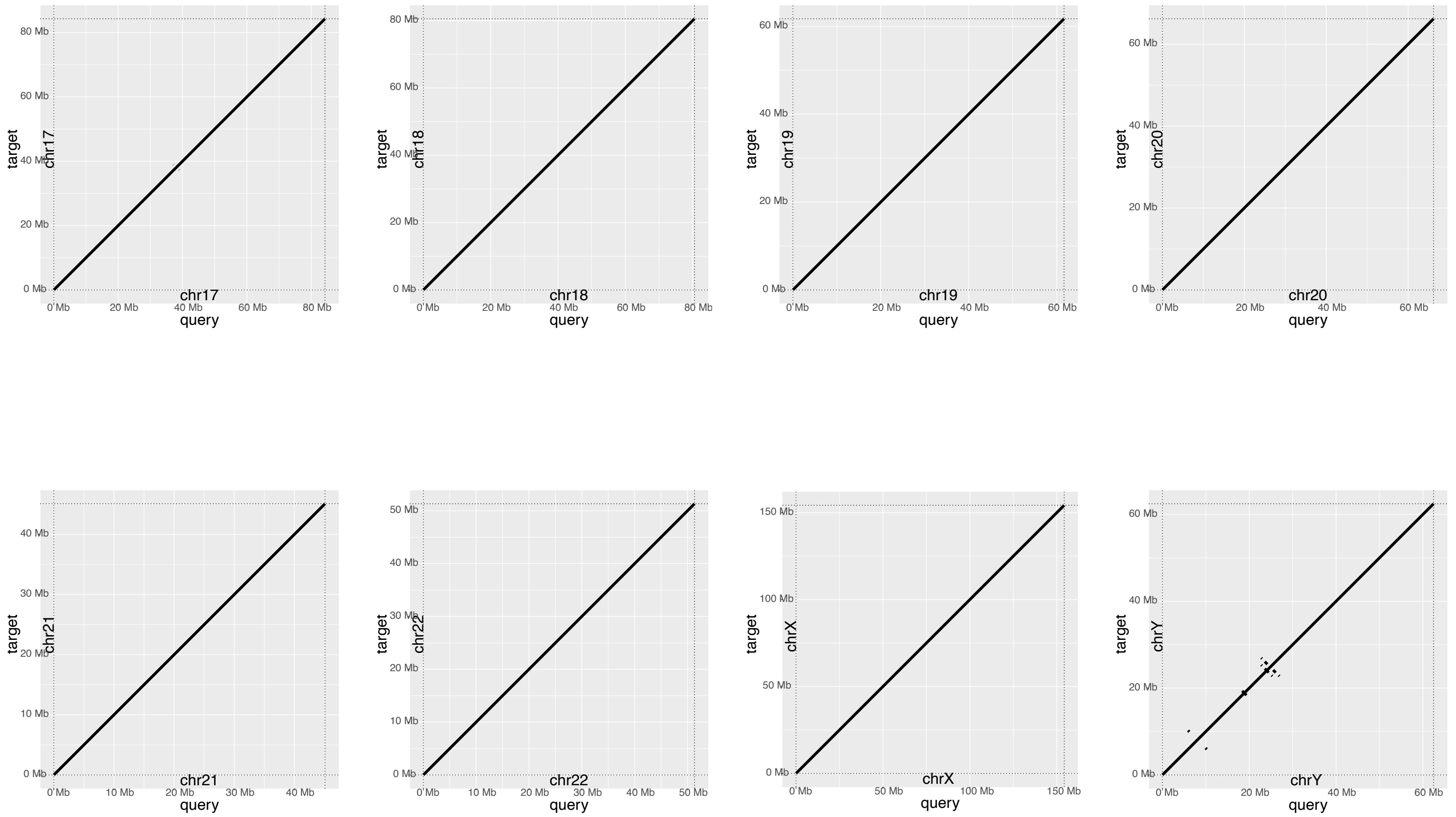


**Figure S1: Dot plots comparing GPatch-patched simulated-NA12878 no-indel draft assembly with T2T-CHM13.** Dot plots compare patched chromosomes constructed with GPatch from simulated NA12878 contigs containing no indels or SNVs. All Panels: target=T2T-CHM13, query=GPatch simulated-NA12878.

**Figure S2: Dot plots comparing GPatch-patched simulated-HG002 no-indel draft assembly with T2T-CHM13.**

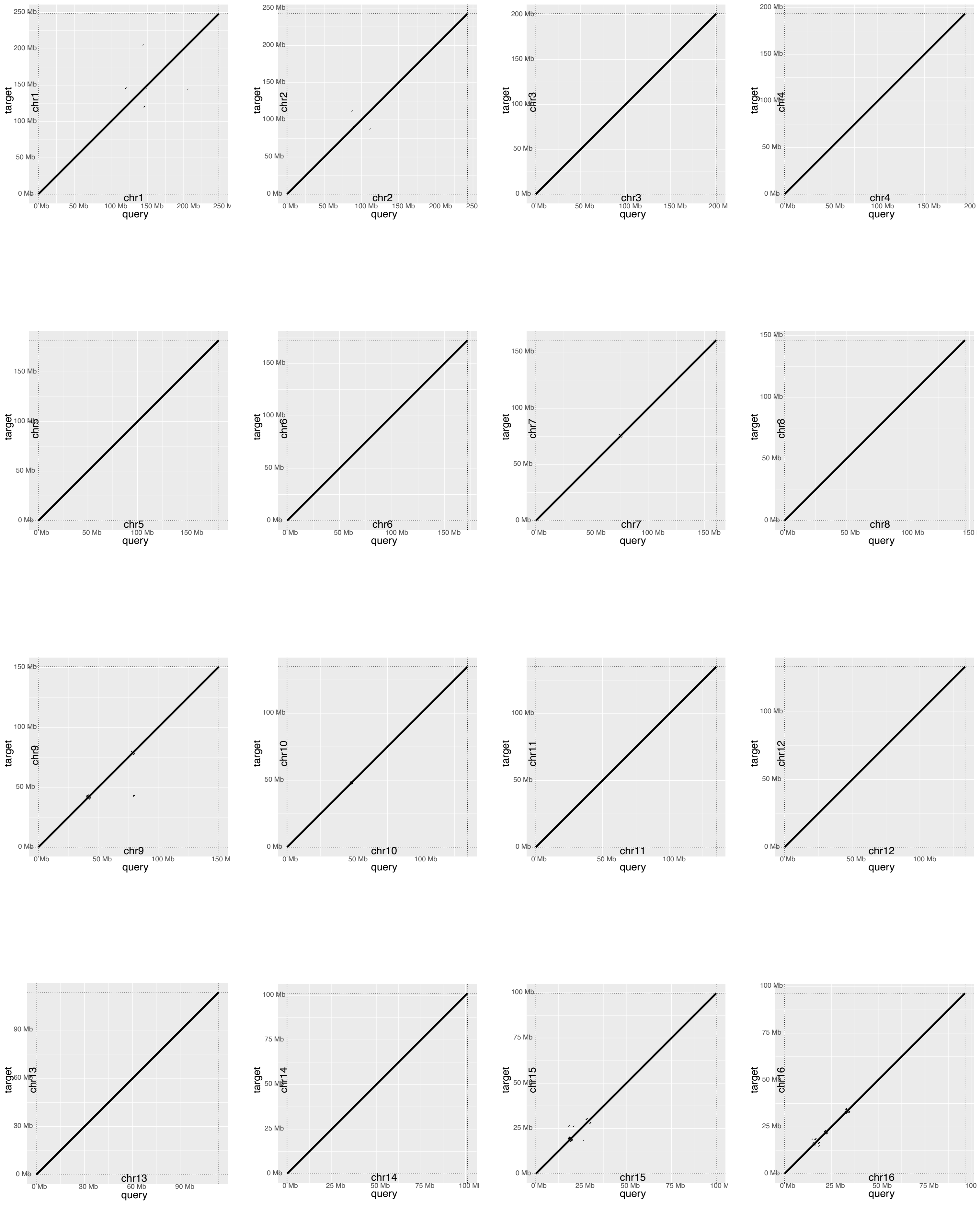


All panels: target=T2T-CHM13, query=GPatch simulated-HG002.

**Figure S2: Dot plots comparing GPatch-patched simulated-HG002 no-indel draft assembly with T2T-CHM13.**

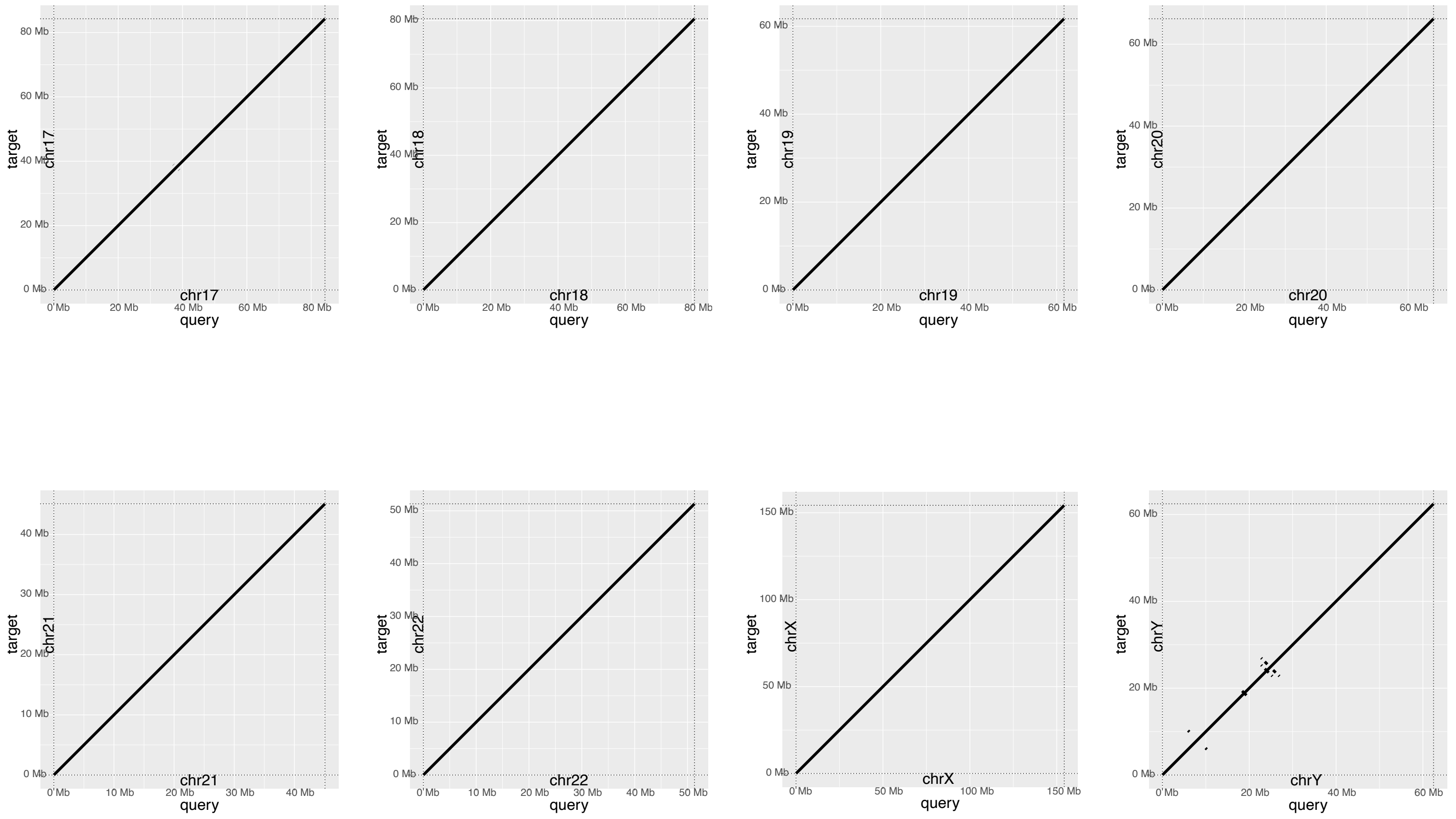


**Figure S2: Dot plots comparing GPatch-patched simulated-HG002 no-indel draft assembly with T2T-CHM13.** Dot plots compare patched chromosomes constructed with GPatch from simulated HG002 contigs containing no indels or SNVs. All panels: target=T2T-CHM13, query=GPatch simulated-HG002.

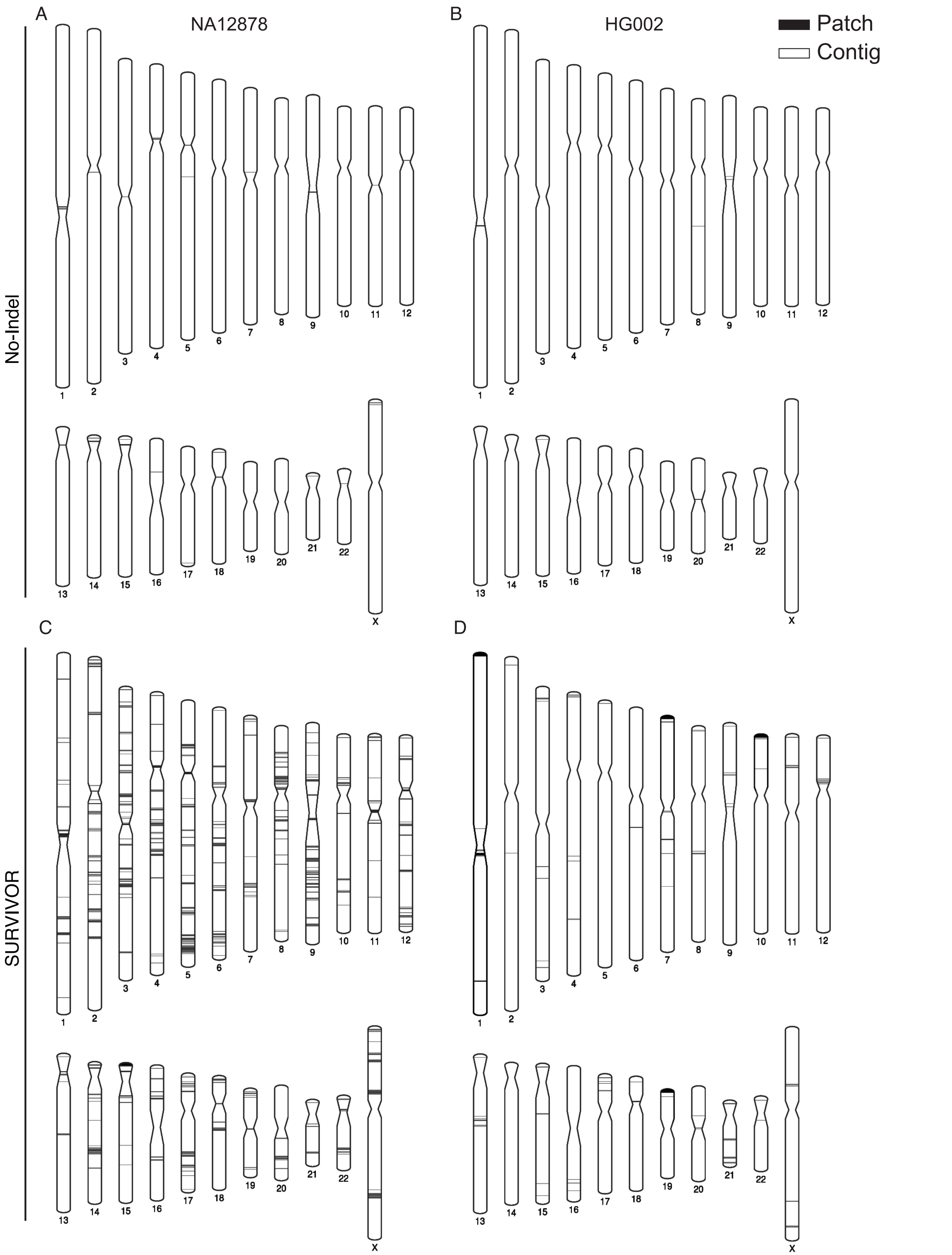


**Figure S3: Patch locations in GPatch-patched simulated genomes.** Ideograms show the locations of T2T-CHM13-derived patches in patched genomes assembled by GPatch from simulated contigs in the following genomes: A) GPatch simulated-NA1278, no-indel set. B)GPatch simulated-HG002, no-indel set. C) GPatch simulated-NA1278, SURVIVOR set. D) GPatch simulated-HG002, SURVIVOR set.

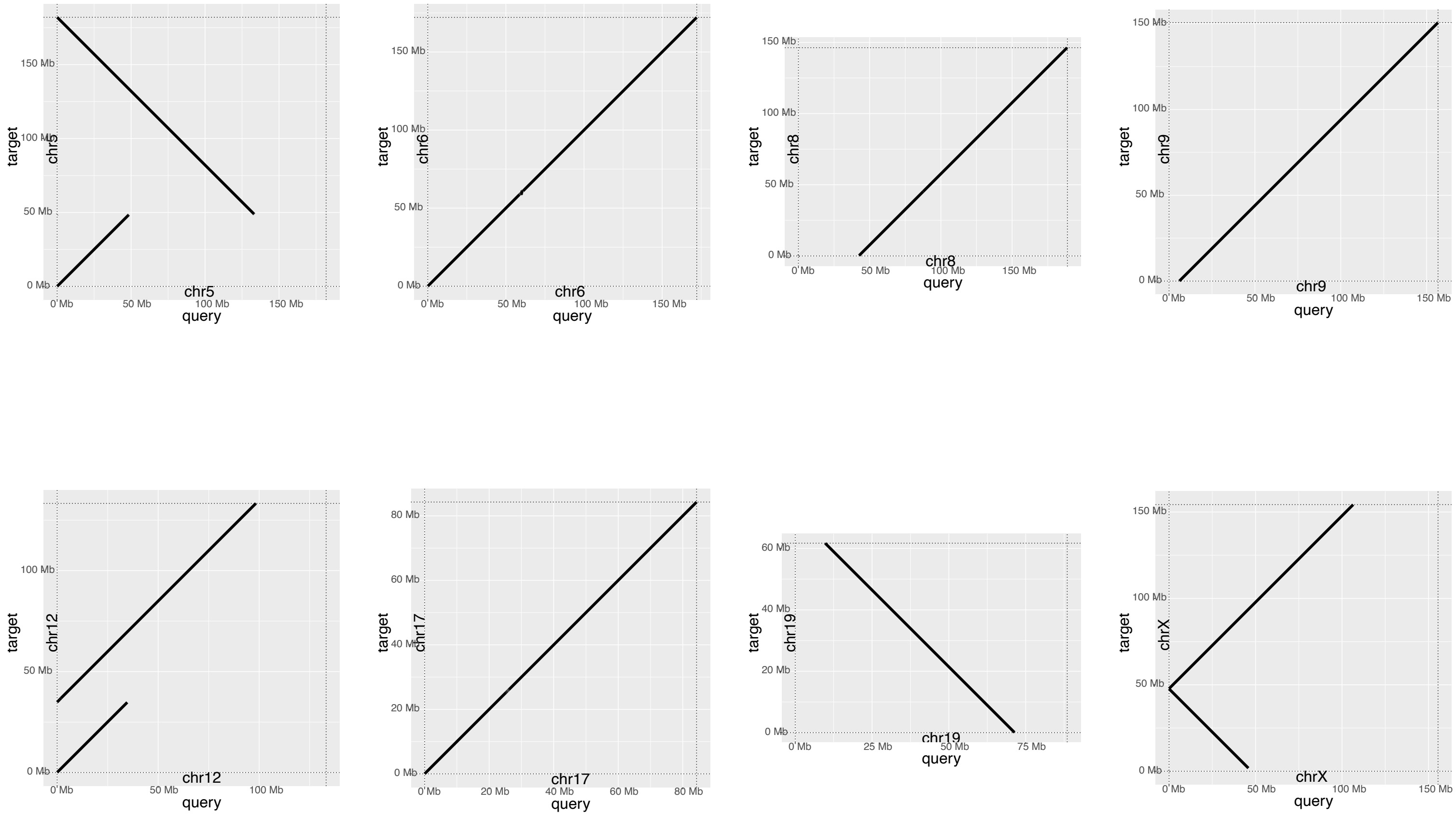


**Figure S4: Dot plots comparing RagTag Patch-patched simulated NA12878 no-indel draft assembly with T2T-CHM13.** Dot plots compare patched chromosomes constructed with RagTag patch from simulated NA12878 contigs containing no indels or SNVs. All Panels: target=T2T-CHM13, query=RagTag Patch NA12878.

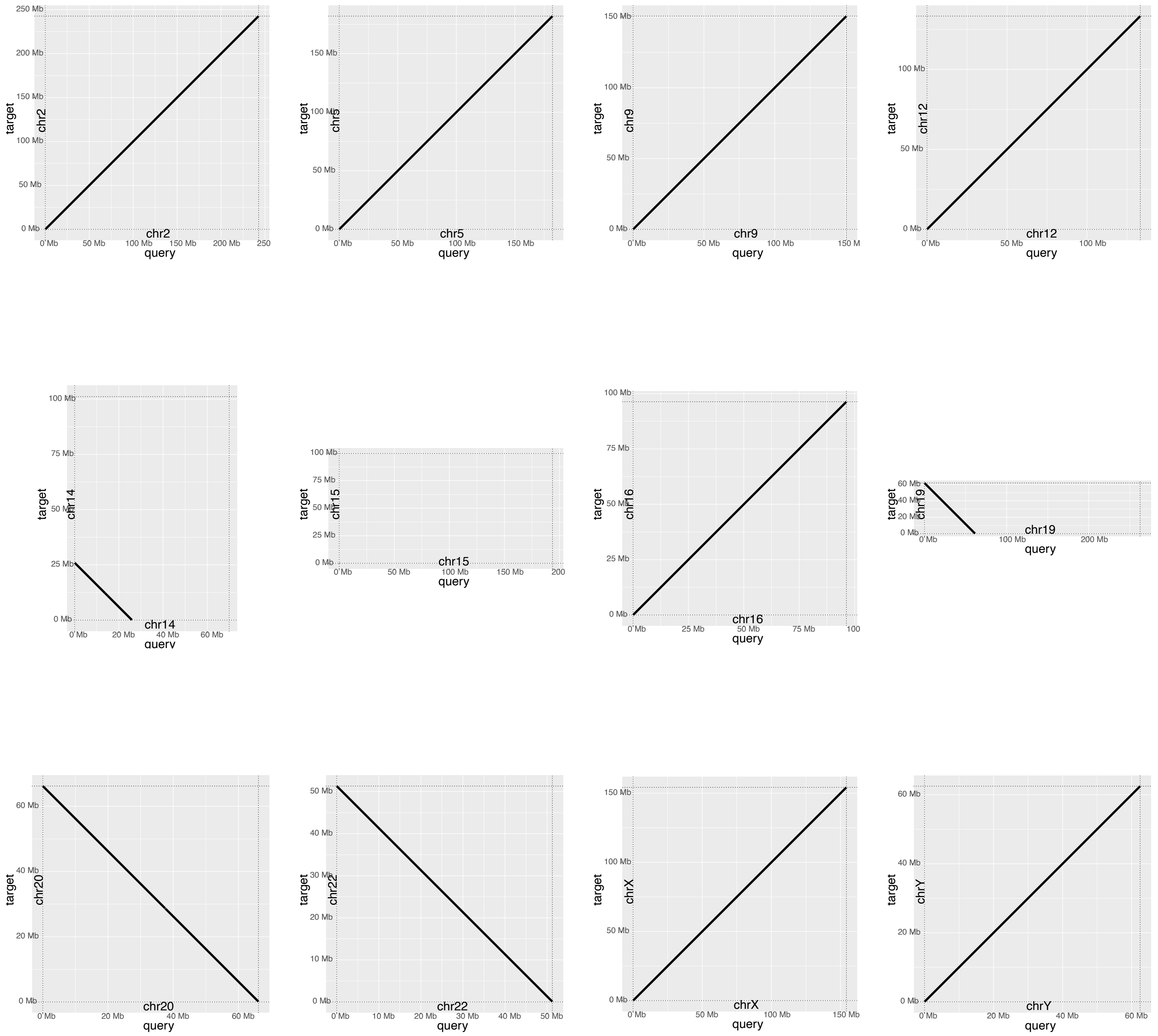


**Figure S5: Dot plots comparing RagTag Patch-patched simulated HG002 no-indel draft assembly with T2T-CHM13.** Dot plots compare patched chromosomes constructed with RagTag patch from simulated HG002 contigs containing no indels or SNVs. All Panels, target=T2T-CHM13, query=RagTag Patch-HG002.

**Figure S6: Dot plots comparing GPatch-patched simulated NA12878 SURVIVOR draft assembly with T2T-CHM13.**

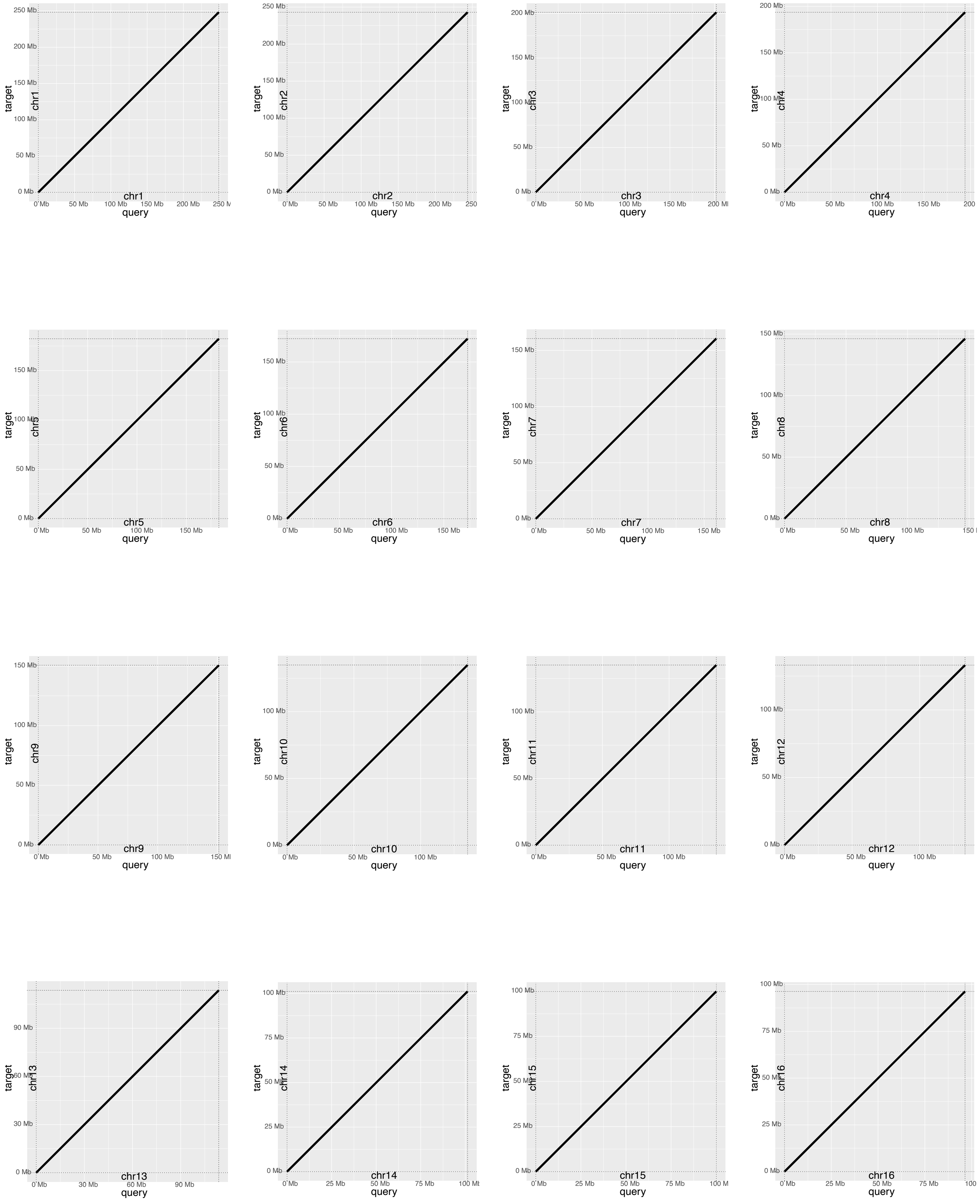


All Panels, target=NA12878-SURVIVOR simulated genome, query=GPatch NA12878-SURVIVOR.

**Figure S6: Dot plots comparing GPatch-patched simulated NA12878 SURVIVOR draft assembly with T2T-CHM13.**

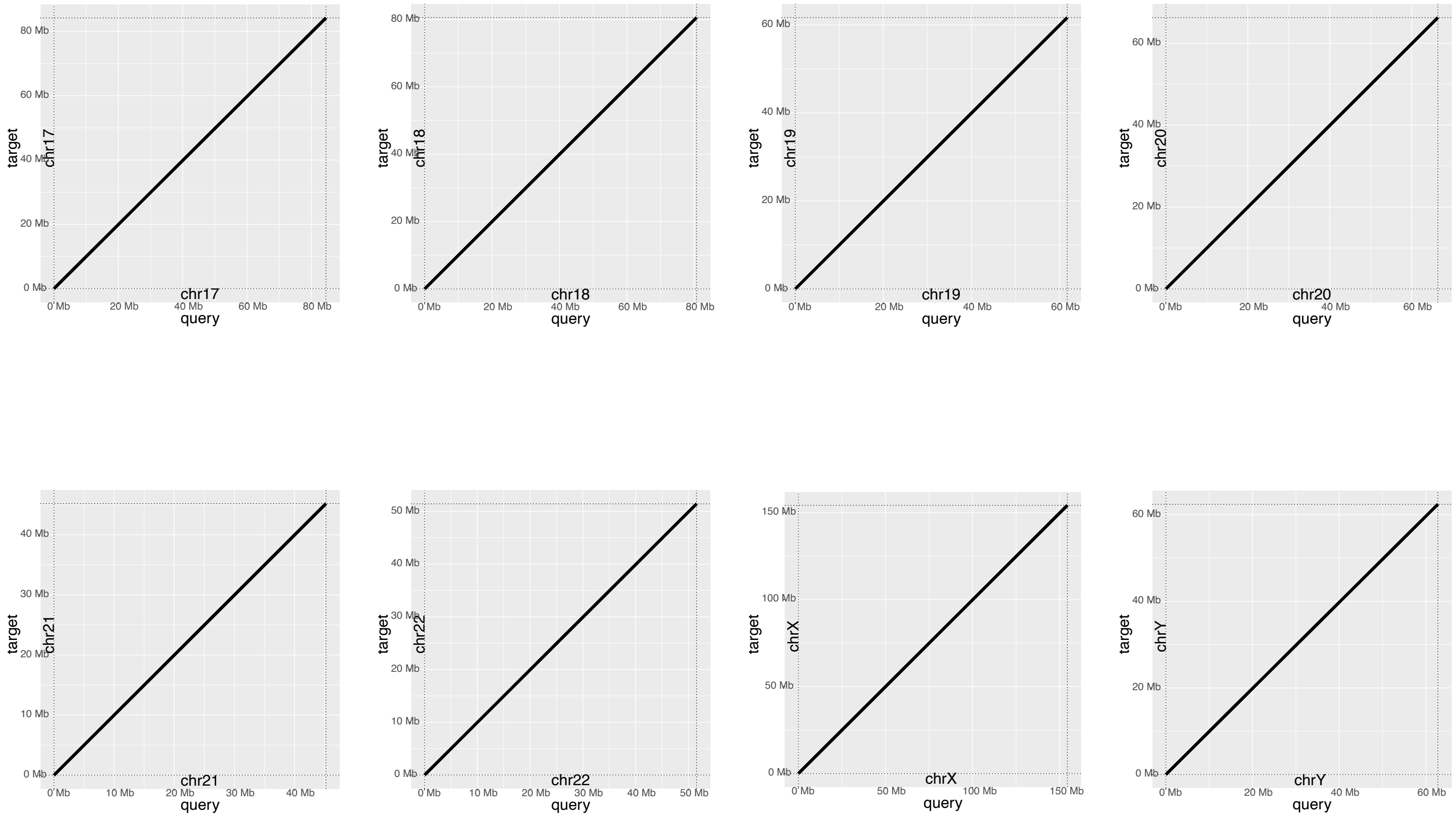


**Figure S6: Dot plots comparing GPatch-patched simulated NA12878 SURVIVOR draft assembly with T2T-CHM13.** Dot plots compare patched chromosomes constructed with GPatch from simulated NA12878 contigs containing 10,000 random indels and a 1% rate of single-nucleotide variants, simulated with SURVIVOR. All Panels, target=NA12878-SURVIVOR simulated genome, query=GPatch NA12878-SURVIVOR.

**Figure S7: Dot plots comparing GPatch-patched simulated HG002 SURVIVOR draft assembly with T2T-CHM13.**

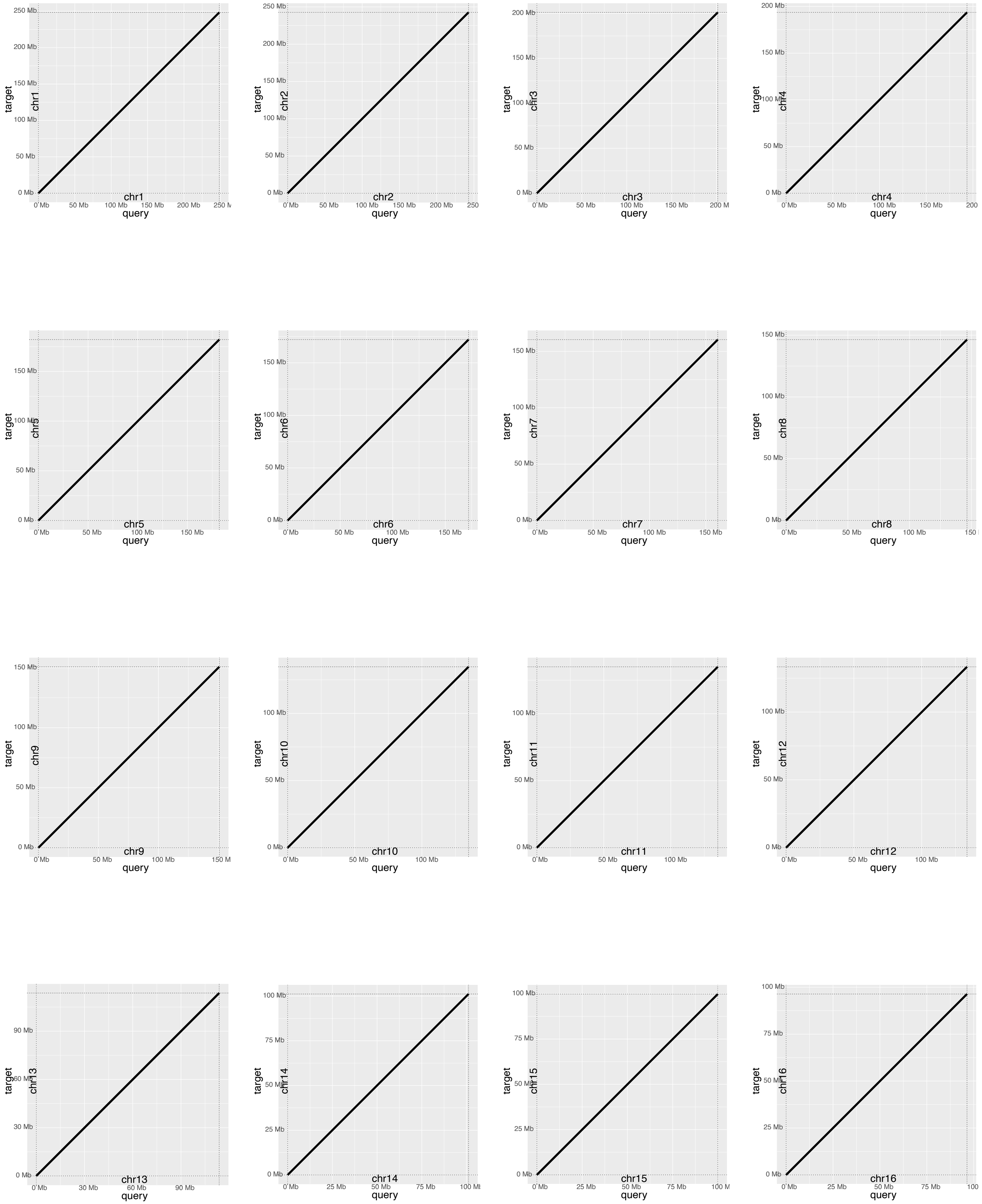


All Panels, target=HG002-SURVIVOR simulated genome, query=GPatch HG002-SURVIVOR.

**Figure S7: Dot plots comparing GPatch-patched simulated HG002 SURVIVOR draft assembly with T2T-CHM13.**

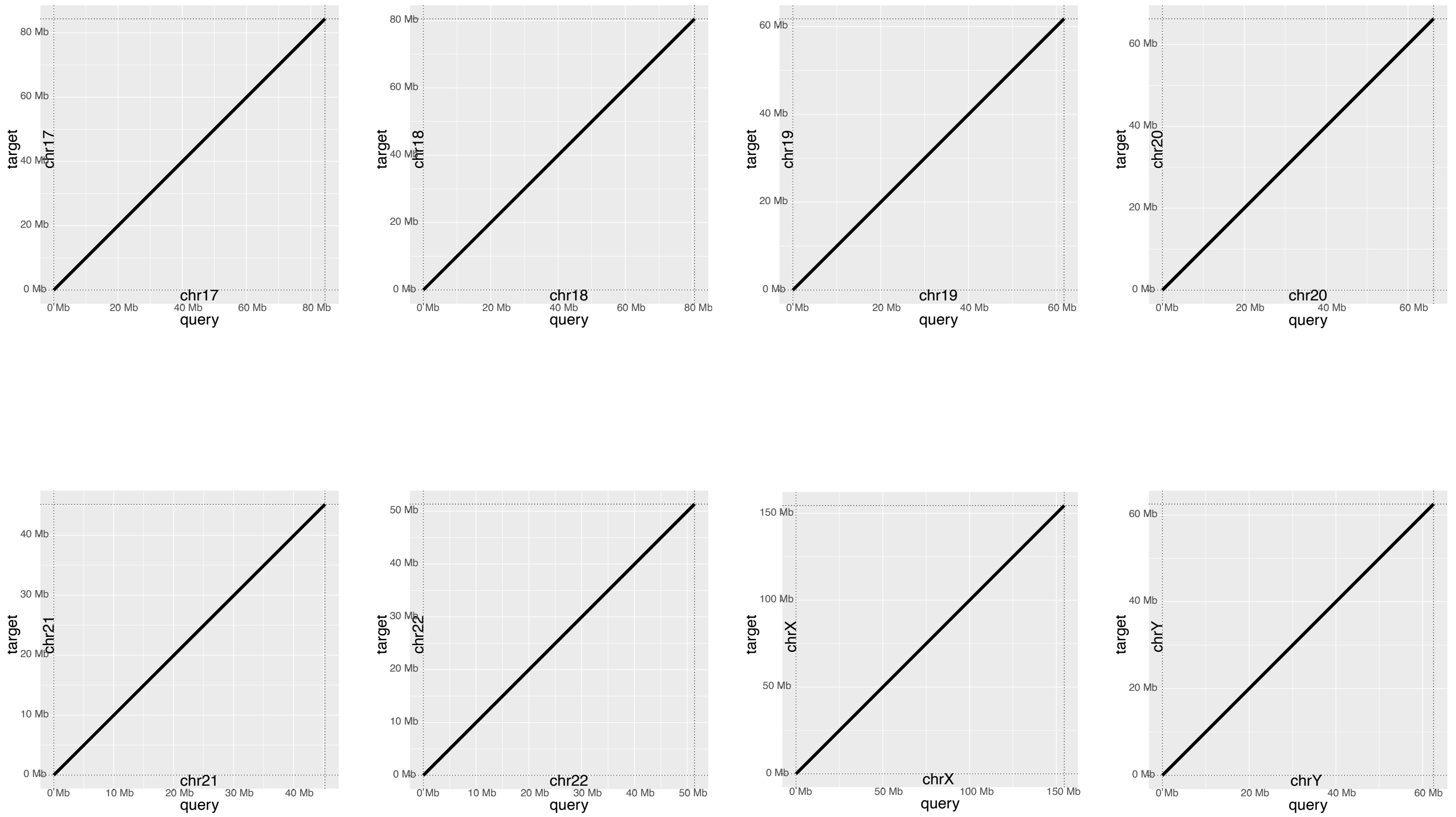


**Figure S7: Dot plots comparing GPatch-patched simulated HG002 SURVIVOR draft assembly with T2T-CHM13.** Dot plots compare patched chromosomes constructed with GPatch from simulated HG002 contigs containing 5,000 random indels and a 1% rate of single-nucleotide variants, simulated with SURVIVOR. All Panels, target=HG002-SURVIVOR simulated genome, query=GPatch HG002-SURVIVOR.

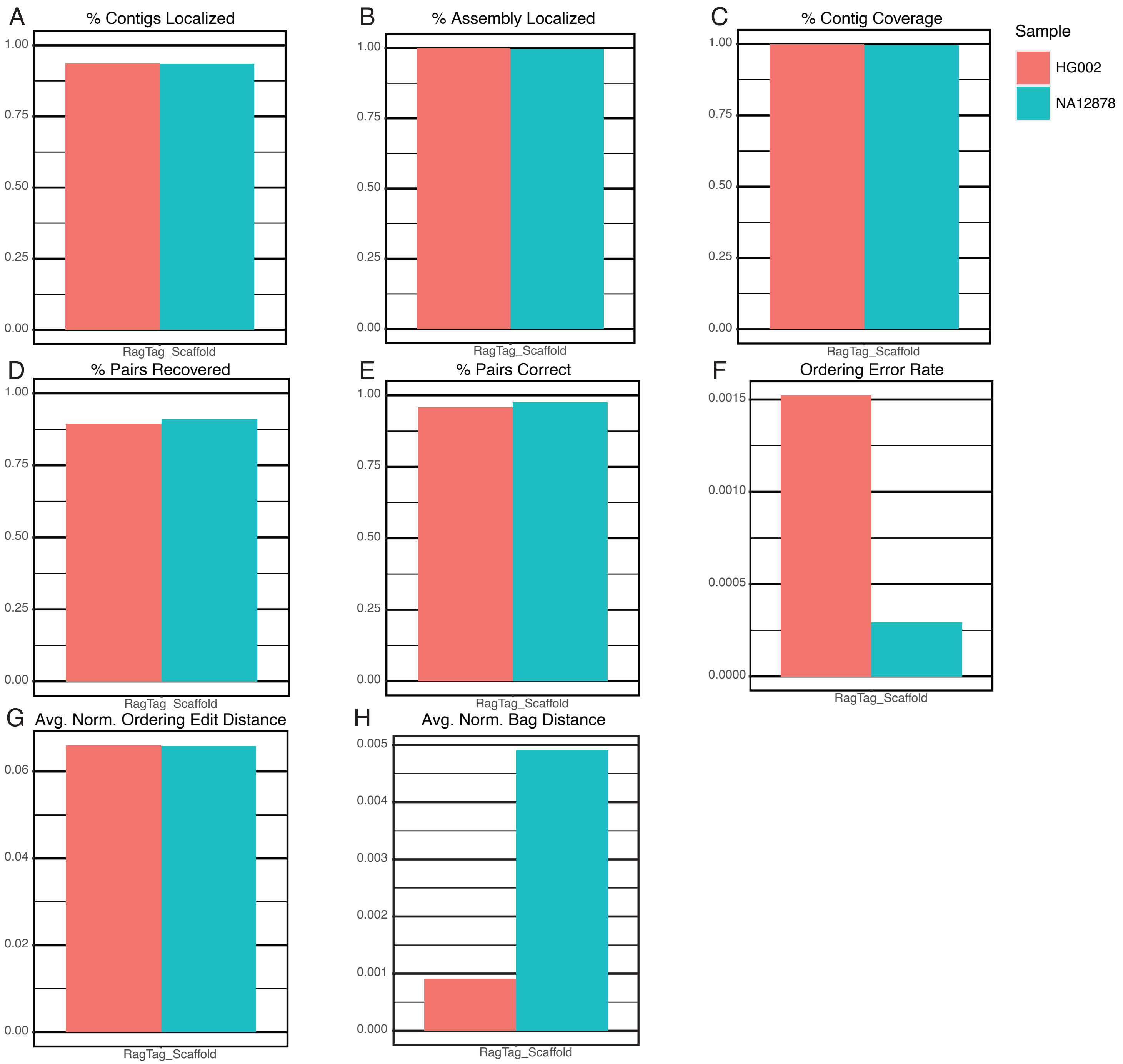


**Figure S8: RagTag Scaffold Performance on Simulated Data with Indels.** A) The percentage of contigs from the source assembly localized in the patched pseudoassembly. B) The percentage of nucleotides from the source assembly localized in the patched assembly. C) The percentage of nucleotides in the patched assembly contributed by contigs. D) The fraction of adjacent contig pairs from the target genome recovered in the patched genome. E) The fraction of adjacent pairs in the patched genome that are also found in the target genome. F) The fraction of contigs placed out-of-order relative to neighboring contigs in the patched sequence. G) Average Levenshtein distance between the observed and actual contig ordering vectors divided by the length of the target sequence. H) Average bag distance between patched and target chromosome sequences divided by target sequence length.

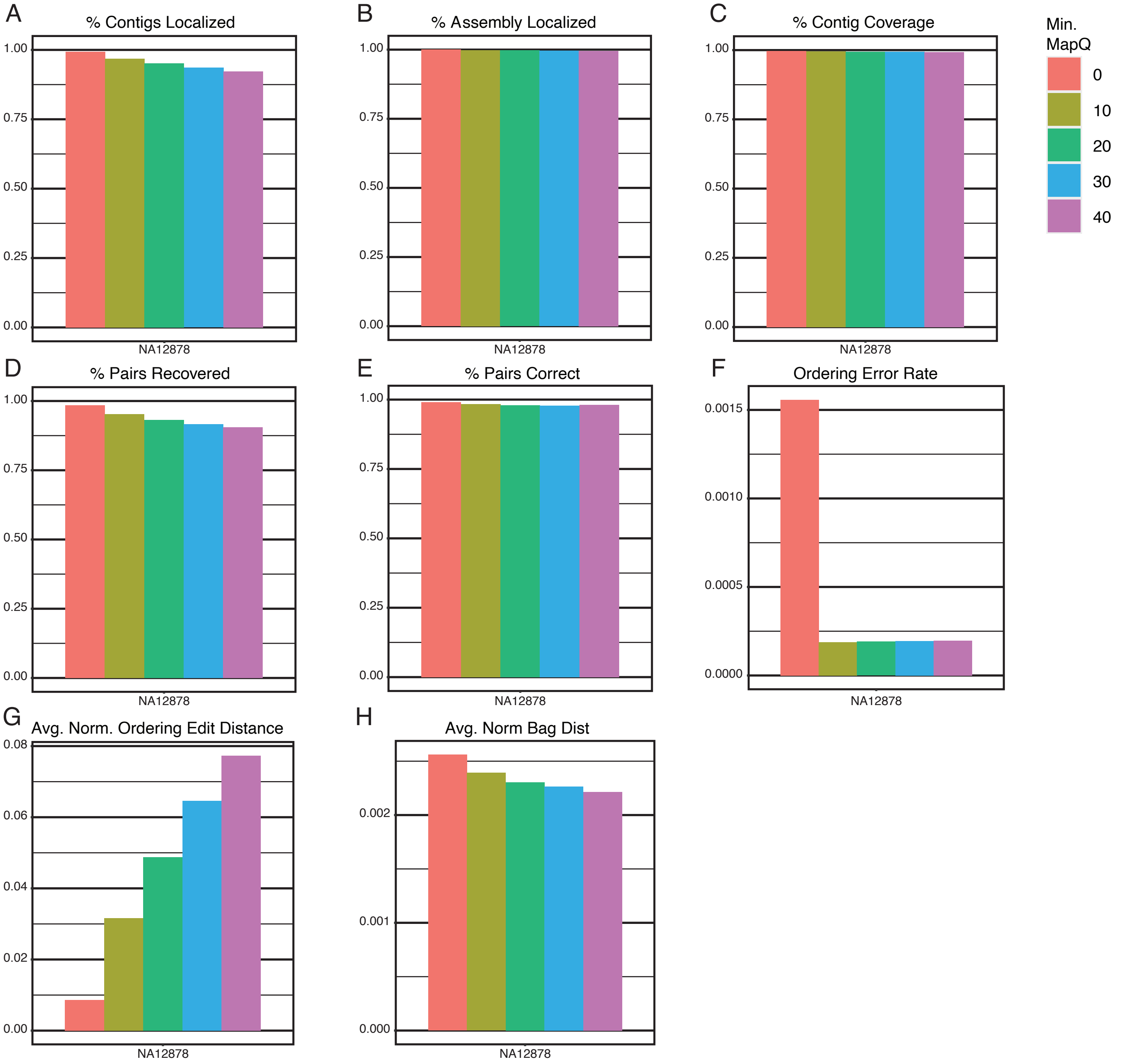


**Figure S9: GPatch Performance vs. Mapping Quality Threshold.** A) The percentage of contigs from the source assembly localized in the patched pseudoassembly. B) The percentage of nucleotides from the source assembly localized in the patched assembly. C) The percentage of nucleotides in the patched assembly contributed by contigs. D) The fraction of adjacent contig pairs from the target genome recovered in the patched genome. E) The fraction of adjacent pairs in the patched genome that are also found in the target genome. F) The fraction of contigs placed out-of-order relative to neighboring contigs in the patched sequence. G) Average Levenshtein distance between the observed and actual contig ordering vectors divided by the length of the target sequence. H) Average bag distance between patched and target chromosome sequences divided by target sequence length.

### Figure S10: Dot plots comparing initial GPatch NA12878 assembly with T2T-CHM13.

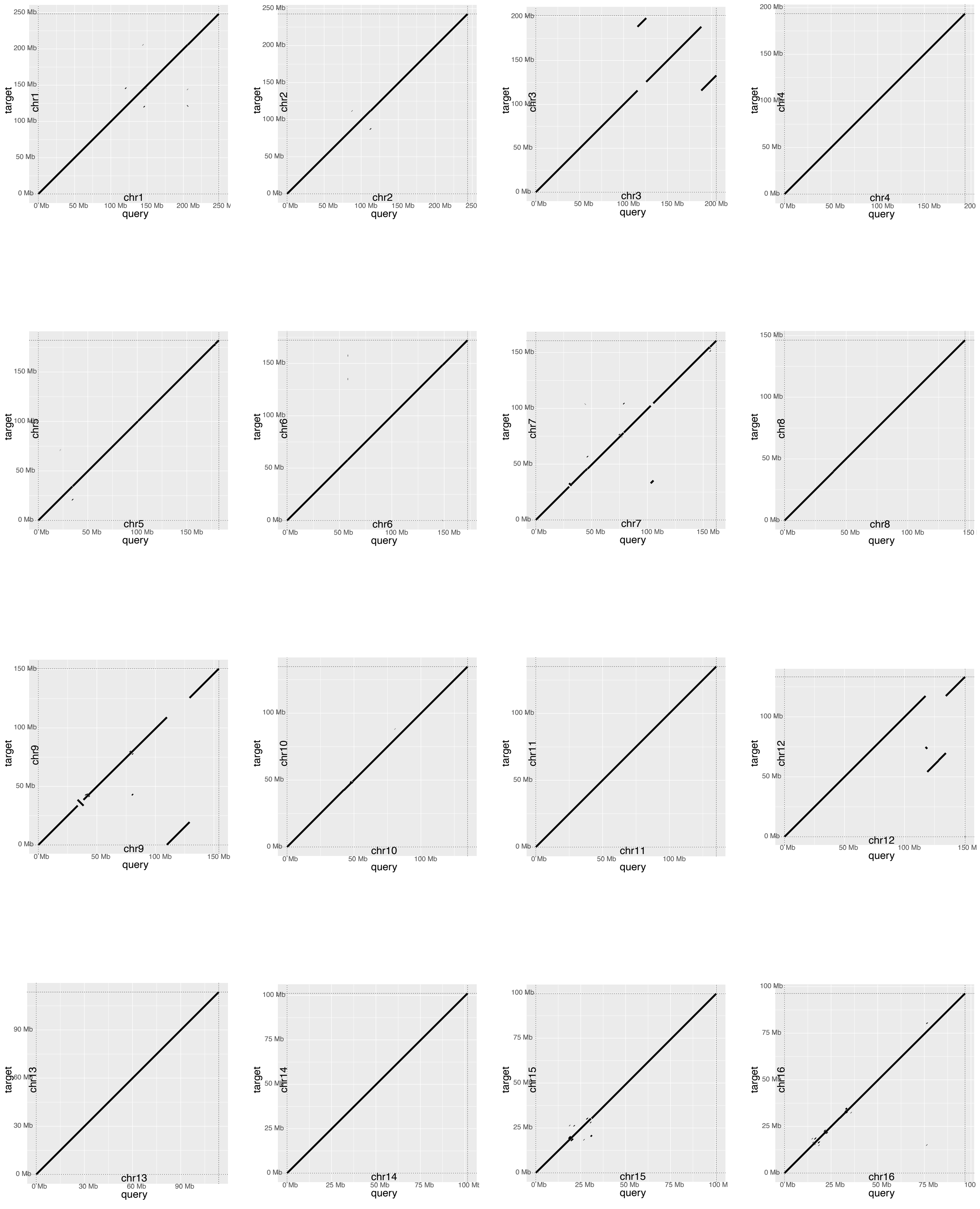


All Panels, target=T2T-CHM13 genome, query=patched-HGSVC-NA12878.

**Figure S10: Dot plots comparing initial GPatch NA12878 assembly with T2T-CHM13.**

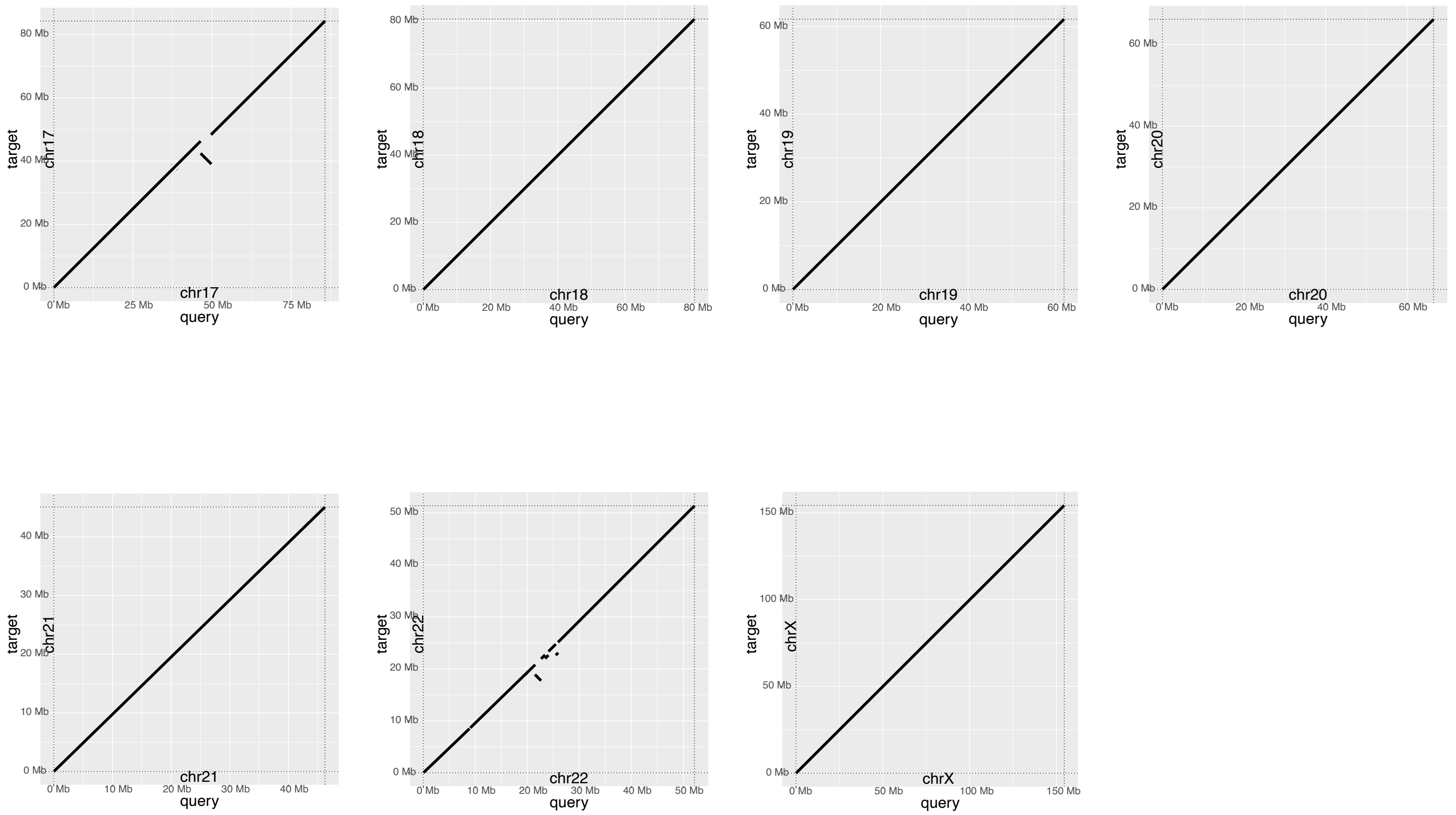


**Figure S10: Dot plots comparing initial GPatch NA12878 assembly with T2T-CHM13.** Dot plots compare patched chromosomes constructed with GPatch from HGSVC NA1878 maternal contigs, plotted against the T2T-CHM13 reference genome used for patching. All Panels, target=T2T-CHM13 genome, query=patched-HGSVC-NA12878.

**Figure S11: Dot plots comparing initial GPatch HG002 assembly with T2T-CHM13**

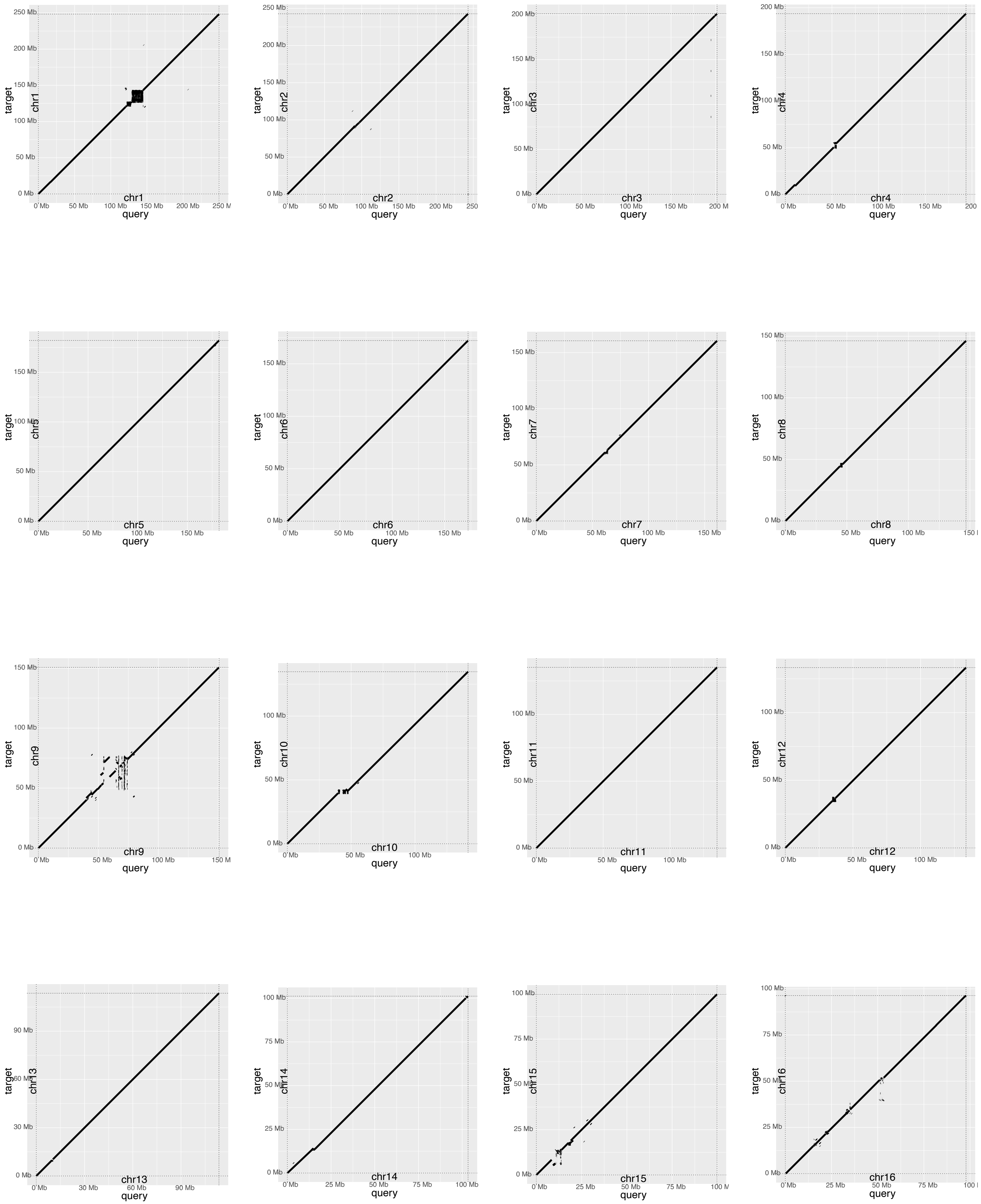


All Panels, target=T2T-CHM13, query=GPatch HG002.

#### Figure S11: Dot plots comparing initial GPatch HG002 assembly with T2T-CHM13

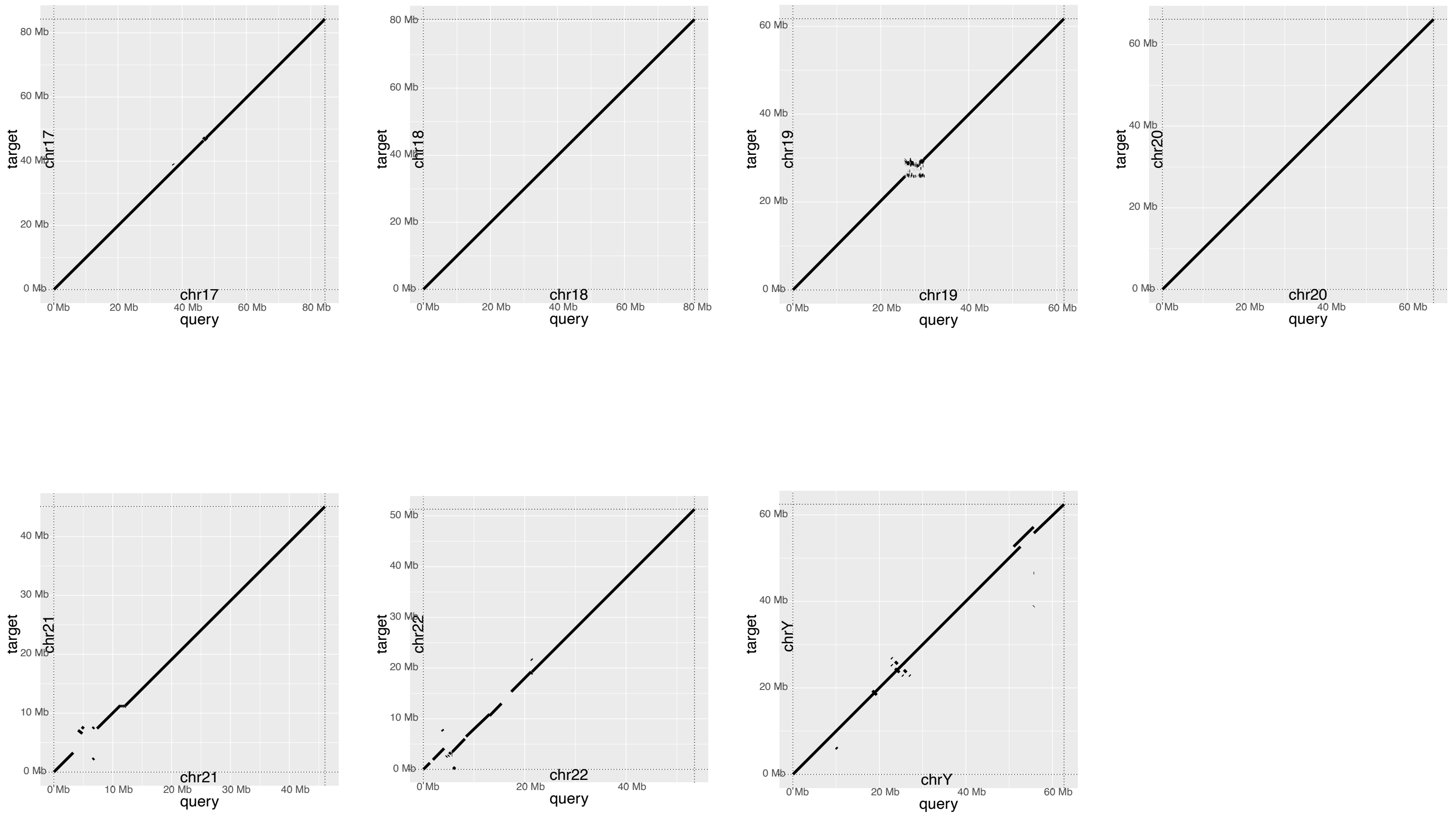


**Figure S11: Dot plots comparing initial GPatch HG002 assembly with T2T-CHM13.** Dot plots compare patched chromosomes constructed with GPatch from HPRC HG002 paternal contigs plotted against the T2T-CHM13 genome assembly. All Panels, target=T2T-CHM13, query=GPatch HG002.

**Figure S12: Dot plots comparing initial GPatch NA12878 assembly with T2T-NA12878.**

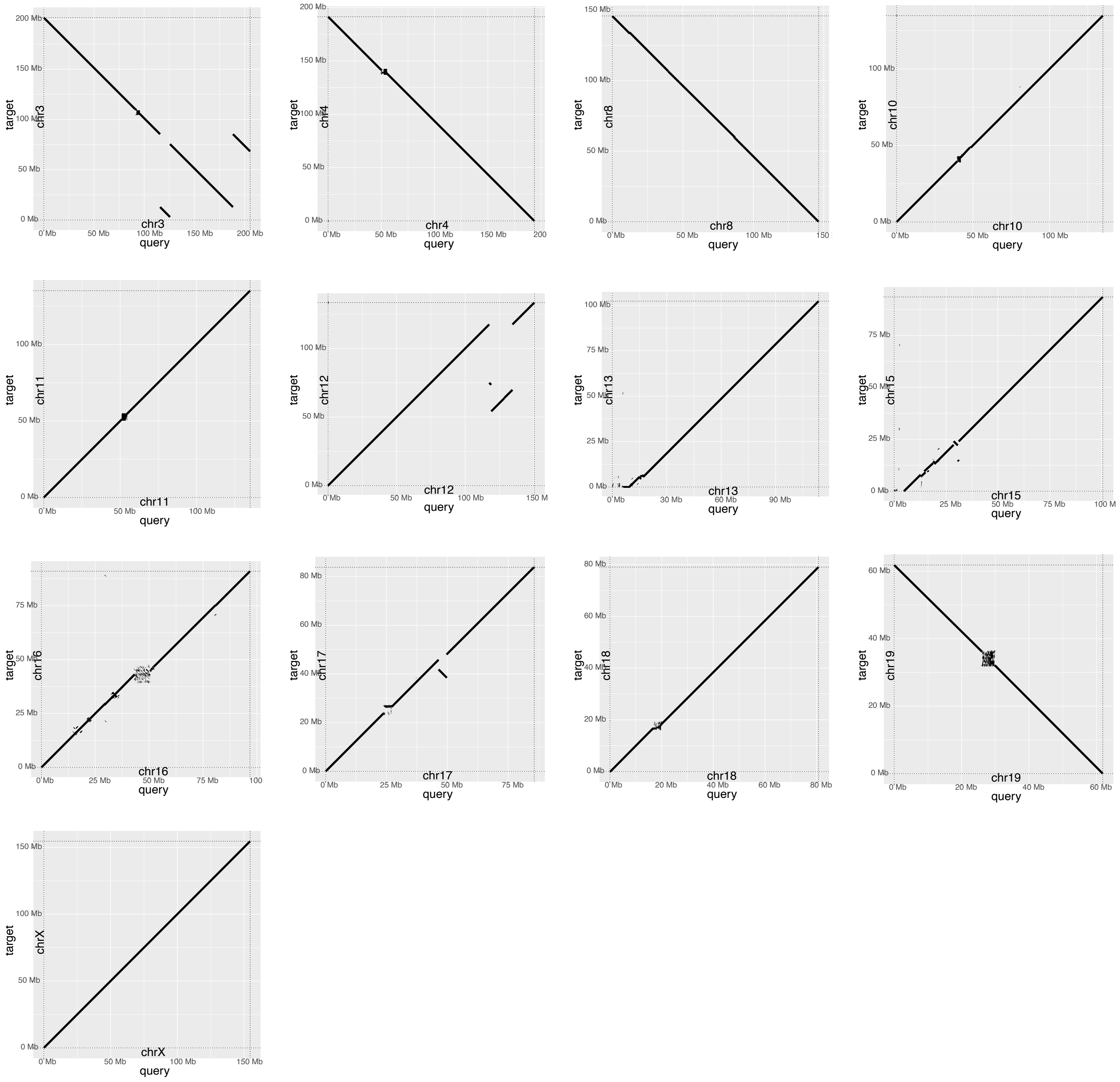


**Figure S12: Dot plots comparing initial GPatch NA12878 assembly with T2T-NA12878.** Dot plots compare patched chromo-somes constructed with GPatch from HGSVC NA1878 maternal contigs, plotted against complete chromosomes from a recently-published chromosome-scale NA12878 genome assembly. All panels, target=T2T-CHM13, query=GPatch NA12878.

#### Figure S13 Dot plots comparing initial GPatch HG002 assembly with T2T-HG002

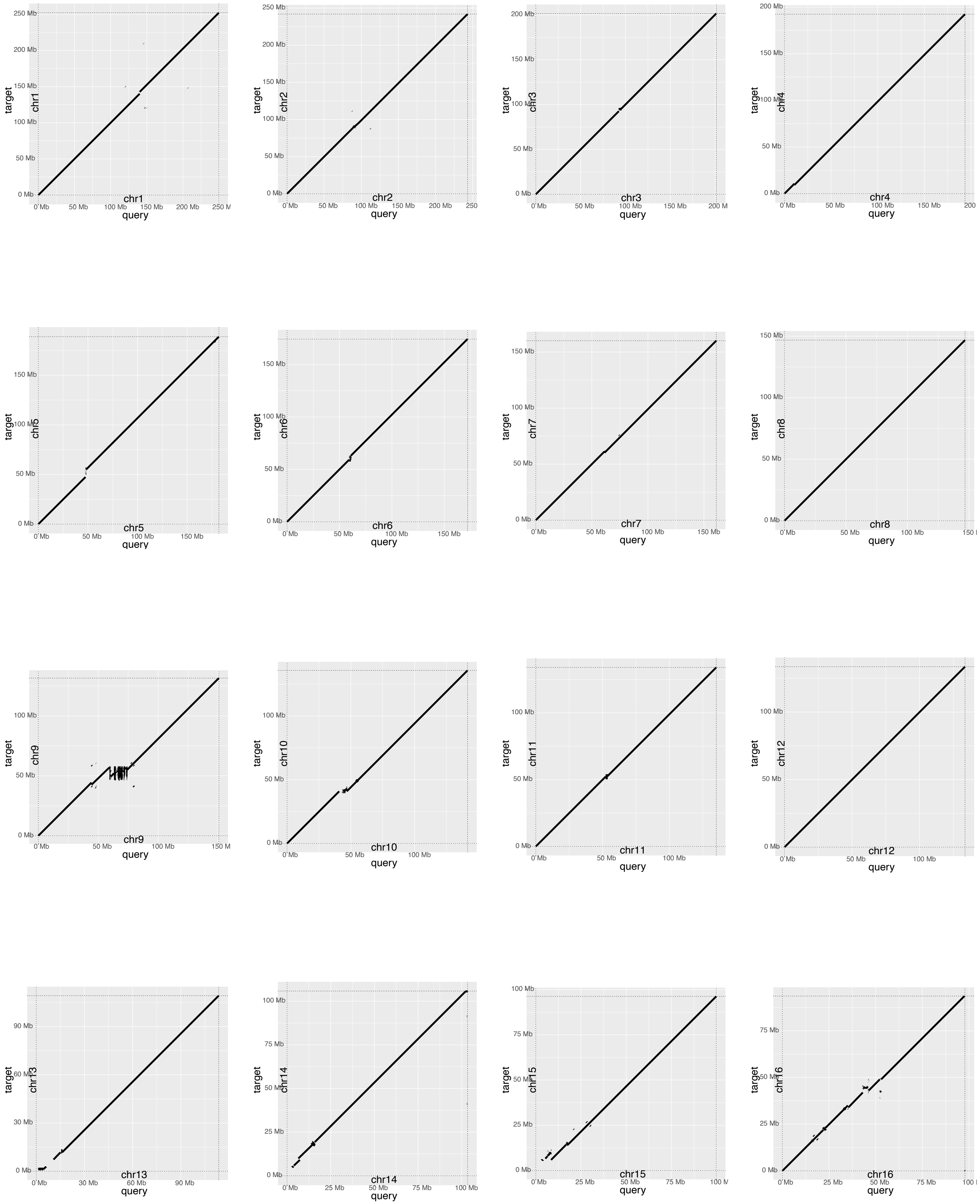


All Panels, target=T2T-HG002, query=patched-GPatch HG002.

**Figure S13: Dot plots comparing initial GPatch HG002 assembly with T2T-HG002**

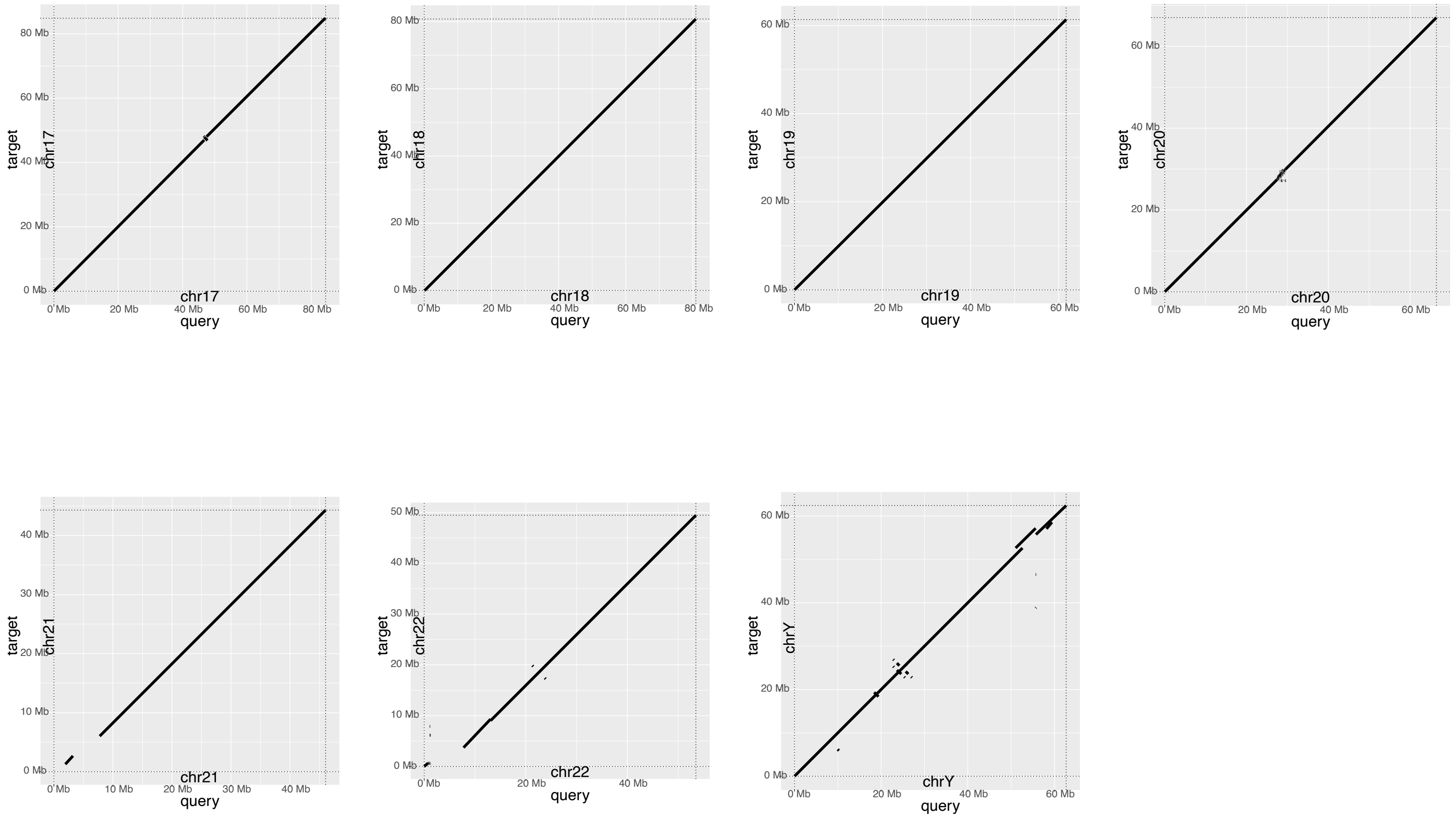


**Figure S13: Dot plots comparing initial GPatch HG002 assembly with T2T-HG002.** Dot plots compare patched chromosomes constructed with GPatch from HPRC HG002 paternal contigs plotted against the T2T-HG002 genome assembly. All Panels, target=T2T-HG002, query=patched-GPatch HG002.

#### Fig. S14: Dot plots comparing GPatch-patched M82 pseudoassembly with SL3 reference

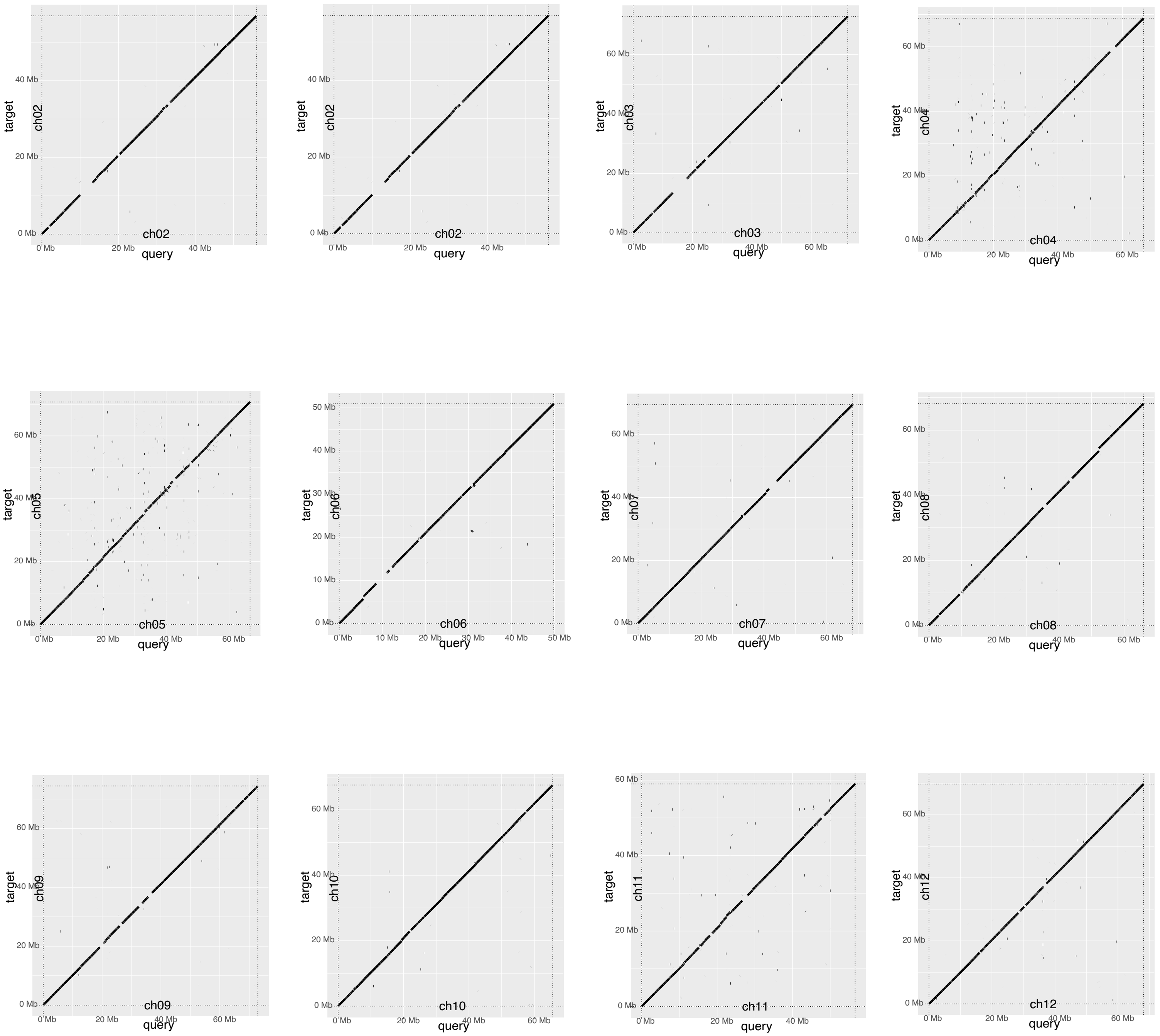


All Panels: target=Tomato SL3 reference assembly, query=GPatch M82 pseudoassembly.

### Fig. S15: Dot plots comparing GPatch-patched M82 pseudoassembly with M82 reference

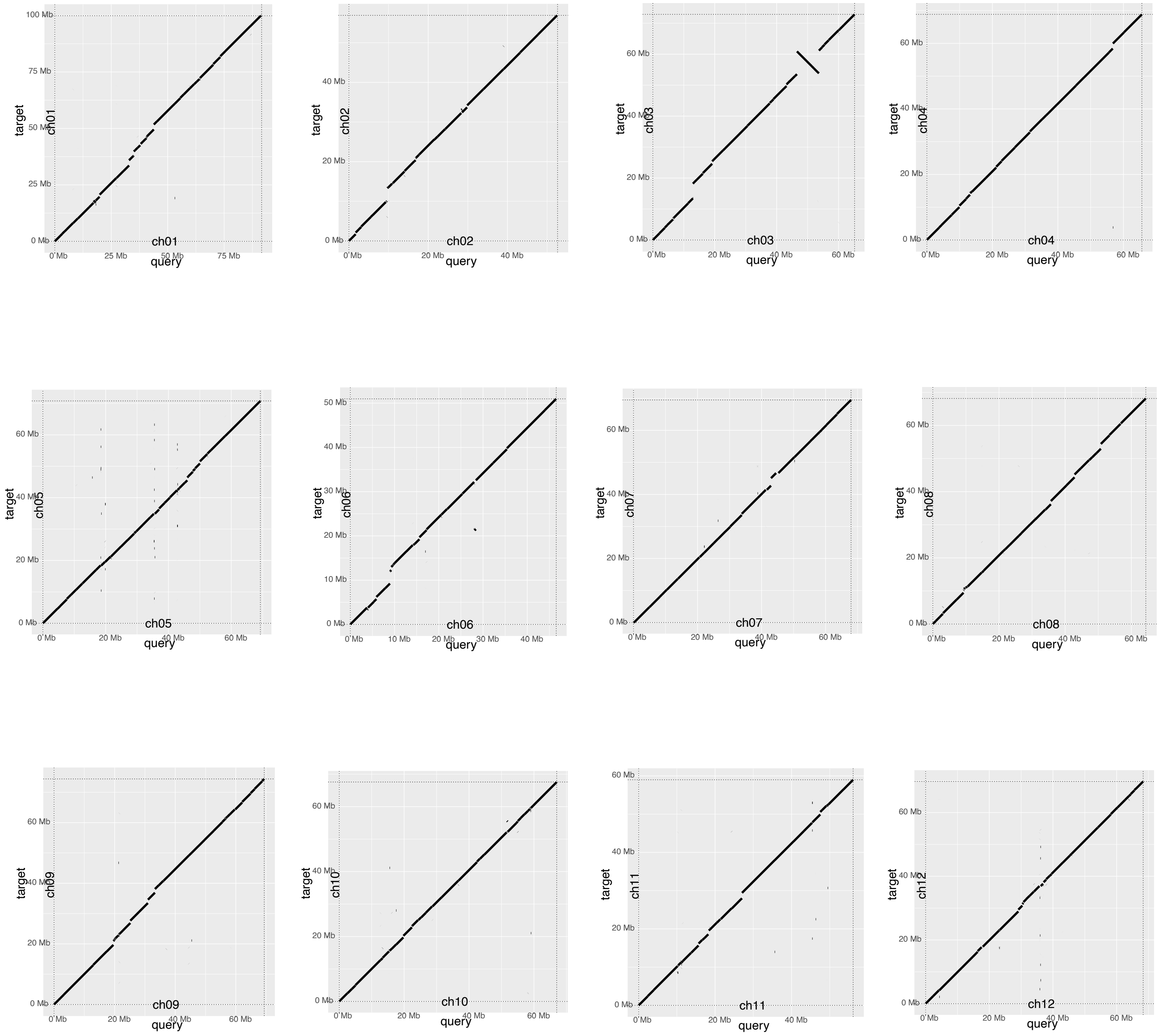


All Panels: target=Tomato M82 reference assembly, query=GPatch M82 pseudoassembly.

**Fig. S16: Dot plots comparing M82 reference assembly with SL3 reference**

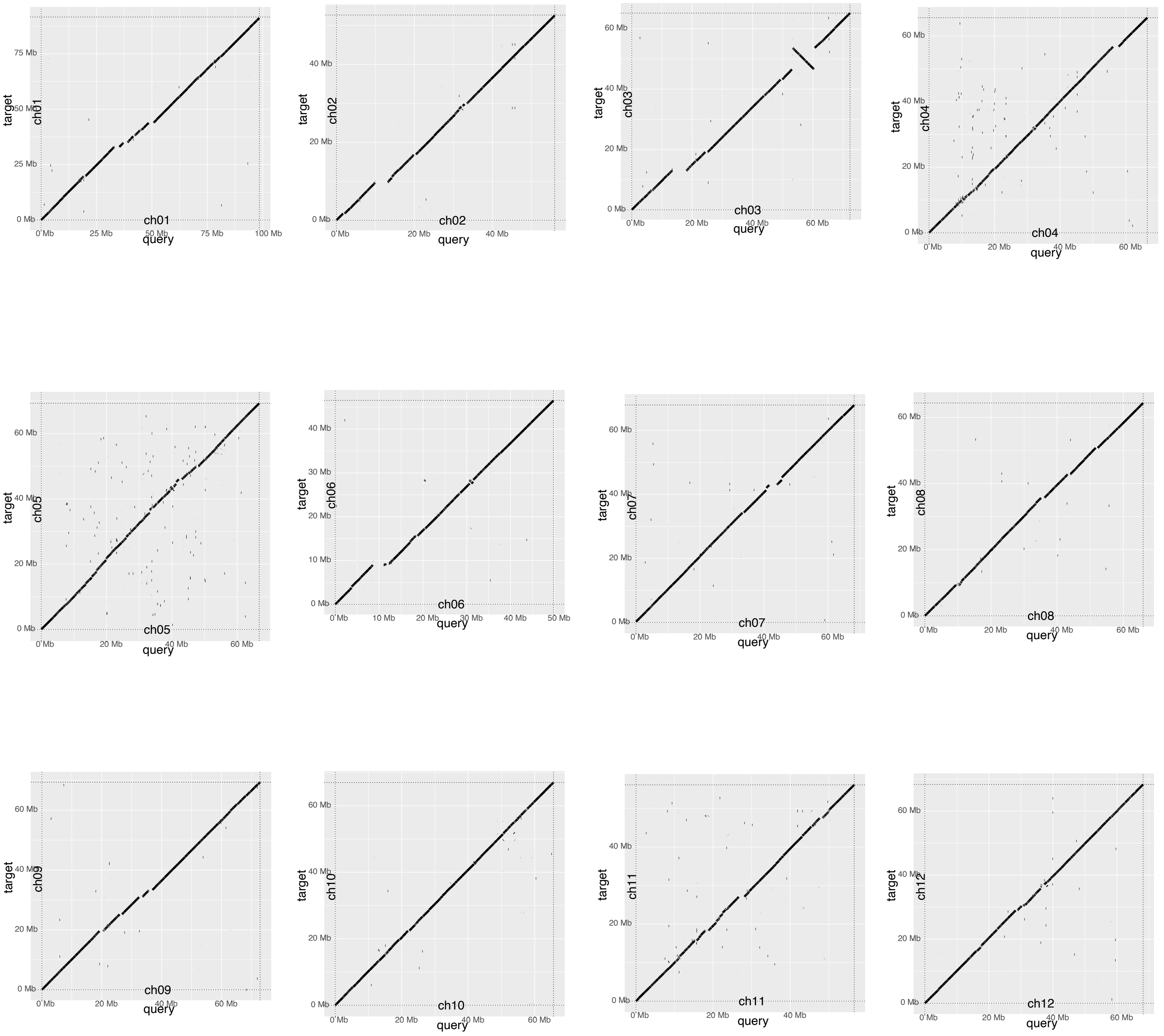


All Panels: target=Tomato SL3 reference assembly, query=M82 reference assembly.

**Fig. S17: Hi-C Validation of NA12878 Misjoins**

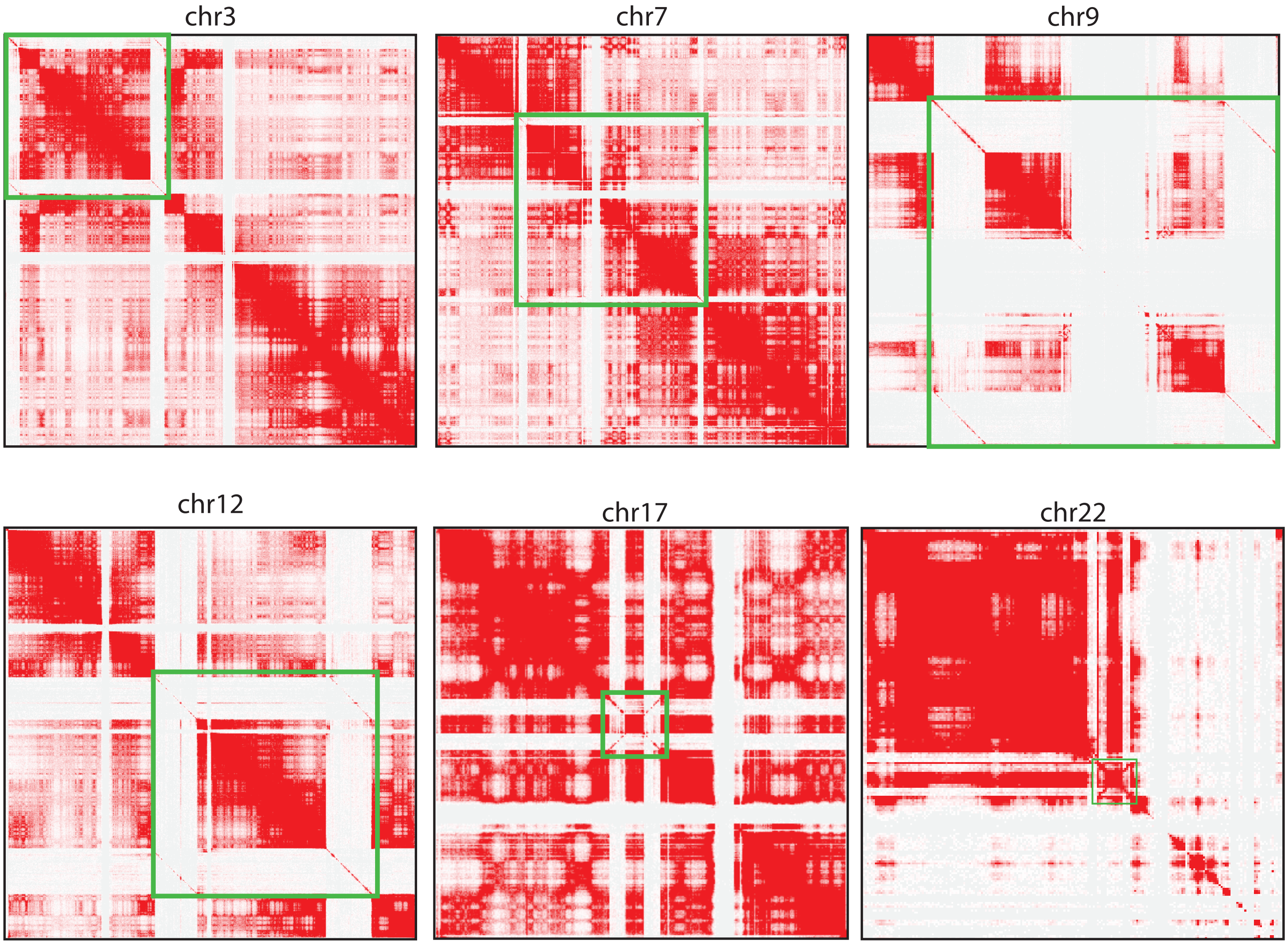


**Figure S17: Hi-C Verification of NA12878 misjoins.** Hi-C data for NA12878 were mapped to the initial patched NA12878 pseudoassembly to verify the presence of putative misjoins observed on chromosomes 3, 7, 9, 12, 17, and 22. Confirmed misjoins are outlined in green.

#### Figure S18: Dot plots comparing GPatch NA12878 assembly with T2T-CHM13 after one round contig-breaking

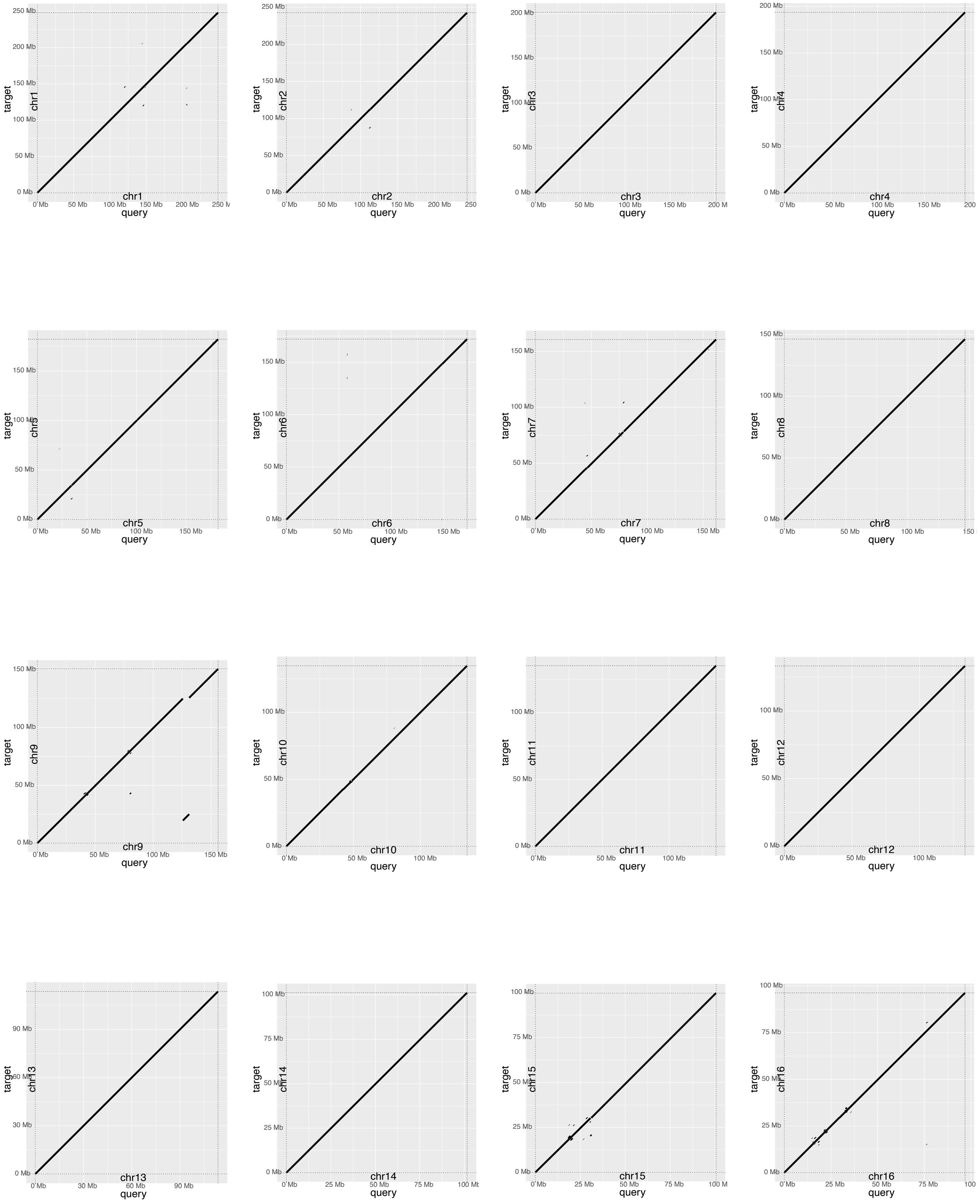


All Panels, target=T2T-CHM13, query=GPatch NA12878 after one round of contig breaking.

### Figure S18: Dot plots comparing GPatch NA12878 assembly with T2T-CHM13 after one round contig-breaking

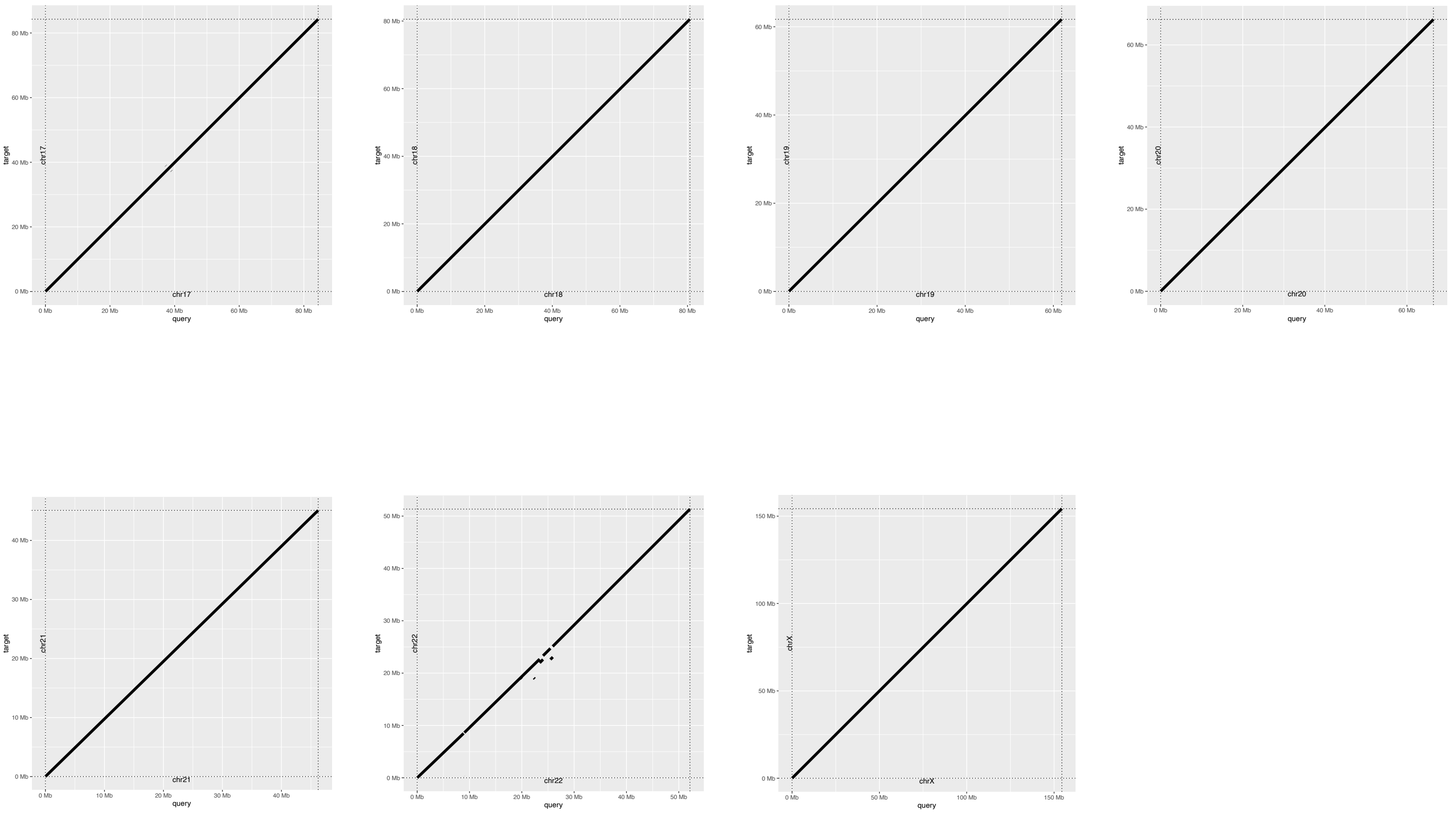


**Figure S18: Dot plots comparing GPatch NA12878 assembly with T2T-CHM13 after one round contig-breaking.** Dot plots compare patched chromosomes constructed with GPatch from NA1878 maternal contigs, after a single round of contig-breaking at the breakpoints of rearrangements >= 1MB in length, plotted against the T2T-CHM13 reference genome used for patching. All Panels, target=T2T-CHM13, query=GPatch NA12878 after one round of contig breaking.

#### Figure S19: Dot plots comparing GPatch NA12878 assembly with T2T-NA12878 after one round contig-breaking

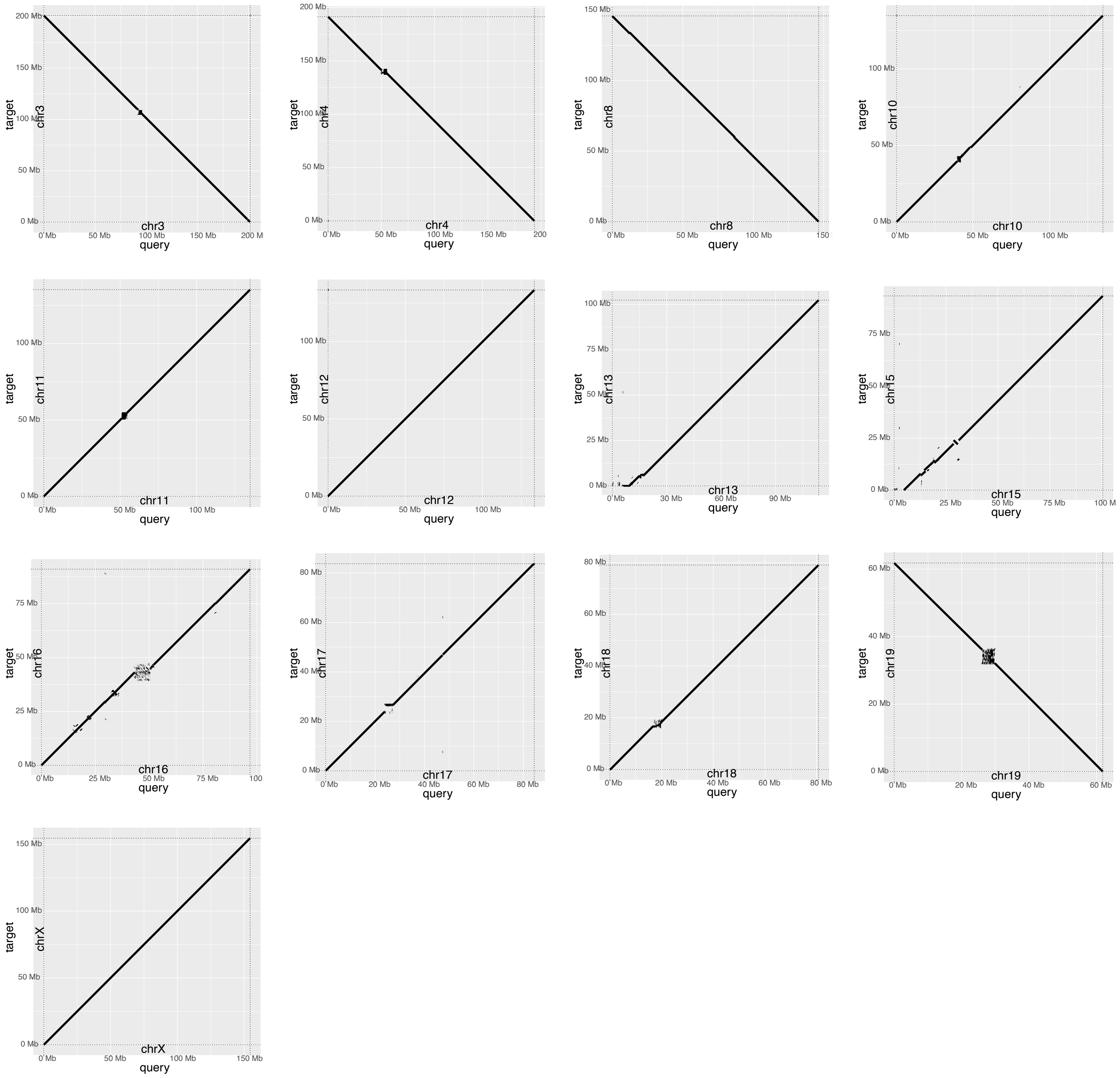


**Figure S19: Dot plots comparing GPatch NA12878 assembly with T2T-NA12878 after one round contig-breaking.** Dot plots compare patched chromosomes constructed with GPatch from HGSVC NA1878 maternal contigs, after one round of contig-breaking at the breakpoints of rearrangements >= 1MB in length, plotted against complete chromosomes from a recent chromosome-scale NA12878 genome assembly (T2T-NA12878). All Panels, target=T2T-NA12878, query=GPatch NA12878 after one round of contig breaking.

### Figure S20: Dot plots comparing GPatch NA12878 assembly with T2T-CHM13 after two rounds contig-breaking

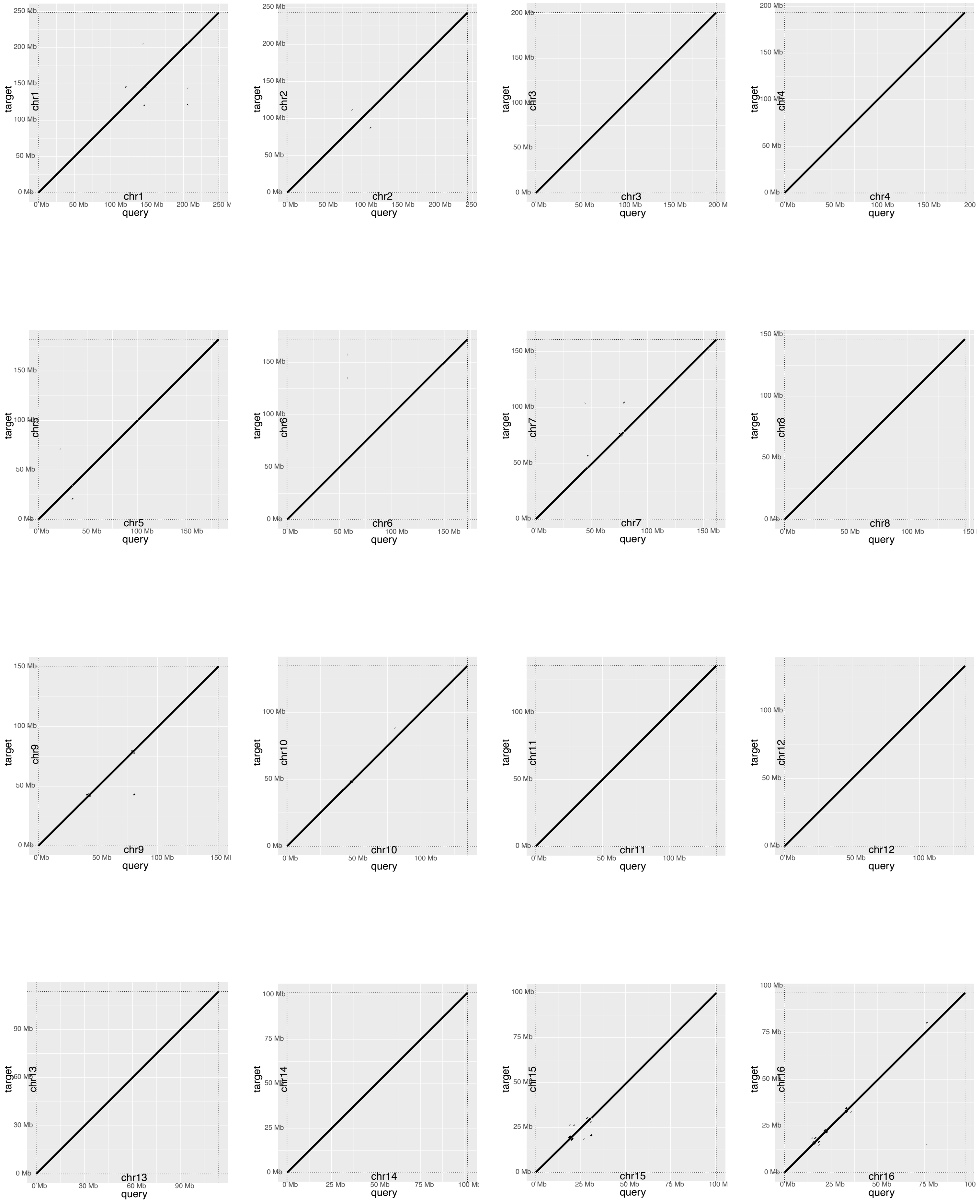


All Panels, target=T2T-CHM13, query=GPatch NA12878 after two rounds of contig breaking.

#### Figure S20: Dot plots comparing GPatch NA12878 assembly with T2T-CHM13 after two rounds contig-breaking

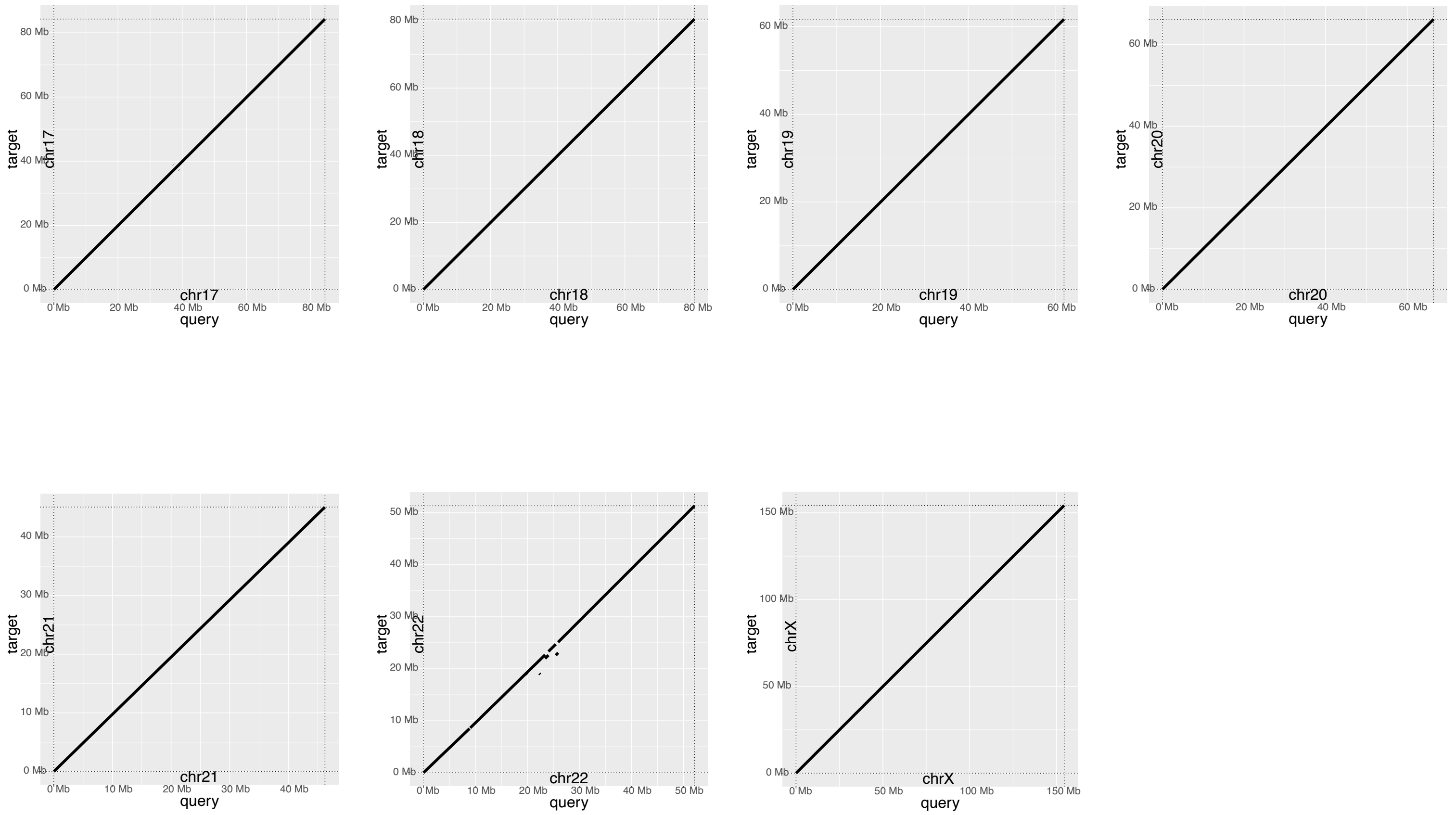


**Figure S20: Dot plots comparing GPatch NA12878 assembly with T2T-CHM13 after two rounds contig-breaking.** Dot plots compare patched chromosomes constructed with GPatch from NA1878 maternal contigs, after a second round of contig-breaking at the breakpoints of rearrangements >= 1MB in length, plotted against the T2T-CHM13 reference genome used for patching. All Panels, target=T2T-CHM13, query=GPatch NA12878 after two rounds of contig breaking.

**Figure S21: Dot plots comparing GPatch NA12878 assembly with T2T-NA12878 after two rounds contig-breaking**

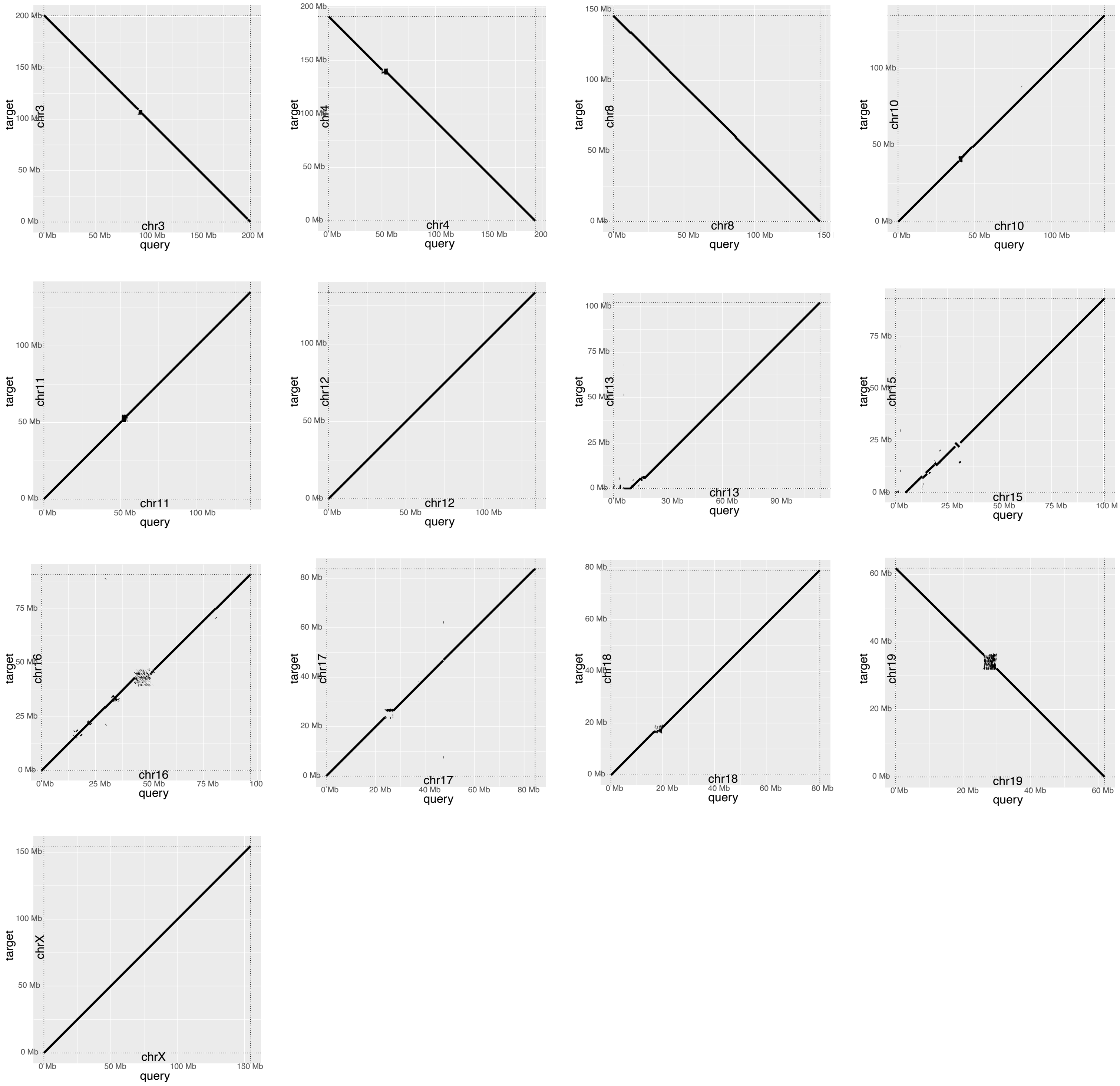


**Figure S21: Dot plots comparing GPatch NA12878 assembly with T2T-NA12878 after two rounds contig-breaking.** Dot plots compare patched chromo-somes constructed with GPatch from HGSVC NA1878 maternal contigs, after a second round of contig-breaking at the breakpoints of rearrangements >= 1MB in length, plotted against complete chromosomes from a recent chromosome-scale NA12878 genome assembly. All Panels, target=T2T-NA12878, query=GPatch NA12878 after two rounds of contig breaking.

#### Figure S22: Dot plots comparing GPatch HG002 assembly with T2T-CHM13 after one round contig-breaking

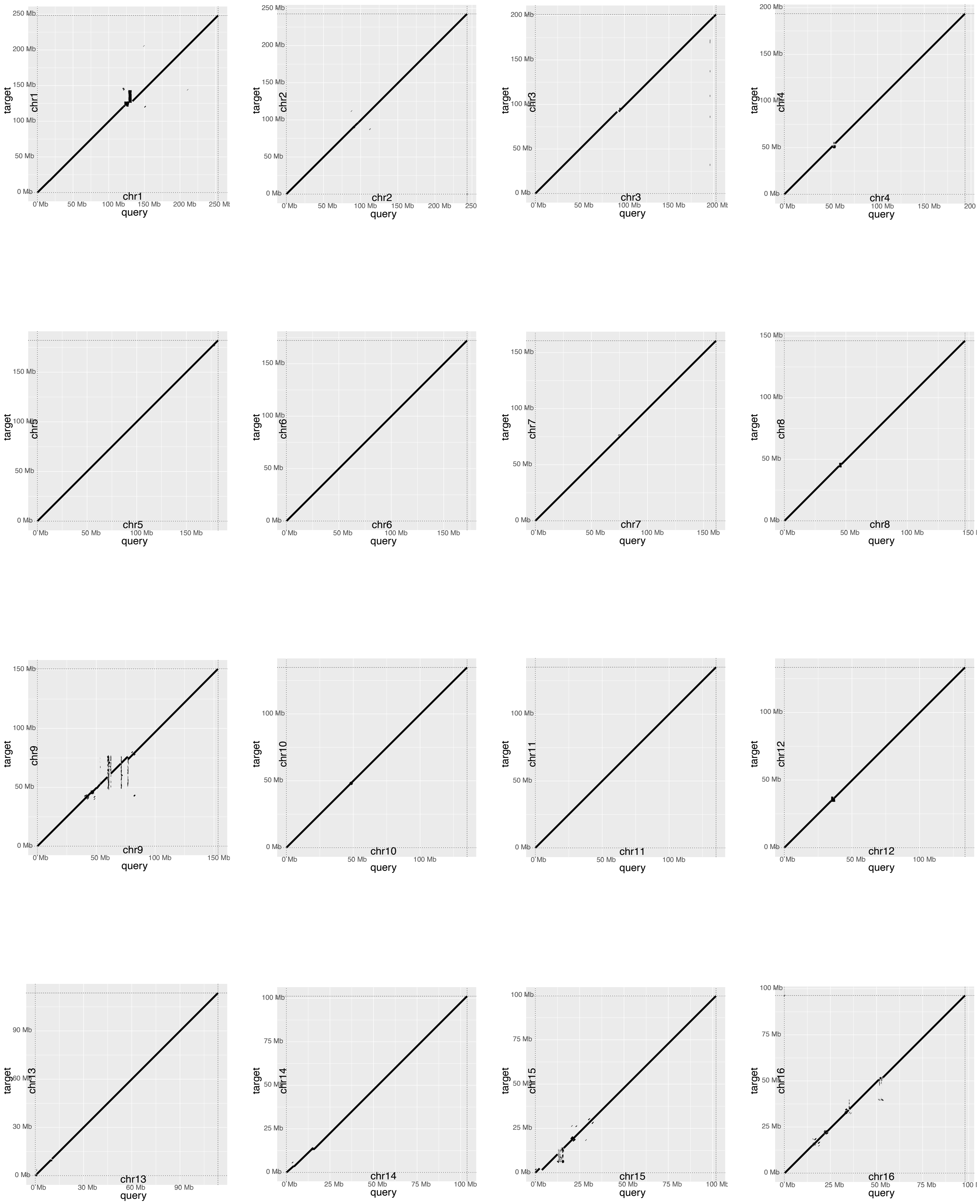


All Panels, target=T2T-CHM13, query=GPatch HG002 after one round of contig breaking.

**Figure S22: Dot plots comparing GPatch HG002 assembly with T2T-CHM13 after one round contig-breaking**

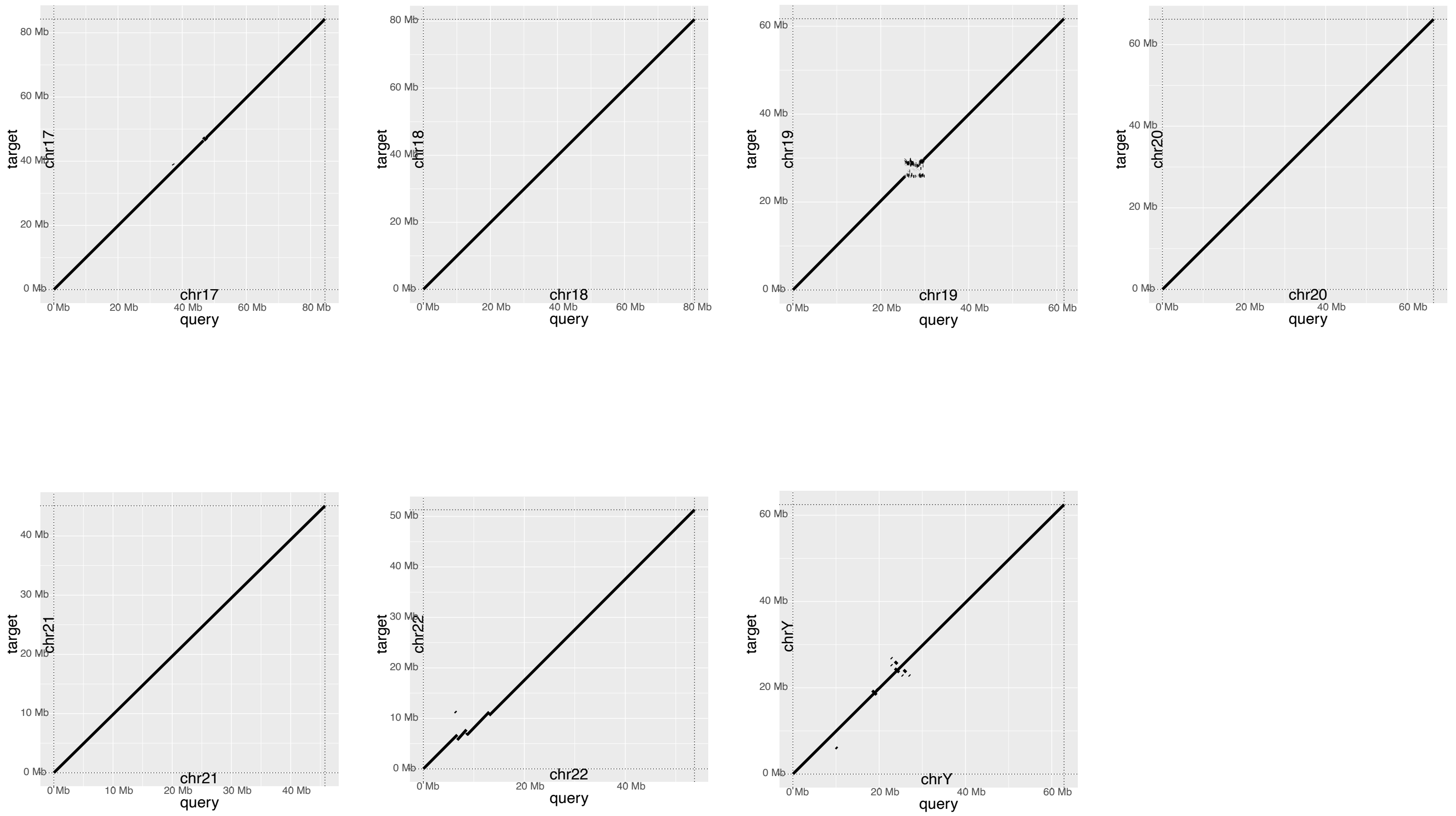


**Figure S22: Dot plots comparing GPatch HG002 assembly with T2T-CHM13 after one round contig-breaking.** Dot plots compare patched chromosomes constructed with GPatch from HPRC HG002 paternal contigs, after one round of contig-breaking at the breakpoints of rearrangements >= 1MB in length, plotted against the T2T-CHM13 genome assembly. All Panels, target=T2T-CHM13, query=GPatch HG002 after one round of contig breaking.

#### Figure S223 Dot plots comparing GPatch HG002 assembly with T2T-HG002 after one round contig-breaking

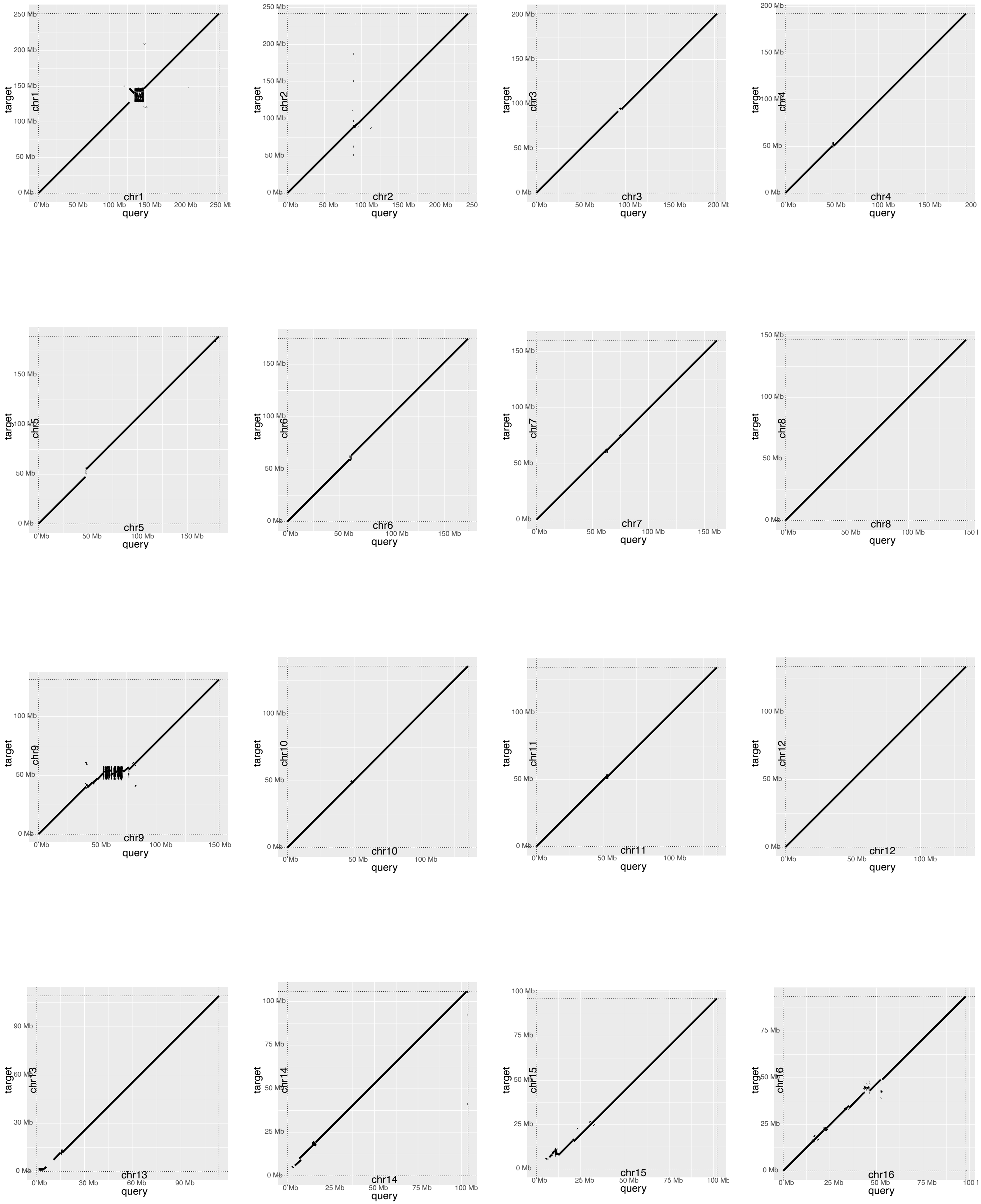


All Panels, target=T2T-HG002, query=GPatch HG002 after one round of contig breaking.

**Figure S23: Dot plots comparing GPatch HG002 assembly with T2T-HG002 after one round contig-breaking**

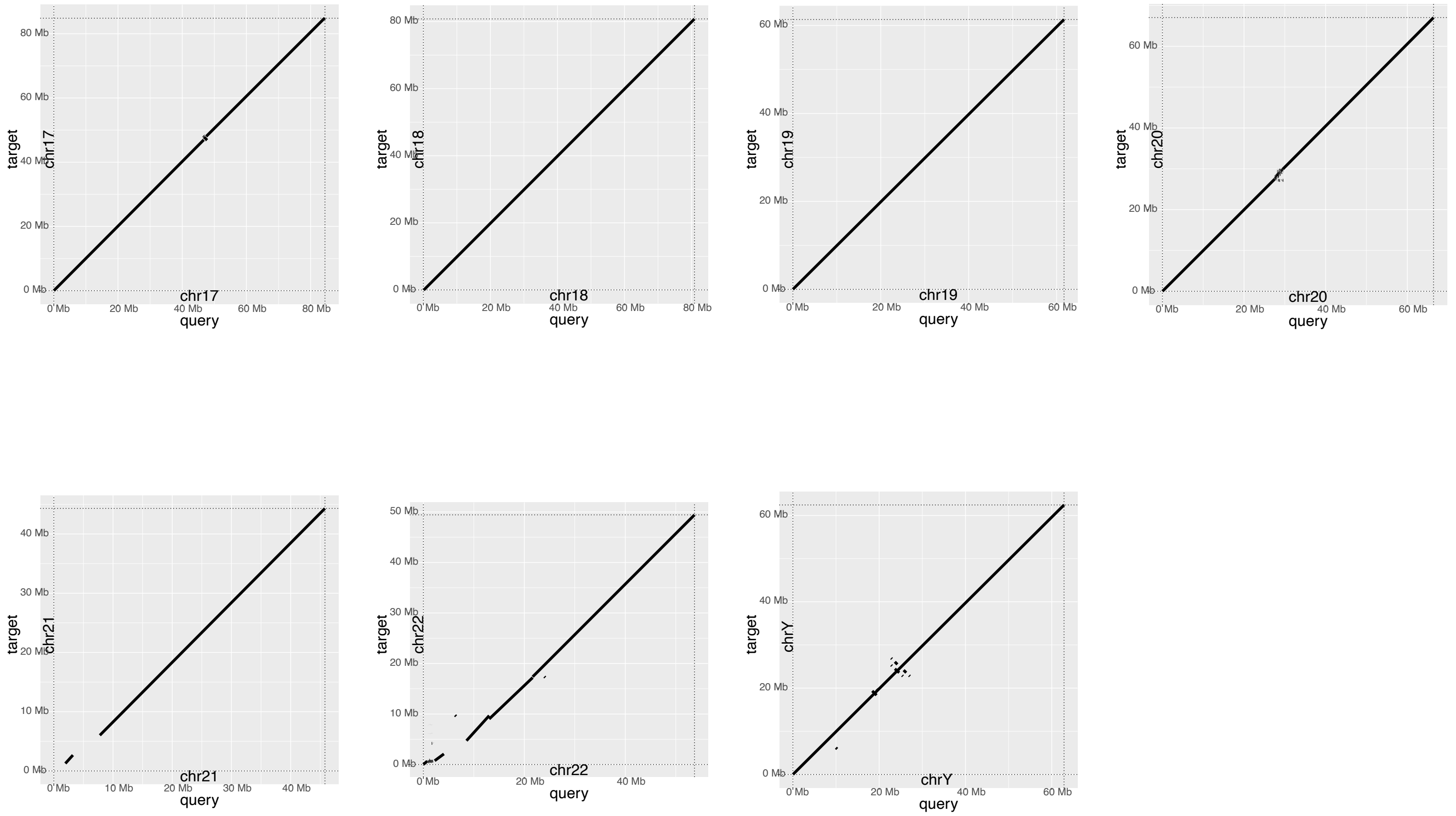


**Figure S23: Dot plots comparing GPatch HG002 assembly with T2T-HG002 after one round contig-breaking.** Dot plots compare patched chromosomes constructed with GPatch from HPRC HG002 paternal contigs, after one round of contig-breaking at the breakpoints of rearrangements >= 1MB in length, plotted against the T2T-HG002 genome assembly. All Panels, target=T2T-HG002, query=GPatch HG002 after one round of contig breaking.

**Figure S24: NA12878 Hi-C Data Processed with the GPatch NA12878 Assembly as Reference**

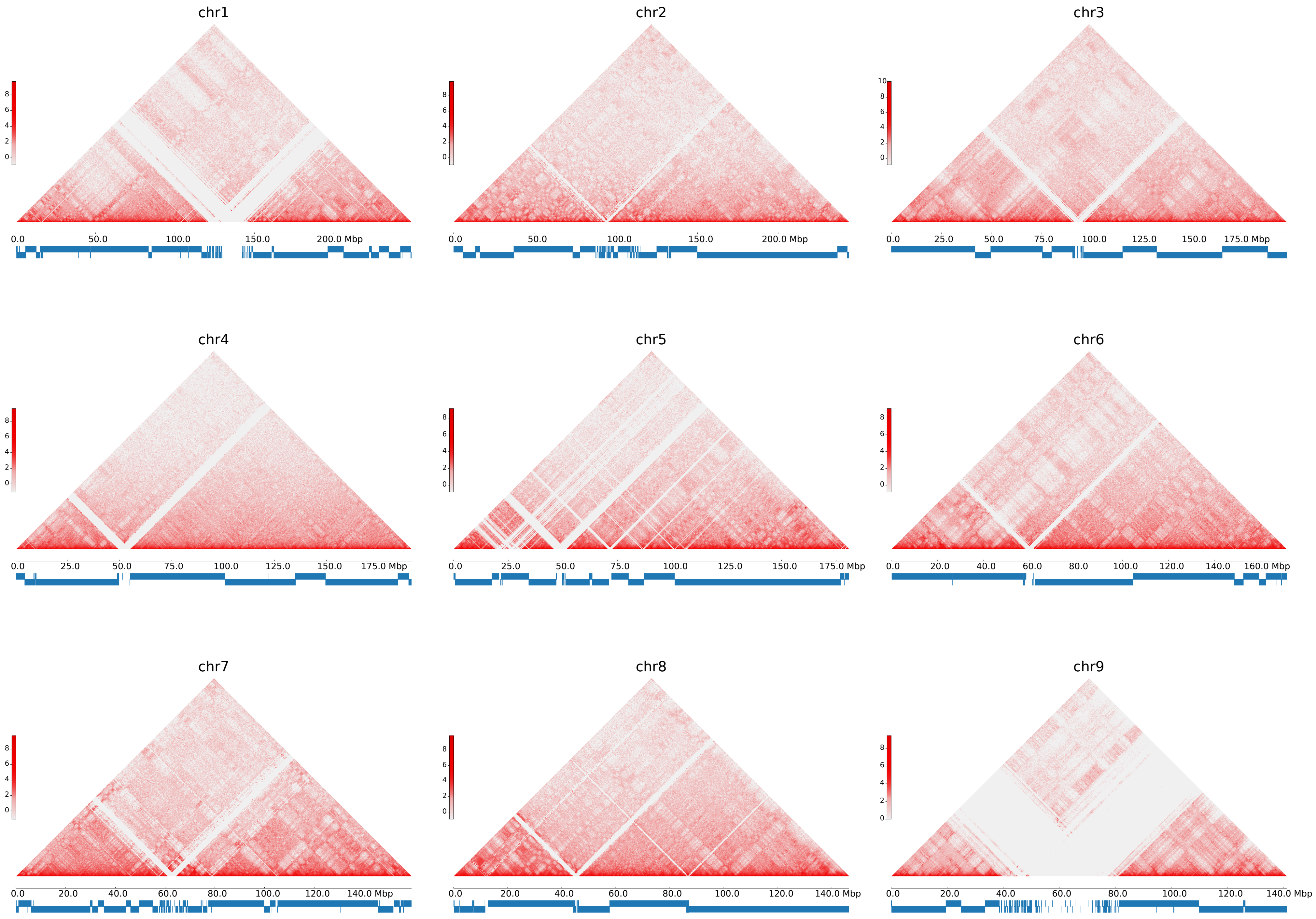

**Figure S24: NA12878 Hi-C Data Processed with the GPatch NA12878 Assembly as Reference**

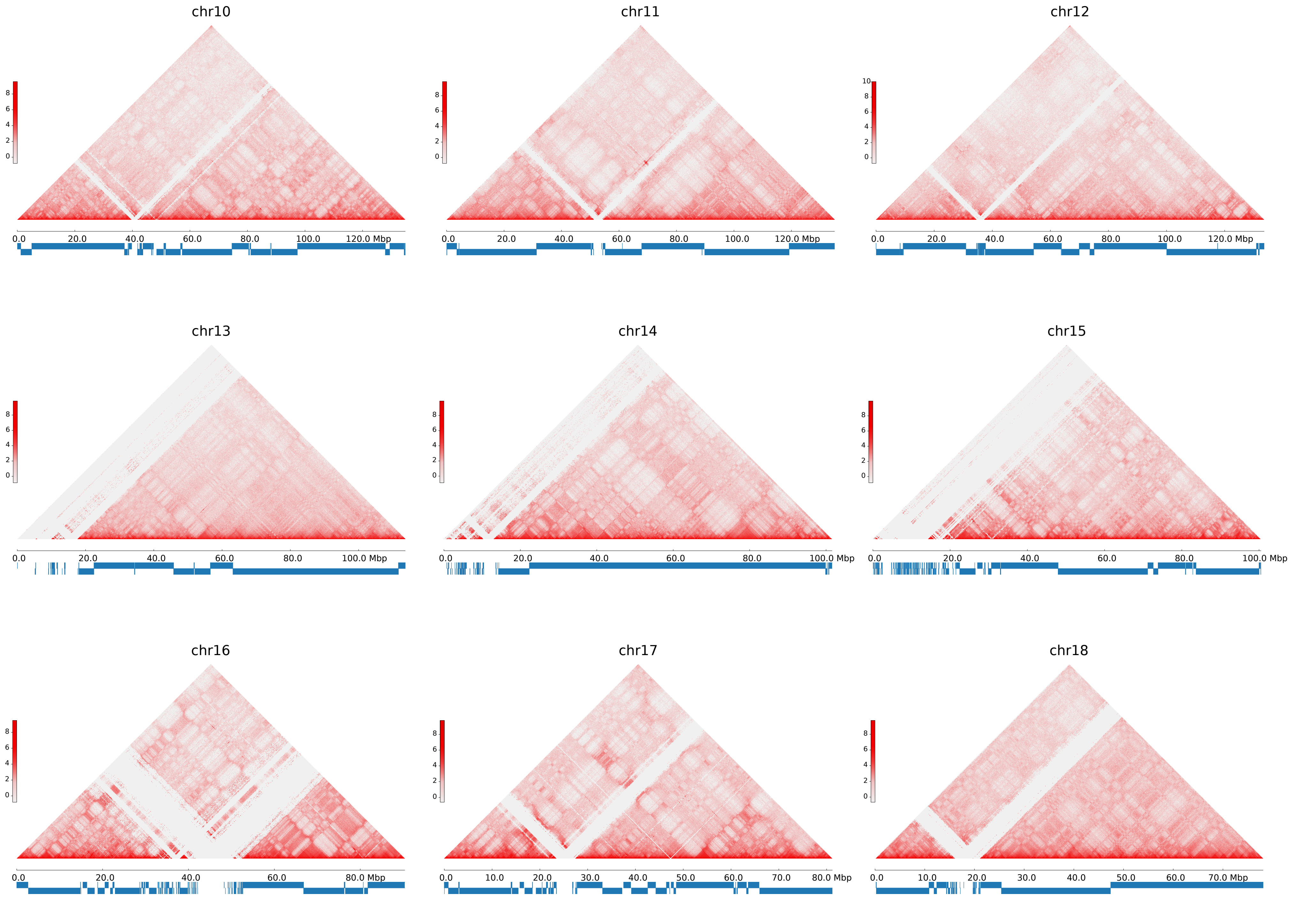

#### Figure S24: NA12878 Hi-C Data Processed with the GPatch NA12878 Assembly as Reference

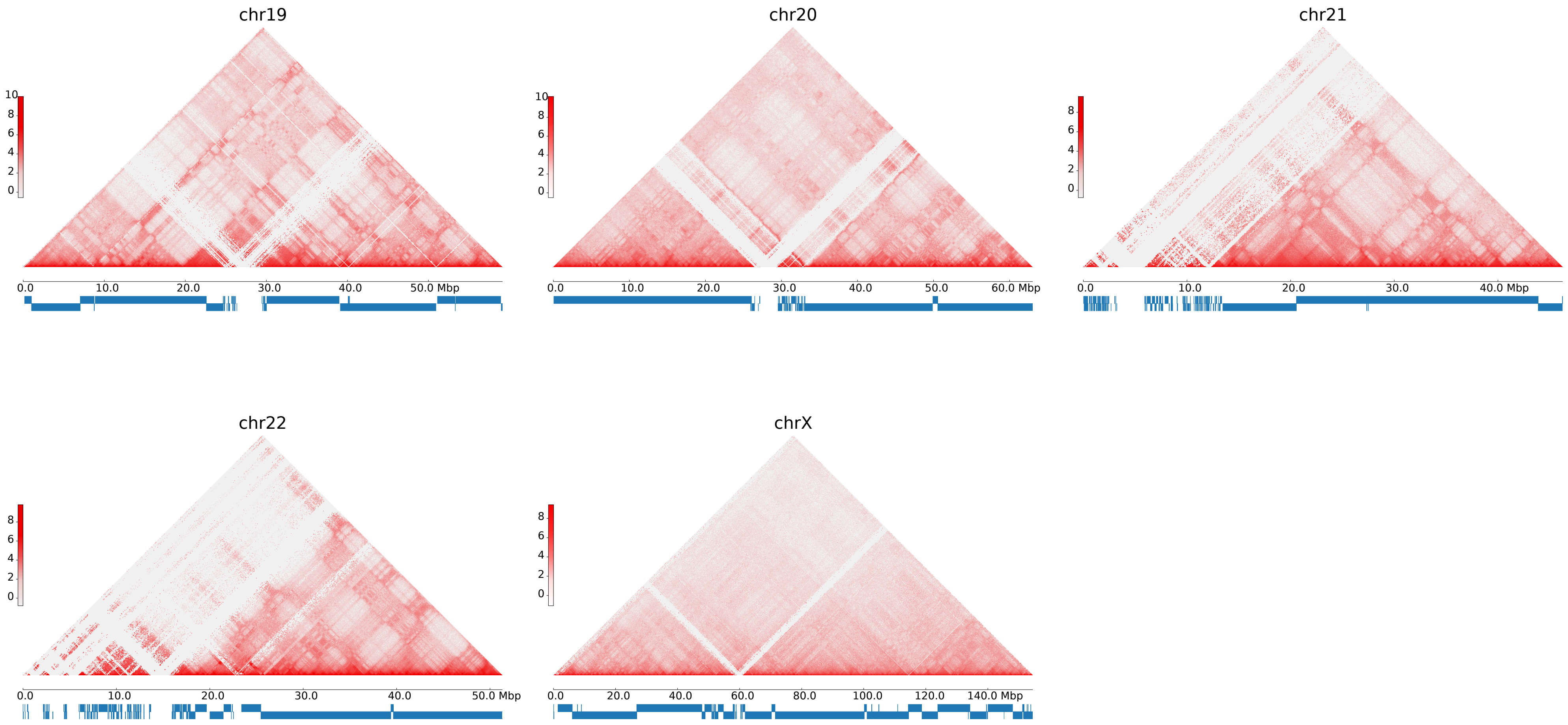


**Figure S24: NA12878 Hi-C Data Processed with the GPatch NA12878 Assembly as Reference.** All panels: Hi-C contact matrices plotted at 80kb resolution with balanced normalization. Blue bars below the X-Axis indicate the positions of contigs in the patched chromosome sequence, with intervening white regions indicating patches.

**Figure S25: NA12878 Hi-C Data Processed with the T2T-CHM13 Assembly as Reference**

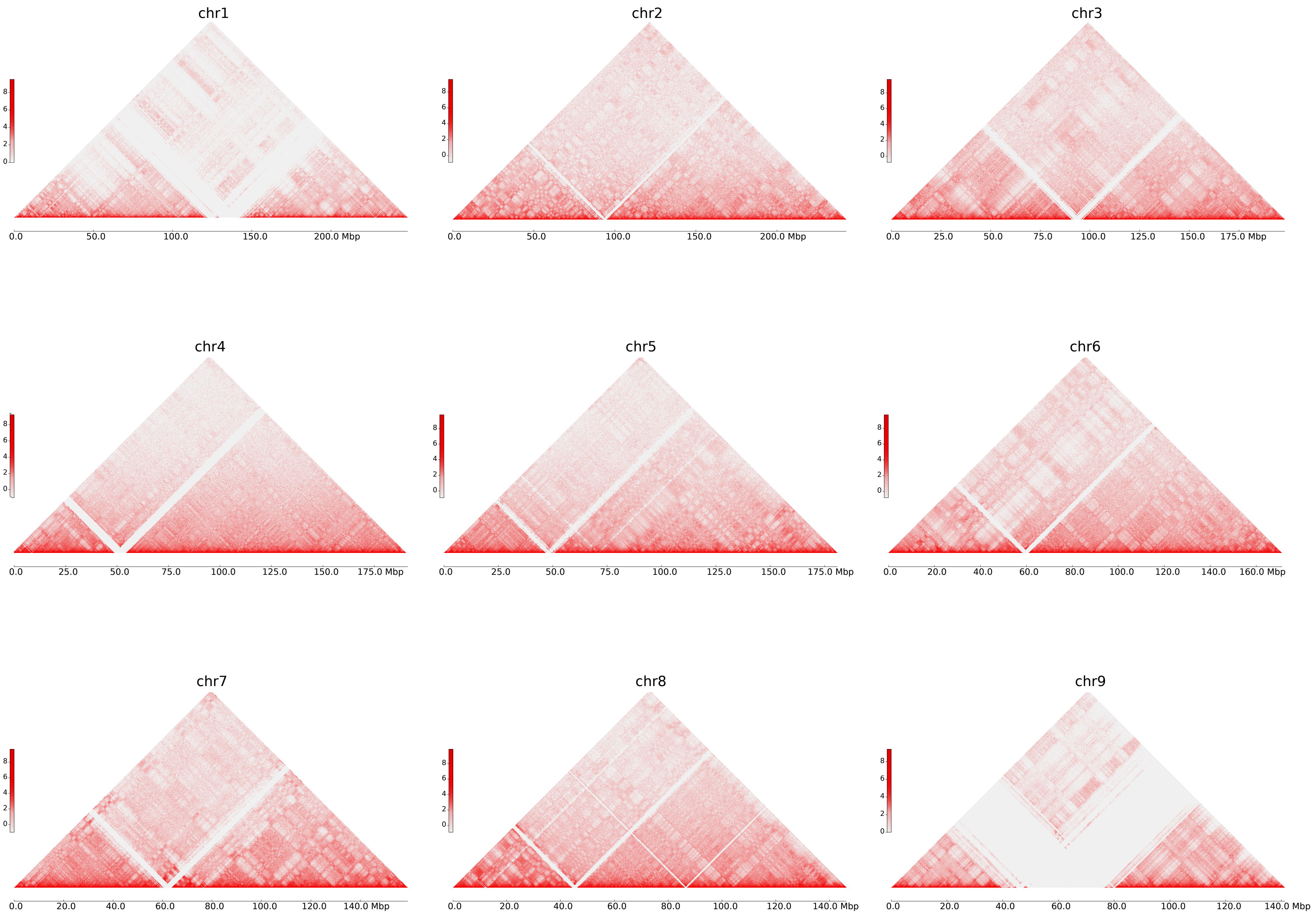

### Figure S25: NA12878 Hi-C Data Processed with the T2T-CHM13 Assembly as Reference

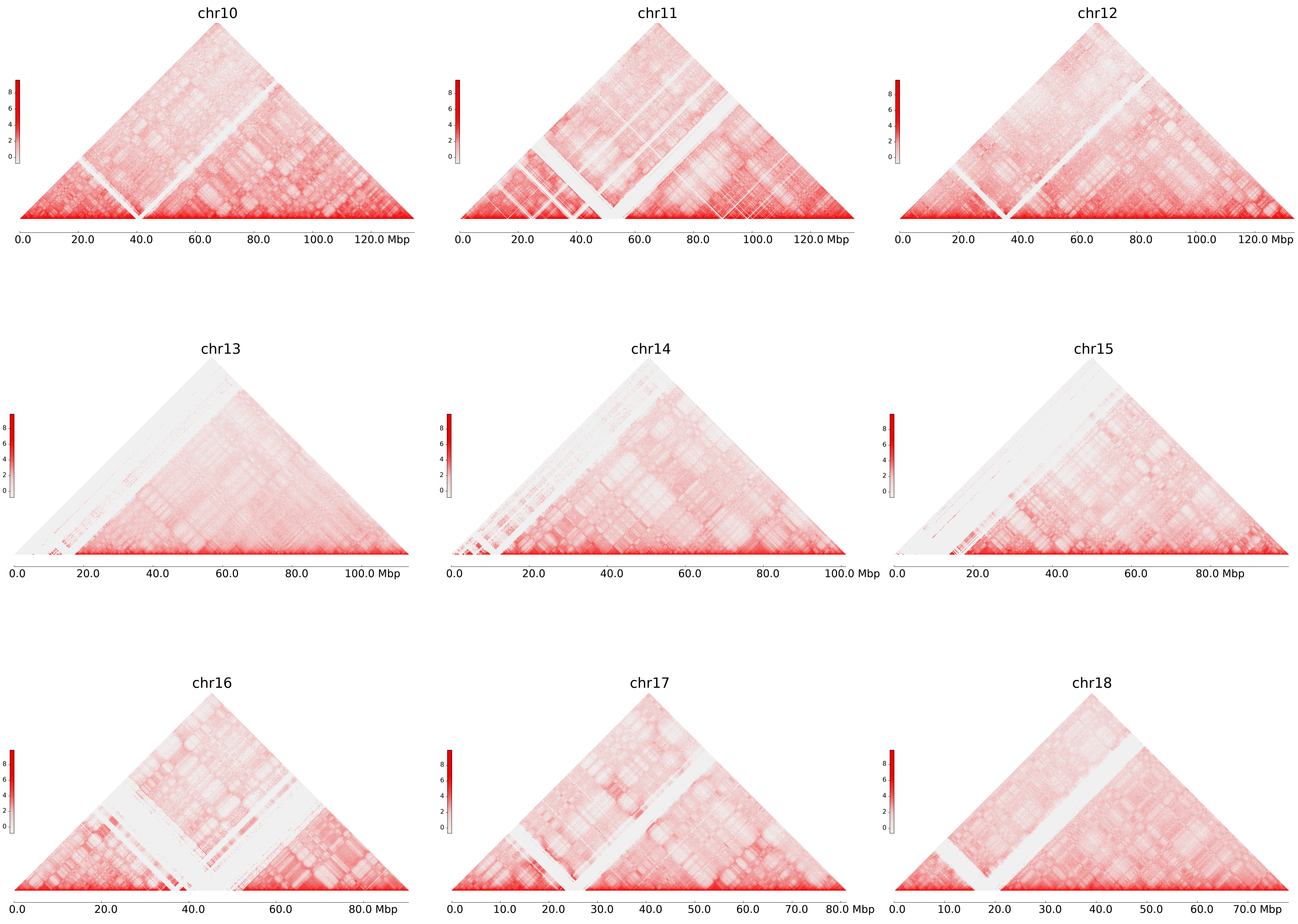

#### Figure S25: NA12878 Hi-C Data Processed with the T2T-CHM13 Assembly as Reference

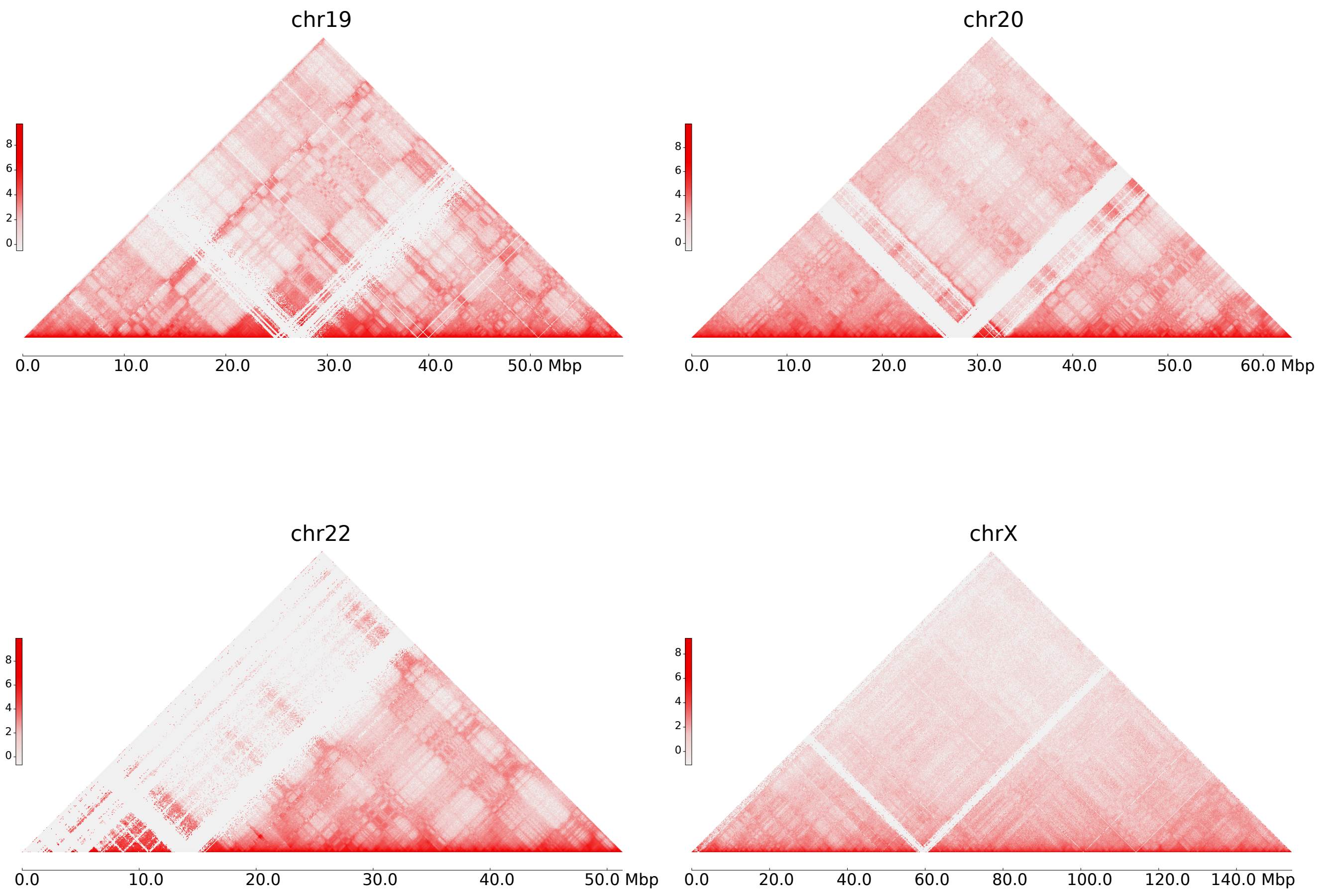


**Figure S25: NA12878 Hi-C Data Processed with the T2T-CHM13 Assembly as Reference.** All panels: Hi-C contact matrices plotted at 80kb resolution with balanced normalization.
